# The Genome of the Zebra Mussel, *Dreissena polymorpha*: A Resource for Invasive Species Research

**DOI:** 10.1101/696732

**Authors:** Michael A. McCartney, Benjamin Auch, Thomas Kono, Sophie Mallez, Ying Zhang, Angelico Obille, Aaron Becker, Juan E. Abrahante, John Garbe, Jonathan P. Badalamenti, Adam Herman, Hayley Mangelson, Ivan Liachko, Shawn Sullivan, Eli D. Sone, Sergey Koren, Kevin A. T. Silverstein, Kenneth B. Beckman, Daryl M. Gohl

## Abstract

The zebra mussel, *Dreissena polymorpha*, continues to spread from its native range in Eurasia to Europe and North America, causing billions of dollars in damage and dramatically altering invaded aquatic ecosystems. Despite these impacts, there are few genomic resources for *Dreissena* or related bivalves, with nearly 450 million years of divergence between zebra mussels and its closest sequenced relative. Although the *D. polymorpha* genome is highly repetitive, we have used a combination of long-read sequencing and Hi-C-based scaffolding to generate the highest quality molluscan assembly to date. Through comparative analysis and transcriptomics experiments we have gained insights into processes that likely control the invasive success of zebra mussels, including shell formation, synthesis of byssal threads, and thermal tolerance. We identified multiple intact Steamer-Like Elements, a retrotransposon that has been linked to transmissible cancer in marine clams. We also found that *D. polymorpha* have an unusual 67 kb mitochondrial genome containing numerous tandem repeats, making it the largest observed in Eumetazoa. Together these findings create a rich resource for invasive species research and control efforts.

## Introduction

Native to a small region of southern Russia and Ukraine ^1^ zebra mussels (*Dreissena polymorpha*, Fig. 1a) have spread throughout European ^2,3^ and North American ^4^ fresh waters to become one of the world’s most prevalent and damaging aquatic invasive species ^5^. Fouling of water intake pipes cost the power generation industry over $3 billion USD from 1993-1999 in the Laurentian Great Lakes region alone ^6^, where *Dreissena* cause extensive damage to hydropower, recreation and tourism industries, and lakefront property ^7,8^. Dense infestations smother and outcompete native benthic species and remove large amounts of phytoplankton from lakes and rivers, causing population declines and extinctions of native freshwater mussels and other invertebrates, damage to fish populations ^9–14^, and dramatic restructuring of aquatic food webs ^15–17^. The congener *D. rostriformis* (the quagga mussel), while far less widespread than zebra mussels in inland waters, has ecologically replaced zebra mussels in much of the Laurentian Great Lakes proper and in deep European lakes, and may cause even greater ecological damage in those systems ^18–20^.

**Figure 1.**
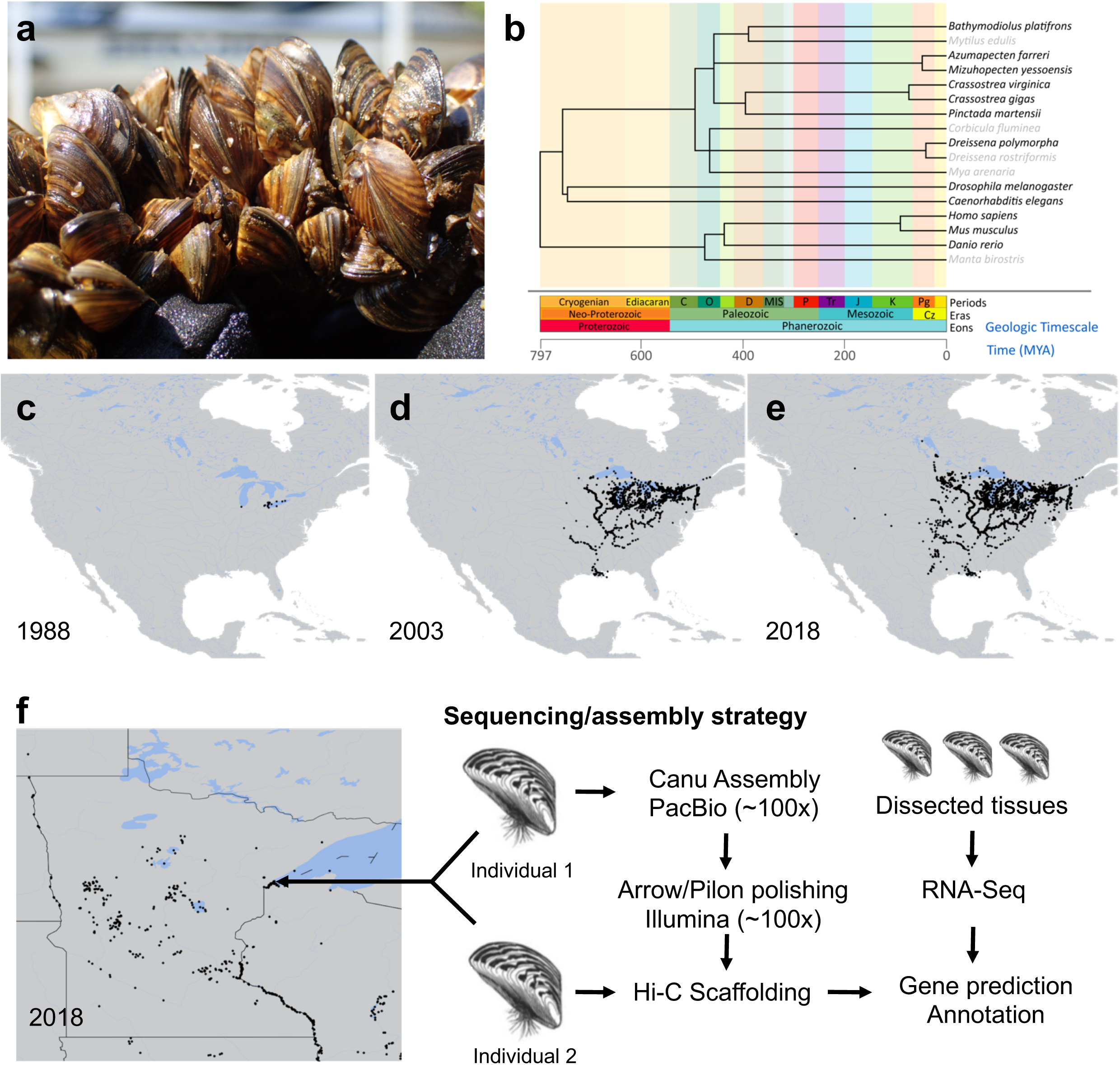
Zebra mussel biogeography and genome sequencing strategy. a) Photo of *D. polymorpha* (by N. Blinick). b) Phylogenetic tree showing the evolutionary divergence between *D. polymorpha* and other sequenced bivalve genomes. For context, the evolutionary divergence of humans, mice, zebrafish, manta rays, nematodes, and fruit flies are shown. Bolded text indicates that a genome sequence for that organism is publicly available. Divergence times and tree construction based on Kumar *et al.* ^135^ c-e) Maps depicting the spread of *D. polymorpha* in the United States of America from 1988 through 2018. Data from US Geological Survey, NAS database^136^. f) Map showing the extent of zebra mussel infestation in Minnesota lakes as of 2018 and depicting the location where the specimens for genome sequencing and scaffolding were collected (left). Summary of the sequencing and annotation strategy (right).

The ongoing European and North American invasions (Fig. 1c-e) have spurred an explosion in research effort on *Dreissena*, particularly focused on physiology, autecology, and ecosystem impacts ^21^. Aside from molecular systematic and population genetic studies ^1,22–25^, comparatively little genetic work has been accomplished, with transcriptomes from a few tissues ^26,27^ being the only genomic resources.

Bivalves are a diverse Class of Mollusca with over 10,000 described species in marine and freshwater environments ^28,29^. To date, complete genomes have been sequenced and analyzed in only eight species—most of them marine organisms of commercial value (Fig. 1b, Supplemental Table 1). Yet 21 invasive bivalve species cause damage to aquatic and marine ecosystems worldwide ^30^ and only the golden mussel, *Limnoperna fortunei* ^31^ has a published genome available (the quagga mussel is also being sequenced at present ^32^). Moreover, the divergence time between *Dreissena* and other bivalve species with published genomes is estimated at more than 400 million years ago (Fig. 1b). Sequencing of the zebra mussel genome will provide a resource for comparative genomic and other studies of an underexplored lineage of bivalves that includes two of the world’s most notorious and damaging invasive species ^20,33^.

Here we present the genome sequence of *D. polymorpha*. Using short and long-read sequencing technologies as well as Hi-C based scaffolding, we generated a chromosome-scale genome assembly with high contiguity and completeness. Through comparative analysis and RNA-sequencing experiments, we provide insights into the process of shell formation, the formation of byssal thread attachment fibers, and mechanisms of thermal tolerance—three processes of critical importance to continued spread. The genomic resources we describe lay the groundwork for further investigation of the traits that allow zebra mussels to thrive as an invasive species and are a step towards developing control strategies for this economically and ecologically damaging aquatic invader.

## Results

To sequence the *D. polymorpha* genome, we used the strategy outlined in Fig. 1f. We generated a size-selected PacBio library with *≥*20 kb inserts (Supplemental Fig. 1–2). Using the PacBio Sequel Single Molecule Real Time (SMRT) sequencing platform, we generated 168.97 gigabases (Gb) of sequencing data for an estimated coverage over 100x, assuming a genome size (from densitometry measures of DNA content in stained nuclei) of 1.66 Gb ^34^. The subread N50 for the PacBio reads was 16,524 bp, validating the high quality of the input DNA and PacBio sequencing library.

Canu ^35^ yielded a 2.92 Gb assembly, with 15,311 contigs and a contig N50 of 549,263 bp. The assembly was 1.3 Gb larger than previously estimated ^34^ due to the relatively high heterozygosity of the sample (2.13% estimated from GenomeScope and previous Illumina sequencing). Identification of allelic contigs ^36^ removed redundancy and yielded a 1.8 Gb assembly containing 2,863 contigs with a contig N50 value of 1,111,027 bp (Table 1). Hi-C ^37^ analysis of the polished assembly generated 16 scaffolds spanning 98.4% of the assembled genome (128 unscaffolded contigs comprised the remaining assembled material, Supplemental Fig. 3). Earlier cytogenetic work found 1N=16 chromosomes for *D. polymorpha* ^38,39^. The scaffold N50 value was >107 Mb and the scaffold L50 value was 7, consistent with a chromosome-scale assembly. The resulting scaffolds and contigs were checked for contamination from bacterial genomic DNA and sequencing adapters, and a single contig was removed because it mapped to the PacBio sequencing control.

**Table 1.**
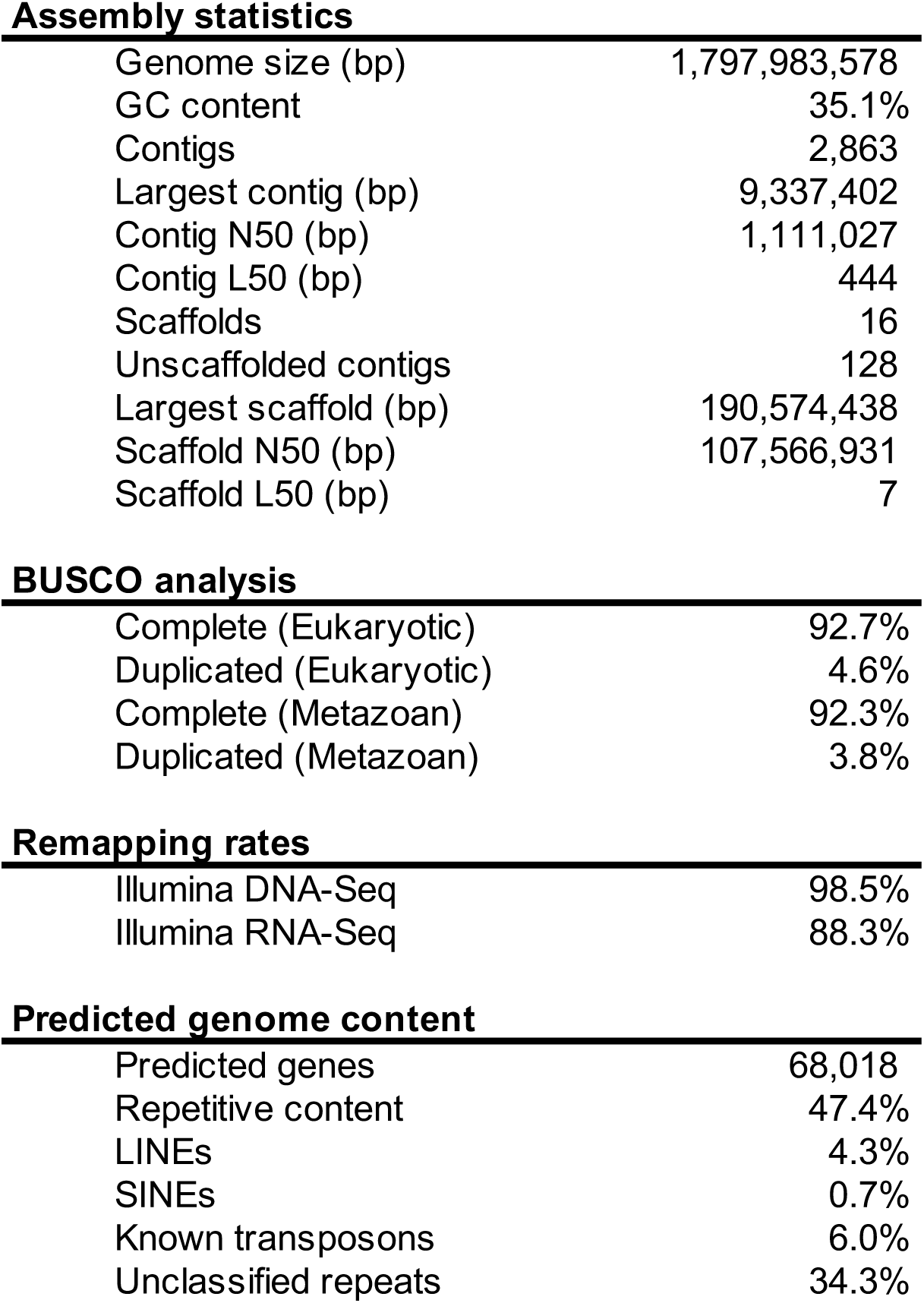
Genome assembly statistics. Statistics summarizing the contiguity, completeness, and content of the *D. polymorpha* genome.

BUSCO analysis ^40^ demonstrated that in addition to having high contiguity, the *D. polymorpha* genome assembly is highly complete, with >92% of eukaryotic and metazoan BUSCOs identified and <5% duplication (Table 1). Also consistent with high completeness, 98.5% of the Illumina DNA sequencing reads mapped to the *D. polymorpha* assembly (Table 1, Supplemental Table 2).

### Features of the *D. polymorpha* genome

The genome assembly was annotated using *de novo* as well as protein and transcript-guided methods. This analysis resulted in a list of 68,018 genes. Functional annotation was carried out by mapping to a number of databases, including PFAM ^41^, InterPro ^42^, UniProtKB ^43^, Merops ^44^, and CAZymes ^45^. Due to the large evolutionary divergence between *D. polymorpha* and other sequenced genomes, the majority of the predicted genes had no annotations assigned. However, 12,772 genes had recognizable orthologs.

Repetitive DNA is abundant in bivalve genomes ^46–49^, which makes assembly challenging. The *D. polymorpha* genome is also highly repetitive (47.4% repetitive content, Fig. 2a, Table 1) and AT-rich (35.1% GC). While a portion of this repetitive content could be assigned to long or short interspersed elements (LINEs or SINEs), or to known transposons. The majority of the repeats, or 34.3% of the genome, could not be classified (Table 1).

**Figure 2.**
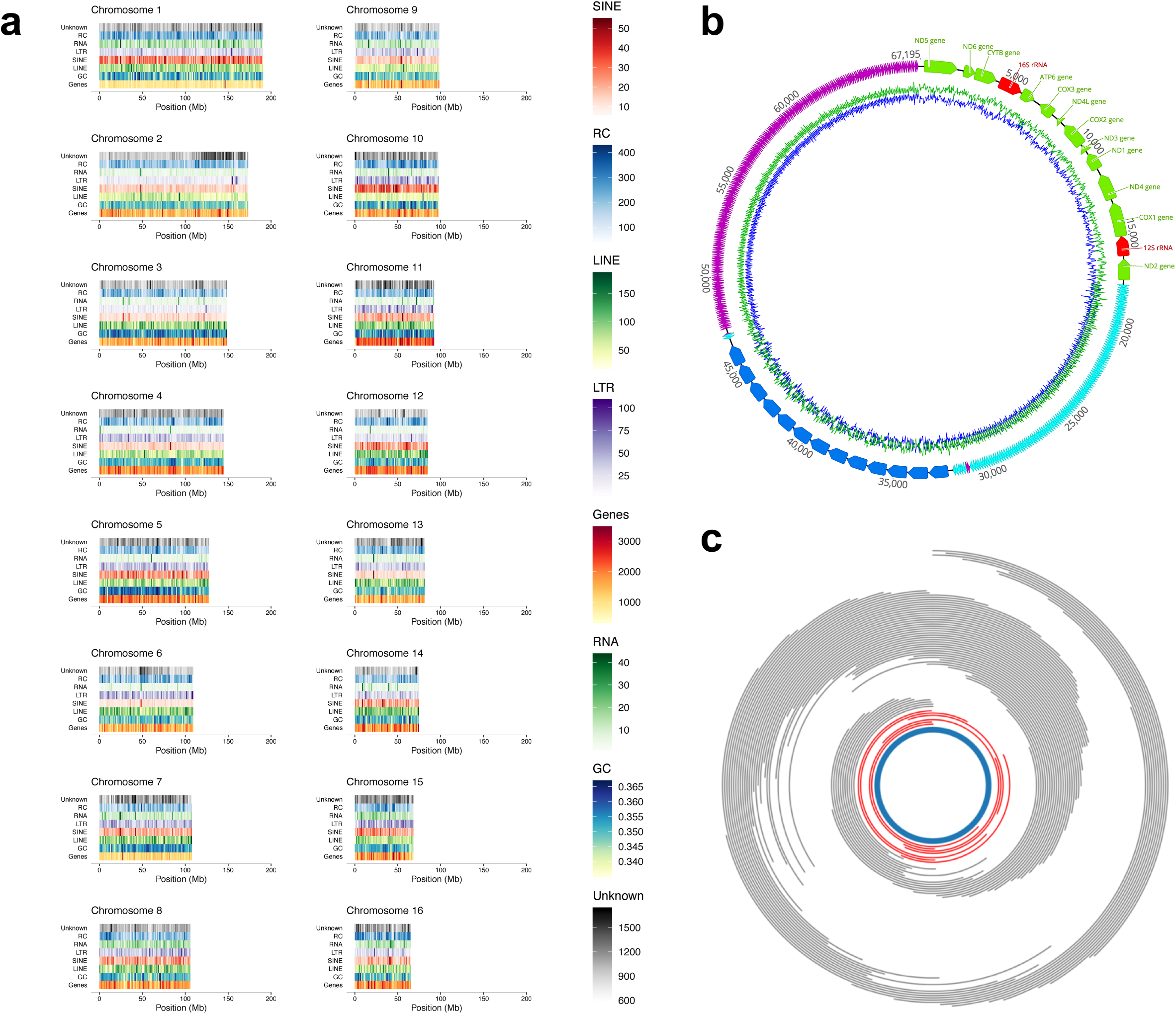
*D. polymorpha* genome and mitogenome structure and content. a) Plots depicting the gene content, repeat and transposon density, and GC content of the 16 *D. polymorpha* chromosomal scaffolds. b) Proposed circular mitochondrial genome structure. GC content plots (blue) based on 40 bp sliding window. Annotations based on sequence similarity to previously published partial mitochondrial genome ^26^. Coding regions are in green and red, and the three large repeat blocks are colored turquoise, blue, and purple. c) Plot of long (>25 kb) Oxford Nanopore (red) and PacBio (grey) reads supporting the proposed 67 kb circular mitogenome structure. Orientation of mitochondrial genome (blue) is the same as in panel b.

The zebra mussel genome contains several notable gene family expansions (Supplemental Fig. 4, Supplemental Files 1-2). Relative to humans, *D. polymorpha* shows expansions of genes related to cellular stress responses and apoptosis that in several cases even surpass those in Pacific oyster (*Crassostrea gigas*) ^49^. Expansions of gene families encoding Hsp70s (heat shock chaperones), caspases (apoptosis), and IAPs (Inhibitor of Apoptosis Proteins) exceeds Pacific oyster; expansion of Cu-Zn superoxide dismutases (antioxidant defense) and C1q domain-containing proteins (innate immunity) show expansions that are, respectively, equal to and smaller than *Crassostrea gigas*, while cytochrome P450s (xeniobiotic detoxification) are contracted relative to humans (Table 2).

**Table 2.**
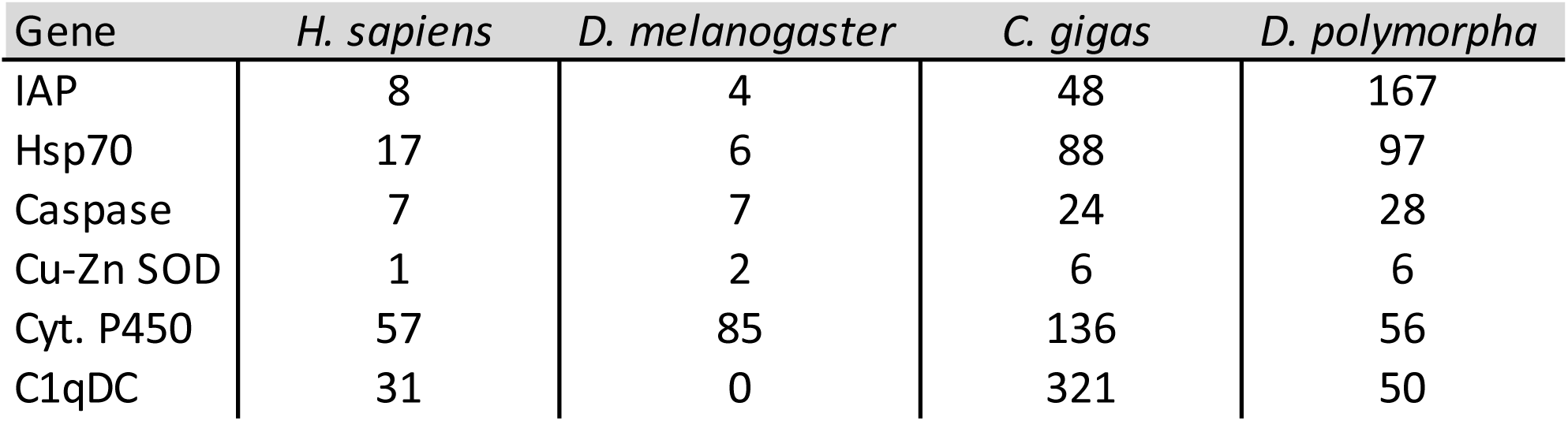
D. polmorpha gene family expansions. Selected gene family expansion data comparing *D. polymorpha* to *C. gigas*, *D. melanogaster*, and *H. sapiens*. Data for *C. gigas*, *D. melanogaster*, and *H. sapiens* from Zhang *et al.* ^49^.

Examination of orthology to eastern oyster (*Crassostrea virginica*) identified 10,065 orthologous groups (Supplemental Files 3-4). A total of 26.3% of zebra mussel genes that were used for orthologous group identification were assigned to a group within *C. virginica*. This is consistent with the low sequence similarity between zebra mussel and *C. virginica*, even at the amino acid level. A majority (5,753; 57.16%) of the orthologous groups involved equal numbers of genes from zebra mussel and *C. virginica*. Of orthologous groups of unequal size, there were far more groups with contracted than expanded gene families in zebra mussel, relative to this distantly related bivalve (76.86% contracted and 23.14% expanded).

In the initial assembly we recovered a single contig containing the *D. polymorpha* mitochondrial genome (Figure 2b). A partial *D. polymorpha* mitogenome sequence was previously published ^26^, but contained a gap which short-read sequencing and targeted PCR were unable to resolve. PacBio and Oxford Nanopore sequencing (Fig. 2c) reveals that this “gap” is a large highly repetitive segment of nearly 50kb, making the *D. polymorpha* mitogenome the largest reported so far from Eumetazoa at 67,195 bp. The repetitive segment consists of three distinct blocks of direct tandem repeats (Supplemental Fig. 5), with individual repeat elements of approximately 125 bp, 1030 bp, and 86 bp, each copied many times. The 86bp repeat element was discovered only after re-mapping of long reads to the initial assembly, which indicated an area of especially high coverage and read-clipping (Supplemental Fig. 6). An alternate mitochondrial assembly generated using FALCON revealed this anomaly to be an additional repeat sequence, to which the PacBio and Oxford Nanopore reads mapped seamlessly. Thus, the FALCON mitogenome assembly has been used in datasets associated with this paper (Fig. 2b-c). We further validated that the mitochondrial contig was not associated with chromosomal sequences by examining Hi-C data, where the association between the mitochondrial contig and the *D. polymorpha* chromosomes was much lower than the association between contigs on the same scaffold, and was comparable to background levels of crosslinking seen between contigs on different scaffolds (Supplemental Fig. 5). Eumetazoan mitogenomes, with few exceptions, generally lack length variation and non-coding DNA content ^26^. Among these few exceptions are the long enigmatic mitogenomes of scallops^50^, but unlike scallops the coding genes of *D. polymorpha* remain contiguous, instead of being interrupted by interspersed repeats. Typical of animals, the coding region in D. polymorpha is compact (~17.5 kb), but as is common in bivalves ^50^, the order of mitochondrial genes is unique to the species. The reason for this unusual mitochondrial DNA (mtDNA) structure is unknown, but similar repetitive sequences have been observed in the mtDNA of plants and it has been suggested that such repeats may result from increased double-stranded break repair due to the need to cope with desiccation-related DNA damage ^51^.

Some mussels exhibit doubly uniparental inheritance (DUI) of mtDNA, or transmission of two gender-associated mitogenomes: an F-type through eggs and M-type through sperm ^52,53^. DUI is present in *Venerupis;* i.e, in Superorder Imparidentia, containing Dreissenidae. We found no evidence for a second divergent mitogenome. We located no other contigs (via tblastx) that contain mitochondrial genes. Further, re-mapping of high-accuracy Illumina reads from the same mussel to the mitochondrial genome revealed no SNPs within the coding region (Supplemental Fig. 6), indicative of homoplasmy. The tissues used for DNA extraction included ripe male gonad with abundant motile sperm. With DUI, extracts would be expected to contain both mtDNAs, as the M-type is transmitted exclusively through male germline, while in somatic tissues the F-type is predominant ^53,54^.

### Steamer-Like Elements

We identified a number of Long Terminal Repeat (LTR) retrotransposons that are similar in structure to *Steamer*, a transposable element (TE) that in the soft-shelled clam *Mya arenaria* causes a leukemia that is transmissible between conspecifics ^55,56^. A high incidence of horizontal transmission of TEs (HTT) has spread these Steamer-Like Elements (SLEs) across several bivalves that also contract transmissible cancers, and across phyla to several marine animal species that do not ^57^. We identified eight copies of putative SLEs in the *D. polymorpha* genome with intact polycistronic open reading frames (ORFs) that span the conserved Gag-Pol polyprotein and are flanked by LTRs (Fig. 3a). The *D. polymorpha* elements were aligned to the full length ORFs of 99 *Ty3/Gypsy* LTR-retrotransposons. Phylogenetic analysis confirmed that the TEs in *D. polymorpha* are *Steamer*-like elements (SLEs: Supplemental Fig. 7). The *D. polymorpha* elements grouped within the Mag C clade with 100% bootstrap support, and sister to *Steamer*. Next, we performed phylogenetic analysis of the *D. polymorpha* elements and amplicons from within the RNaseH-integrase domain of Gag-Pol from 47 other bivalve species, characterized in an earlier study of HTT events^57^. Our phylogenetic analysis identified a minimum of three HTTs leading to their spread to zebra mussels from marine bivalves (Fig. 3b), including an event that is additional to and independent of the two HTTs identified previously ^57^. It is unknown whether SLEs are currently undergoing active transposition within zebra mussels. However, the high levels of sequence similarity between Gag-pol regions of different SLE loci, and between the two LTRs of each SLE, indicates that the latest wave of transposition in this genome was recent. We also identified numerous degenerate copies that are missing portions of *Gag-Pol* or LTR sequences, as well as isolated LTR scars on most chromosomes (Supplemental Fig. 8).

**Figure 3.**
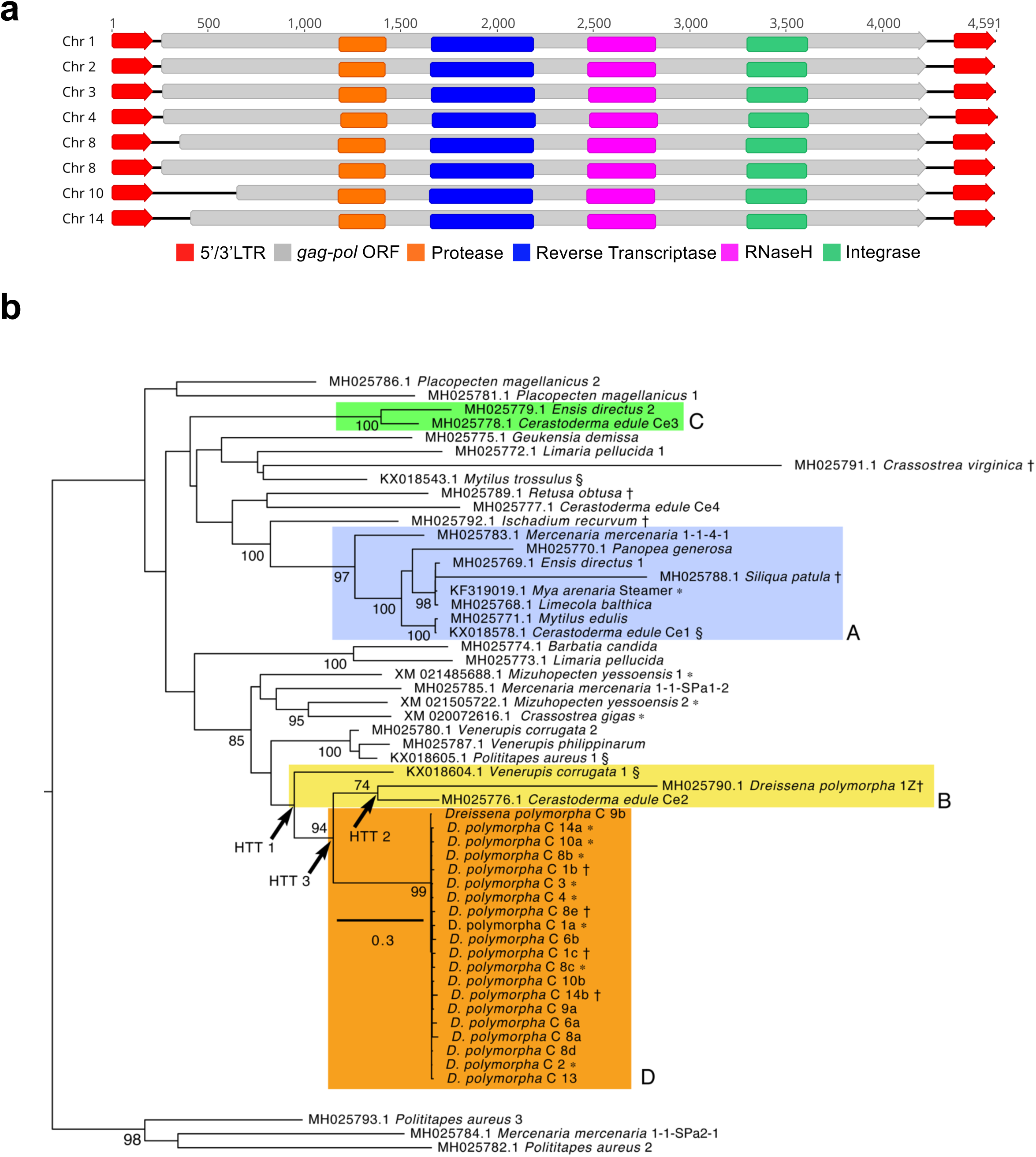
Steamer-Like Elements in the *D. polymorpha* genome. a) Schematics depicting the eight SLE copies, each with 2 LTRs flanking the longest ORFs among all similar elements in the *D. polymorpha* genome. b) Maximum likelihood phylogenetic tree of nucleotide sequences from the RNaseH-Integrase domain of *Gag-Pol* in *D. polymorpha* and other bivalve SLEs. The selected model^137^ of DNA sequence evolution was the GTR +G (rates *Γ*-distributed, *α* = 1.190) +I (estimated proportion of invariant sites = 0.011). The tree was rooted on the *Polititapes aureus* 2/3/*Mercenaria mercenaria* branch (bottom) and bootstrap support values > 70 are shown. Colored boxes A, B, and C contain taxa involved in all HTT events within bivalves that were identified previously^57^. Arrows label HTT events 1 and 2, identified previously^57^ and HTT 3, which we identified based on the same criteria. Together these account for two independent insertions of SLEs into zebra mussels. Clade D contains SLE sequences from the zebra mussel genome; “*D. polymorpha* C” = chromosomal location of the SLE, with letters to order multiple insertion sites. Taxon labels include NCBI Accession number, *taxon*, followed by isolate number or code. *\** = Sequence is from full length ORF encoding *Gag-Pol*, † = pseudogene sequence (one or more stop codons), § = sequence derived from neoplastic hemocytes^119^.

### Tissue-specific gene expression

We next conducted a number of RNA-Seq experiments in order to identify genes that are expressed in a tissue-specific manner, or genes that are regulated in response to different experimental conditions. We examined gene expression in the following tissues (Fig. 4a): mantle (the organ that secretes shell), gill (the focal organ for thermal stress response), and the foot (the organ that forms and attaches the byssal threads). RNA-Seq data from these three tissues was mapped to the reference containing the 68,018 annotated genes. A tissue-specificity index (*τ*) ^58^ was calculated and 577 genes exceeded the threshold of *τ* = 0.95 (Fig. 4b, Supplemental Fig. 9, Supplemental Files 5-7). Mantle contained the most tissue-specific genes—359 or 62.2% of the total unique transcripts. Tissue-specific genes had relatively little overlap with genes that were differentially expressed under the experimental conditions tested, suggesting that most tissue-specific genes are carrying out core as opposed to regulated functions (Supplemental Fig. 9, Supplemental Files 8-10).

**Figure 4.**
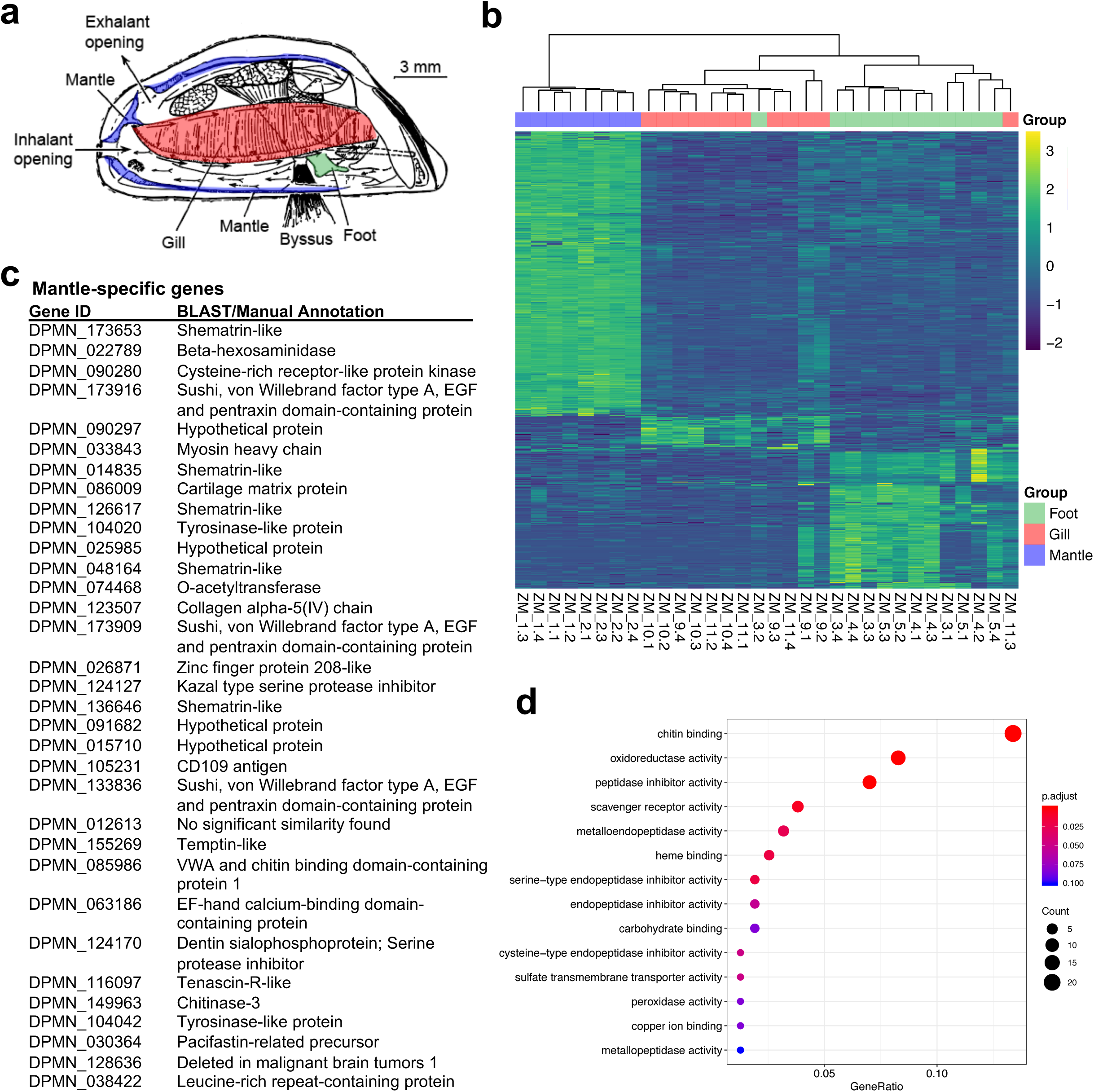
Tissue-specific gene expression patterns: mantle gene expression analysis. a) *D. polymorpha*: lateral view of the left valve with the right valve and the covering mantle fold removed to reveal the organs dissected for transcriptomes. In purple is the margin of the mantle tissue within the left valve. In *D. polymorpha* the mantle tissue is fused to form the siphons. Inhalent and exhalant siphon openings are pictured, as is the gill (ctenidium). Modified from ^138^. b) Heatmap depicting tissue-specific gene expression in the foot, gill, and mantle. c) List of the most highly expressed mantle-specific genes (tau > 0.95). d) Gene ontology term enrichment analysis for the mantle-specific genes.

### Mantle gene expression and shell formation

In dreissenids and other bivalves, the shell is constructed of calcium carbonate of different crystal forms (typically calcite in adult and aragonite in larval shells) that are deposited in an organic matrix, either through an extracellular or cell-mediated mechanism ^59,60^. Positive correlations have been found between shell strength/calcification and ambient Ca^2+^ has been found in some freshwater mollusk species, and selection favoring shell strength for predator defense has been found in others ^61,62^. In order to identify biomineralization-related genes we dissected mantle from adult zebra mussels. We collected mussels from both a calcium-rich (Lake Ore-be-gone: 35.4 mg/L) and a calcium-poor (Lake Superior: 14.4 mg/L) water body.

By inspecting highly expressed mantle-specific genes using the automated annotations as well as BLASTp and other manual annotation methods, we identified orthologs of a set of genes that have been previously implicated in shell formation (Fig. 4c, Supplemental File 11). These include tyrosinases, which are required for DOPA production, and other proteins that likely have structural roles, such as collagen. Six shematrin-like proteins are among the most specific and highly expressed mantle genes. Shematrins are glycine-rich proteins that have been identified in the mantle of other mollusks ^63–66^. Glycine-rich peptides in other organisms include structural proteins in rigid plant cell walls (60-70% glycine residues) as well as the major connective tissue in animals, collagen ^67,68^. The exact function of shematrins in shell formation is not clear, but their high expression levels and unusual structure is intriguing; *D. polymorpha* shematrins are characterized by arrays of G(n)Y repeats (Supplemental Fig. 10). Also highly expressed in the mantle were a number of Sushi, von Willebrand factor type A, EGF and pentraxin domain-containing proteins (SVEPs), which have been implicated in osteogenesis in mammals and have been identified in the mantle of other bivalves. In contrast to shell formation in pearl oysters^69^, no nacrein genes were identified in the zebra mussel genome and a tBLASTn search of the zebra mussel genome with *P. fucata* nacrein yielded no hits. Gene ontology term enrichment analysis also showed that the chitin binding molecular function was significantly enriched in the mantle-specific genes, along with a number of peptidase inhibitors (Fig. 4d).

Among the most specific and highly expressed mantle genes in *D. polymorpha* were two genes with sequence similarity to temptin, a pheromone that serves as a chemoattractant for mating in *Aplysia* ^70^. Zebra mussels attach to one another in clusters known as druses. Settlement of larvae near adults ^71^ and gregarious post-settlement behaviors^72^ create massive aggregations on lake and river bottom. These behaviors increase settlement success, enable “habitat engineering” in mussel beds ^72^, and may enhance feeding and fertilization success ^73–75^. Further investigation will be required to determine if the *D. polymorpha* temptin orthologs serve as aggregation cues or play some other unknown sensory role.

### Insights into byssal thread formation and attachment

The fibers that zebra and quagga mussels use to anchor themselves to hard surfaces are known as byssal threads. These are key innovations (absent from native North American and European freshwater mollusks) used to attach to conspecific mussels, and to native unionid mussels and other benthic animals, which can be smothered and outcompeted. Byssal attachment to boat hulls, docks, boat lifts and other recreational equipment allows rapid rates of spread between water bodies ^76–78^. Expression of genes during byssogenesis has been studied in zebra mussels^27^ but a majority of mRNAs that are up or down-regulated could not be identified.

Previous work identified a full byssal protein cDNA sequence (named Dpfp1) ^79,80^ and peptide fragments from a second foot byssal protein ^81^. More recent proteomic work also identified peptide tags associated with several *D. polymorpha* foot proteins (Dpfps) that are secreted by the foot and together form the stem, threads and attachment plaques (Fig. 5) of the byssus ^82,83^. The zebra mussel genome provided significant additional information to identify full-length proteins corresponding to the Dpfp peptide tags and to gain further insight into their function (Fig. 5). BLAST was used to align known Dpfp polypeptides against the genome-predicted protein sequences to determine the full Dpfp’s (Supplemental File 12). Dpfp2 and Dpfp12 were found to be the C-and N-terminal regions, respectively, of a single protein (Dpfp2). Similarly, Dpfp6 was found to be the C-terminal region of Dpfp1. Surrounding genomic regions on chromosome 8 were searched, and multiple exons were identified that are likely spliced to create mature Dpfp1 and Dpfp 6 (Supplemental File 12). Complete coding sequences were resolved for Dpfp5 and Dpfp8, and Dpfp8 was found to lack a signal peptide. Dpfp8 shows similarity to *Staphylococcus saprophyticus* cell wall associated fibronectin-binding protein (SCS67603.1). Putative homologs and paralogs to all other previously described byssal proteins were found in the zebra mussel genome, with Dpfp7, Dpfp9, Dpfp10 and Dpfp11 showing similarity to several protein coding DNA regions on multiple chromosomes. This information will inform future characterization of zebra mussel byssus formation and mechanism of adhesion.

**Figure 5.**
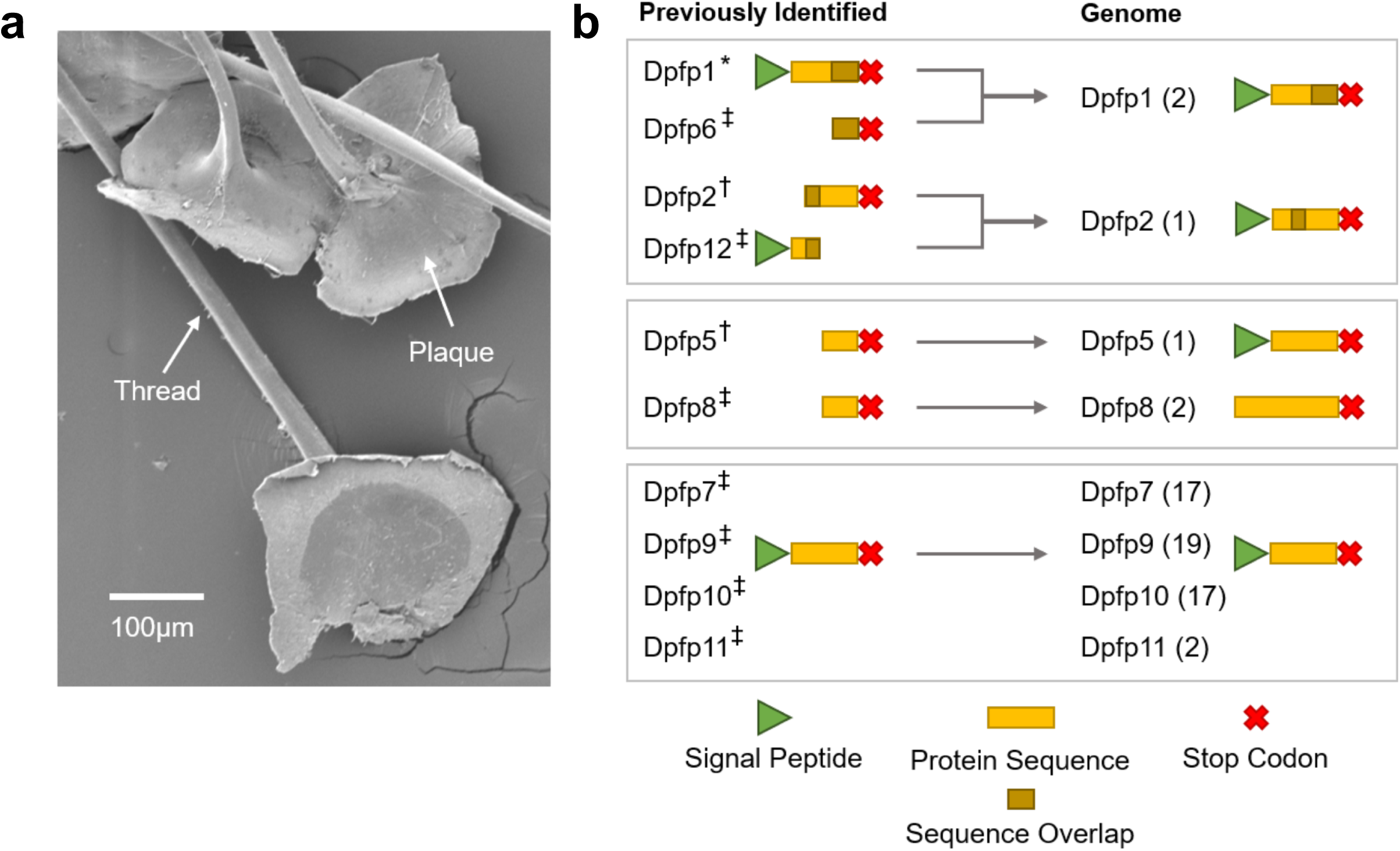
Analysis of proteins involved in byssal thread formation. a) SEM image of byssus, consisting of threads and plaques. b) Summary of the *D. polymorpha* foot protein (Dpfp) proteome re-analysis using the genome assembly. Proteins on the left were identified previously, from cDNA* sequencing ^79,80^, and from peptide sequencing using LC-MS/MS of soluble^†, 82^ and insoluble^‡, 83^ proteins extracted from freshly extruded byssal threads and ESTs from a foot cDNA library ^139^. The proteins on the right represent full-length Dpfp proteins identified in the genome, with the number of putative paralogous loci in parentheses.

We also examined transcripts from the foot following experimental induction of byssogenesis ^27^ (Supplemental Fig. 11). The foot distal to the byssus was dissected immediately after severing the byssal threads where they exit the shell valves, and four and eight days later. Changes were observed at the day-four time point, after which expression broadly returned to baseline by day eight (Supplemental Fig. 11). Some of the up-regulated genes were consistent with function identified in previous work on byssogenesis in the scallop ^48^, including tenascin-X (a connective protein) and a gene with phospholipid scramblase activity (Anoctamin-4-like, Supplemental Fig. 11, Supplemental File 13). In addition, there was a clear inhibition of the TNF pathway, with down-regulation of a TNF-ligand-like protein and up-regulation of Tax1BP1 (a negative regulator of TNF-signaling). The TNF pathway regulates inflammation and apoptosis, suggesting that production of the byssal thread may induce stress in the surrounding tissues and that this stress response may be actively suppressed. Consistent with this, both a cytokine receptor and the pro-apoptotic Bcl2-like gene are down regulated at the day-four time point. While earlier expression studies found otherwise ^27,82,83^ some byssal proteins were absent from our differentially expressed gene set. And while some of these proteins are differentially distributed across the byssus, localized expression in the foot has not been studied. Nevertheless, one explanation is that our dissections missed the secretory cells more proximal to the threads, a possibility that awaits testing.

### Thermal tolerance and chronic heat stress

In *Dreissena*, broad thermal tolerance and ability to adjust to local conditions have clearly played a role in invasion success. Zebra mussels have higher lethal temperature limits and spawn at higher water temperatures in North America than in Europe ^84,85^. In the Lower Mississippi River zebra mussels are found south to Louisiana. There they lack cooler water refuges, and persist near their lethal limit of 29-30°C for 3 months during the summer, while for 3 months in the river ranges from 5-10°C ^86^. In contrast, zebra mussels in the Upper Mississippi River encounter water temperatures > 25°C for just 1 month of the year, and < 2°C for about 3 months ^87^. Seasonal scheduling of growth and reproductive effort appears to be responsible for at least some of the adaptation or acclimation to conditions in the lower river, as populations in Louisiana shift their shell and tissue growth to the early spring and stop growing in summer ^86^ while more northerly populations grow tissue and spawn in summer months ^88,89^.

In order to identify genes involved in the response to thermal stress, we generated transcriptomes from gill tissue in animals exposed to periods of low (24°C), moderate (27°C), and high (30°C) chronic temperature stress (Fig. 6a). Moderate thermal stress led to the induction of a number of genes involved in cellular adhesion or cytoskeletal remodeling, including collagen, gelsolin, MYLIP E3 ubiquitin ligase, and N-cadherin (Fig. 6b, Supplemental File 14). High thermal stress led to strong induction of a large number of chaperones, including HSP70, DNAJ, Calnexin, and HSC70 (several of which were also induced to a lesser extent under moderate thermal stress), as well as the antioxidant protein cytochrome P450 (Fig. 6b-c, Supplemental File 14). The list of down-regulated genes was quite similar for both the moderate and high thermal stress conditions (Fig. 6d-e, Supplemental File 14). In addition to the induction of known stress-response genes, a number of genes with unknown function are also regulated by thermal stress, as is 4-Hydroxyphenylpyruvate Dioxygenase (HPPD), an enzyme which is involved in the catabolism of tyrosine (Fig. 6b-e, Supplemental File 14).

**Figure 6.**
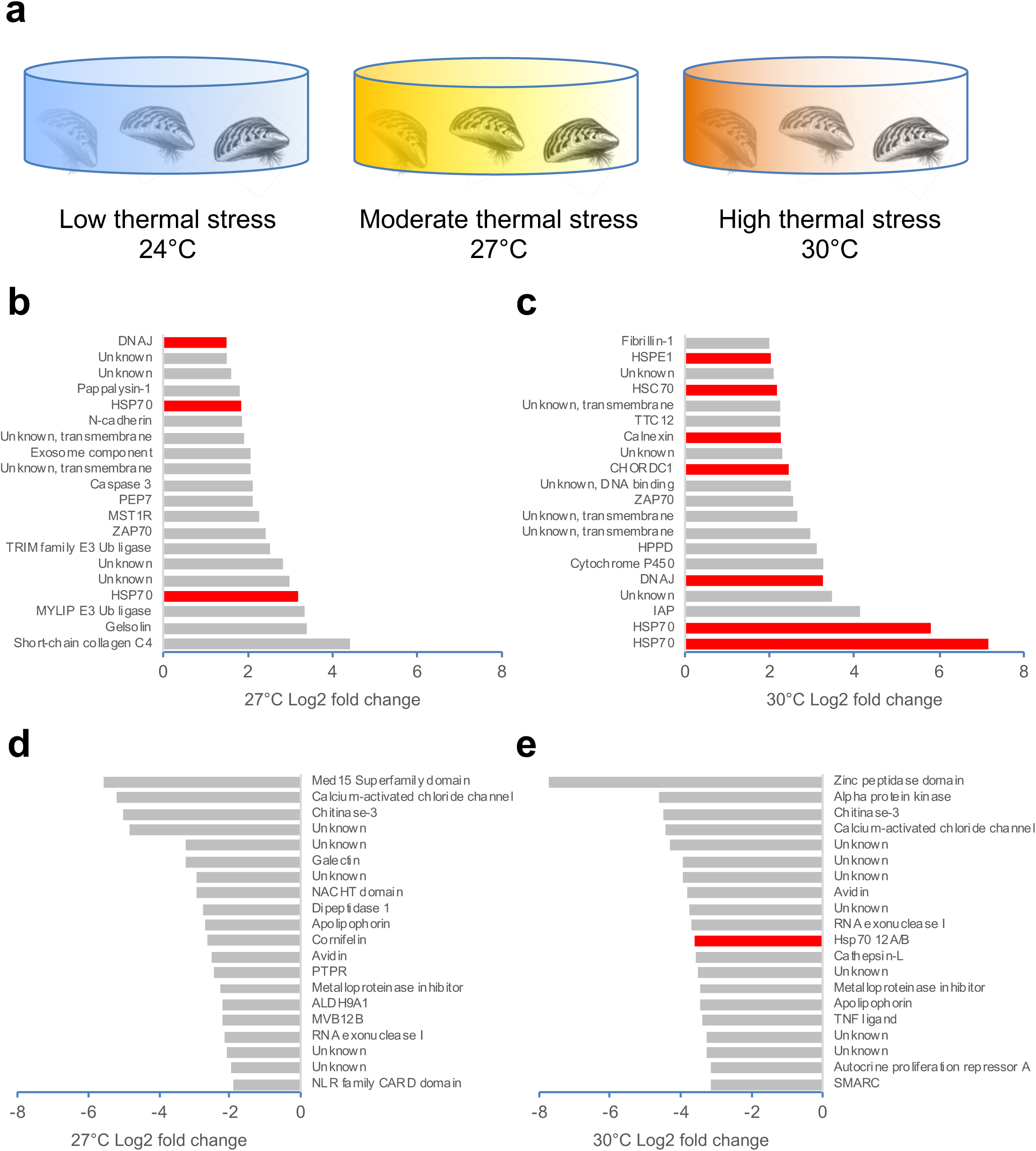
Response of *D. polymorpha* to thermal stress. a) Overview of experimental set-up. Animals were subjected to low (24°C), moderate (27°C), and high (30°C) thermal stress (n = 4 animals per condition). b) Top 20 genes up-regulated during moderate thermal stress by log2 fold-change. c) Top 20 genes up-regulated during high thermal stress by log2 fold-change. d) Top 20 genes down-regulated during moderate thermal stress by log2 fold-change. e) Top 20 genes down-regulated during high thermal stress by log2 fold-change.

## Discussion

Here we describe the genome of the zebra mussel. Consistent with the genomes of other bivalves, the *D. polymorpha* genome is highly repetitive and encodes an expanded set of heat-shock and anti-apoptotic proteins, presumably to deal with the challenges of a sessile existence. We examine the genetic underpinnings of several traits that have been linked to population growth and invasiveness, including shell and byssal thread formation, and response to thermal stress. While these analyses uncovered multiple genes and pathways that seem to function in a conserved manner across multiple bivalve species, they also uncovered a large number of genes of unknown function. In the future, it will be of considerable value to compare the zebra mussel genome with that of its congener, the quagga mussel (*D. rostriformis*), in order to gain further insights into ecological displacement of zebra by quagga mussels, and to investigate genetic underpinning of their relative invasiveness, such as comparative work on byssogenesis that may help account for the slower geographic spread of quagga mussels.

The existence of genomic resources for *D. polymorpha* and the catalog of genes we have identified will enable multiple new lines of investigation, as well as provide researchers with an improved tool for population genetic experiments, for instance, tracking the spread of mussels using Genotyping-by-Sequencing (GBS) approaches, or designing new targeted assays for the presence or activity of zebra mussels.

While it is clear that changes in transportation networks (e.g. canal building, opening of shipping channels, ballast water discharge) were the events that initiated primary invasions of European and North American waters^5,90^, several biological characteristics are responsible for the rate of spread of zebra and quagga mussels across both continents, while other traits have limited their suitable habitat range. Genomics offers a path to understanding these traits at the genetic level, which may ultimately guide the development of control methods and management strategies.

## Methods

### Genomic DNA extraction and PacBio Library Creation

Zebra mussel individuals were collected by SCUBA from off the Duluth waterfront beach (46.78671°N, −92.09114°W), in Lake Superior in June of 2017. Mature adults were dissected. To sex the animals, gonad squashes were prepared and examined under a compound microscope for gametes, and a set of large males (25-30 mm shell length) were selected for genome sequencing and analysis. Genomic DNA was extracted using the Qiagen Genomic Tip 100/G kit, with all tissues (except gut) from each selected individual split across six total extractions to prevent clogging of Genomic Tips. Pooled extractions from one chosen individual yielded >100 ug genomic DNA as assessed by PicoGreen DNA quantification (ThermoFisher). The Agilent TapeStation Genomic DNA assay indicated that the majority of gDNA extracted was well over 20kb (not shown). Further analysis by Pulsed-Field Gel Electrophoresis indicated a broad distribution from 20-120kb, with a modal size of 40kb (not shown).

Thirty µg of gDNA was sheared by passing a solution of 50 ng/uL DNA through a 26G blunt-tipped needle for a total of 20 passes. This sheared DNA was cleaned and concentrated using AMPurePB beads with a 1X bead ratio, and further library preparation was performed following the PacBio protocol for >30kb libraries using the SMRTbell®Template Prep Kit 1.0. Size-selection of the final library was carried out using the >20kb high-pass protocol on the PippinHT (Sage Science), and an additional PacBio DNA Damage Repair treatment was performed following size-selection.

### PacBio Sequencing

Sequencing was carried-out on a PacBio Sequel between November 2017 and February 2018 using 1M v2 SMRT Cells with 2.1 chemistry and diffusion loading.

### Nanopore Library Creation and Sequencing

Genomic DNA from the individual used for PacBio sequence was prepared for Nanopore sequencing using the Oxford Nanopore Ligation Sequencing Kit (SQK-LSK109). The resulting library was sequenced on a single Oxford Nanopore R9.4.1 flowcell on a GridION X5. Reads were collected in MinKNOW for GridION release 18.07.9 (minknow-core-gridion v. 1.15.4) and basecalled live with guppy v. 1.8.5-1.

### Illumina Polishing Library Creation and Sequencing

High molecular weight DNA from the individual used for the PacBio sequencing was also used as input for Illumina TruSeq DNA PCR-Free library creation, targeting a 350 bp insert size. The resulting library was sequenced on a single lane of HiSeq 2500 High Output (SBS V4) in a 2 × 125 cycle configuration, yielding 68 Gb of data representing ~37X coverage of the genome.

### Hi-C Library Creation and Sequencing

A previously frozen male individual from the same collection date and site in Lake Superior was thawed and mantle, gonad, and gill tissues were dissected using a razor blade. This was a different mussel, because insufficient tissue remained after earlier DNA extractions of the other mussel for genome assembly and polishing. Hi-C library creation was carried out with a Proximo™ Hi-C kit (Feb 2018) from Phase Genomics using the Proximo™ Hi-C Animal Protocol version 1.0. This method is largely similar to previously published protocols ^91^. The resulting library was sequenced on a single lane of HiSeq 2500 High Output (SBS V4) in a 2 × 125 cycle configuration, yielding 234M clusters passing filter.

### Sample collection for transcriptome studies

*Mantle*. Adult zebra mussels (20-25 mm shell length) were collected from a high-Ca^2+^ (35-38 mg/L) site: the Lake Ore-Be-Gone mine pit in Gilbert, MN (47.4836°N, −92.4605°W) and from a “low-Ca^2+^” (14.4 mg/L) site: Lake Superior near the Duluth Lift Bridge (46.7867°N, 92.0911°W). Mussels and water were collected underwater by SCUBA, and mussels were stored on ice and returned to the laboratory for dissection within six hours. This approach was used in lieu of experimental manipulations, because chronic exposure to low calcium concentrations are difficult to achieve in the laboratory—slow shell growth and poor survival have been observed in these marginal (< 15 mg/L) concentrations^92^. Calcium concentration in unfiltered, undigested lake water was determined by 15-element ICP-OES on the iCAP 7600 (Thermo-Fisher, Waltham MA).

#### Gill and foot

For these transcriptomes, experiments were used to study differential gene expression in adult mussels that were housed in aquaria for several weeks where they were acclimated, fed laboratory diets, then exposed to experimental treatments. Zebra mussels (15-22 mm shell length) were collected from sites in Lake Minnetonka (44.9533 ° N, −93.4870° W and 44.8980° N, −93.6688° W) and Lake Waconia (44.8711° N, −93.7596° W) then transported in coolers to the University of Minnesota, where they were acclimated, 100 mussels per each of 12 × 40 L glass aquaria with flowing well water (4 L/minute) at 20°C (unheated). Temperature was checked twice daily with digital probes. Mussels were fed 1.8 ml per tank of liquid shellfish diet (Reed Mariculture, Campbell CA) once daily, with water flow shut off for 1.5 hours for feeding. Tank temperatures were raised to 24-25°C over three days by mixing in heated well water; then temperatures were held constant over seven days for acclimation.

Experimental treatments followed, with each group of 4 tanks raised 1°C per day (using a 200 W aquarium heater in each tank) to target temperatures of 25, 27 and 30°C then maintained at target for seven days. For gill transcriptomes, two mussels per each of four treatment tanks were removed, then both ctenidia were dissected and preserved in 750 μL RNAlater per animal at −20°C. For foot, mussels from Lake Waconia, attached firmly to rocks and maintained for seven days in each of two of the 25°C tanks above were selected. Byssal threads were severed where they enter the shell valves to induce byssus growth and reattachment. Immediately thereafter, foot tissue (distal tip region) was dissected from each of eight animals (for a time-zero control) and preserved in RNAlater. Byssus-cut animals were painted with nail polish and placed onto rocks in each of two tanks at 25°C. Mussels that firmly attached overnight were observed for four days and eight days after reattachment, and four firmly attached mussels per time point were selected and foot tissue was dissected and preserved as above. Metadata for transcriptome samples is in Supplemental File 15.

### RNA-Seq Sample Preparation, Library Creation, and Sequencing

Zebra mussel tissue RNA was extracted using the Qiagen RNeasy Plus Universal kit from tissues stored at −20°C in RNA*later*™ (Ambion, Carlsbad, CA). RNA concentration was assessed using Nanodrop, and quantified fluorometrically with the RiboGreen RNA assay kit (ThermoFisher). Further evaluation was based on RNA Integrity Number (RIN) scores generated by the Agilent TapeStation 2200 Eukaryotic RNA assay. Samples with RIN >9.0 and RNA mass >500 ng were used as input for library preparation. Libraries were prepared using the TruSeq® Stranded mRNA kit (Illumina) and sequenced on a HiSeq 2500 High Output (SBS V4) run in a 2 × 50 cycle configuration, generating approximately 15M reads per sample (Mean = 15.8 M, 15% CV).

### Genome Assembly

The primary assembly was generated using Canu 1.7^35^ from 167.8 Gbp of PacBio subreads over 1kbp in length with the command:

> canu -p asm -d asm ‘genomeSize=2g’ ‘correctedErrorRate=0.105’ ‘corMinCoverage=4’
>
> ‘corOutCoverage=100’ ‘batOptions=-dg 3 -db 3 -dr 1 -ca 500 -cp 50’
>
> ‘corMhapSensitivity=normal’.

The assembly used heterozygous parameters due to the relatively high heterozygosity of the sample (2.13% estimated from Genoscope ^93^ and previous Illumina sequencing). BUSCO v3 ^40^ was run using the metazoa_odb9 gene set with the command:

> python run_BUSCO.py -c 16 --blast_single_core -f --in asm.contigs.fasta -o SAMPLE -l –
>
> m metazoa_odb9 genome.

The assembly had 93.9% core metazoan complete genes with 35.2% single copy complete and 58.7% duplicated complete genes. Purge haplotigs ^36^ was run to remove redundancy in the assembly with the commands:

> minimap2 -ax map-pb --secondary=no -t 16 asm.contigs.fasta reads.fasta.gz >
>
> reads.sam
>
> samtools view -b -T asm.contigs.fasta -S reads.sam > reads.bam
>
> samtools sort -O bam -o reads.sorted.bam -T tmp reads.bam
>
> samtools index reads.sorted.bam purge_haplotigs readhist reads.sorted.bam
>
> purge_haplotigs contigcov -i reads.sorted.bam.genecov -l 15 -m 80 -h 120 -j 200
>
> purge_haplotigs purge -t 32 -g asm.contigs.fasta -c coverage_stats.csv –b
>
> reads.sorted.bam -windowmasker

Unassigned contigs were removed from the primary set leaving 1.80 Gbp in 2,863 contigs with an N50 of 1,111,027 bp.

### Genome Polishing

The resulting contigs were re-analyzed using the PacBio standard polishing pipeline – GenomicConsensus v2.3.3 ^94^, which derives a better genomic consensus through long read mapping and variant calling using an improved Hidden Markov Model implemented in the algorithm Arrow. The polished draft assembly was further corrected for Indels using Pilon ^95^ with setting: --fix indels --threads 32 --verbose --changes --tracks. A single contig corresponding to the PacBio sequencing control was removed from the final assembly.

### Repeat analysis

RepeatModeler ^96^ was used to identify repeat families from the primary haploid genome. The resulted unknown repeat families were combined with the default full RepeatMasker ^97^ database. RepeatMasker scanned the primary haploid genome sequences for the combined repeat databases in quick search mode.

### Hi-C Scaffolding

Chromatin conformation capture data was generated using a Phase Genomics (Seattle, WA) Proximo Hi-C Animal Kit v1.0, which is a commercially available version of the Hi-C protocol ^91^. Following the kit protocol, intact cells from two samples were crosslinked using a formaldehyde solution, digested using the *Sau3A*I restriction enzyme, and proximity-ligated with biotinylated nucleotides to create chimeric molecules composed of fragments from different regions of the genome that were physically proximal *in vivo*, but not necessarily proximal in the genome. Continuing with the manufacturer’s protocol, molecules were pulled down with streptavidin beads and processed into an Illumina-compatible sequencing library. Sequencing was performed in a single lane of Illumina HiSeq 2500 High Output (SBS V5) in a 2×125 cycle configuration, yielding 230,479,044 clusters passing filter.

Reads were aligned to the draft assembly also following the manufacturer’s recommendations ^98^. Briefly, reads were aligned using BWA-MEM ^99^ with the -5SP and -t 8 options specified, and all other options default. SAMBLASTER ^100^ was used to flag PCR duplicates, which were later excluded from analysis. Alignments were then filtered with samtools ^101^ using the -F 2304 filtering flag to remove non-primary and secondary alignments and further filtered with matlock ^102^ (default options) to remove alignment errors, low-quality alignments, and other alignment noise due to repetitiveness, heterozygosity, and other ambiguous assembled sequences.

Phase Genomics’ Proximo Hi-C genome-scaffolding platform was used to create chromosome-scale scaffolds from the corrected assembly as described in Bickhart et al. ^37^. As in the LACHESIS method ^103^, this process computes a contact frequency matrix from the aligned Hi-C read pairs, normalized by the number of *Sau3*AI restriction sites (GATC) on each contig, and constructs scaffolds in such a way as to optimize expected contact frequency and other statistical patterns in Hi-C data. Approximately 140,000 separate Proximo runs were performed to optimize the number of scaffolds and construction of to make them as concordant with the observed Hi-C data as possible. This process resulted in a set of 16 chromosome-scale scaffolds containing 1.79 Gb of sequence (98.4% of the contig assembly), with a scaffold N50 of 107.56 Mb and a scaffold N75 of 92.27 Mb.

### Mitochondrial Genome Assembly, Polishing, Mapping, and Annotation

Mapping of PacBio reads to an initial Canu assembly for the mitochondrial genome indicated a small region of very high coverage (Supplemental Fig. 5). An alternate assembly of the mitochondrial genome was substituted which was generated in parallel in FALCON 0.5 (length_cutoff = −1, seed_coverage = 30, genome_size = 2.7G) and which did not collapse this repeat sequence. This assembly was polished for indels via Pilon using Illumina reads as with the nuclear genome, and a single substitution error in the coding region was manually edited (c.14475 C>A, G184W) based on strong support from Illumina reads (data not shown). The mitochondrial genome was annotated based a previously published partial mitochondrial sequence ^26^ in Geneious using the “Annotate from Database” function with a 98% similarity cutoff. The origin point was set to place the tRNA-Val annotation at base 48, matching the previously published sequence.

PacBio and Nanopore reads were mapped against a reference file containing two concatenated copies of the mitochondrial genome sequence, in order to allow reads to map across the origin. Alignments were generated with minimap2 -ax using settings map-pb and map-ont, respectively. Visualization of the resulting alignments (Fig. 2c) was performed using a custom tool, ConcatMap (https://github.com/darylgohl/ConcatMap). Illumina reads from the polishing library were mapped (Supplemental Fig. 5) to the final, polished mitochondrial genome using BWA-MEM^94^.

### Hi-C analysis of the mitochondrial contig

Ten contigs ranging in size from 50kb to 100kb were selected from each of the pseudo-chromosome scaffolds. The total number of Hi-C contacts between each selected contig and each pseudo-chromosome was determined. The same analysis was performed using the mitochondrial contig, then all Hi-C link counts were normalized by dividing the number of contacts between a contig and pseudo-chromosome by the total number of Hi-C contacts associated with the contig. The resulting normalized data was visualized using ggplot2 to develop boxplots that compare the number of links for contigs based on their association with each pseudo-chromosome.

### Transcriptome Assembly

Reads from all zebra mussel RNAseq libraries were pooled for transcriptome assembly. A database of ribosomal RNA was downloaded from SILVA^104–106^, restricting the entries to Bivalvia. The combined RNAseq reads were cleaned of putative ribosomal RNA sequences using “BBDuk” from the BBTools suite of scripts^107^, treating the Bivalvia ribosomal RNA as potential contaminants, using a k-mer size of 25bp and an edit distance of 1. Reads that passed this filter were then assembled with Trinity 2.8.4 ^108^ with a “RF” library type, *in silico* read normalization, and a minimum contig length of 500bp. Assembled transcripts from Trinity were then searched against the non-redundant nucleotide sequence database hosted by NCBI, current as of 2018-10-09. A maximum of 20 target sequences were returned for each transcript, restricted by a minimum of 10% identity and a maximum E-value of 1 × 10^−5^. Assembled transcripts that matched sequences derived from non-eukaryotes or synthetic constructs were discarded.

### Differential Expression Analysis

RNAseq reads were checked for quality issues, adapter content, and duplication with FastQC 0.11.7. Cleaning for sequencing adapters, trimming of low-quality bases, and filtering for length were performed with Trimmomatic 3.3 ^109^. The adapter sequences that were targeted for removal were the standard Illumina sequencing adapters. Quality trimming was performed with a window size of 4bp and a minimum mean quality score of 15. Reads that were shorter than 18bp after trimming were discarded.

Reads were aligned to the HiC-scaffolded genome assembly draft with HISAT2 2.1.0 ^110^, with putative intron-exon boundaries inferred with genes with functional annotation from the draft annotation and a bundled Python script. Read pairs in which one read failed quality control were not used in alignment and expression analysis. BAM files from HISAT2 were cleaned of reads with a mapping quality score of less than 60 with samtools 1.7. Cleaned alignments were used to generate expression counts with the featureCounts program in the Subread package v. 1.6.2 ^111^. Both reads in a pair were required to map to a feature and be in the proper orientation for them to be counted. Raw read counts were imported into R 3.5.0 ^112^ for analysis with edgeR 3.24.3 ^113^. Genes that were less than 200bp were removed from the counts matrix. Tests for differential expression were performed between experimental conditions within tissue. For each tissue, genes with low expression were filtered in the following way: genes in which at least X samples with fewer than 10 were removed, where X is the size of the condition with the fewest replicates. Tests for differential expression used a negative binomial model for dispersion estimation, and genes showing significant levels of differential expression were identified with a quasi-likelihood F test implemented in edgeR ^114^. Genes were identified as differentially expressed if they had a nominal *P*-value of less than 0.01 in the output from the ‘glmQLFTest’ function.

### Tissue Specificity Calculation

Filtered, normalized counts were used to calculate *τ*, a measure of tissue specificity: ^58^

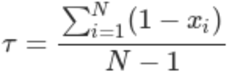

Where *N* is the number of tissues analyzed and *x_i_* are the normalized counts. Normalized and log-transformed counts-per-million (CPM) values for each gene were estimated with edgeR. The mean CPM for samples from each tissue were treated as the expression values for that tissue. *τ* was then calculated for each gene. Genes with *τ* of 0.95 or greater were considered to be specific to the tissue with highest expression.

### Identification of Steamer-Like Elements (SLEs) and phylogenetic analysis

A sequence amplified from *D. polymorpha* using SLE-targeting degenerate primers ^57^ was used as the basis for an initial BLAST search of the genome assembly. Dotplots of the sequence surrounding hits were analyzed to identify fifty putative LTR sequences, and these were aligned to build a consensus LTR sequence specific to our assembly. A subsequent BLAST search with this consensus sequence was performed, and surrounding sequence context was examined for the presence of long (>3 kb) ORFs between flanking LTRs. Eight intact elements identified with these criteria were aligned based on coding sequence (ClustalW), and annotated based on NCBI Conserved Domain search.

First, we evaluated phylogenetic evidence that zebra mussel TEs are SLEs. Amino acid sequences for the full-length *Gag-Pol* polyprotein region from these eight elements and from the *Steamer* element from *Mya arenaria* (Accession AIE48224.1) were aligned to a database of the *Gypsy/T3y* family of LTR-retrotransposons ^115^, using MAFFT^116^ and the E-INS-i method. The alignment included 2078 residues and 105 sequences. The model of sequence evolution was selected based on the AIC option in SMS^117^, using the option to estimate amino acid frequencies from the data. A maximum likelihood genealogy was built using PhyML^118^, using the NNI tree topology search and the BIONJ starting tree options, and support for nodes was evaluated based on 100 bootstrap replications.

Next, we used DNA sequence genealogies to further investigate whether HTT events led to insertions of 20 SLEs that we found in the zebra mussel genome that contained 2 LTRs flanking an intact *Gag-Pol* ORF, including the 8 elements above. From GenBank, we downloaded sequences from multiple bivalve species, from the region located between the RNase H and integrase domains of *Gag-Pol* that was amplified using degenerate primers^57^. We added 3 sequences of long ORFs from *Gag-Pol* that were cloned from neoplastic tissue^119^, 3 that were obtained from *Crassostrea gigas* and *Mizuhopecten yessoensis* genome projects, and the full length *Steamer* clone from *Mya arenaria*. We used MAFFT and the G-INS-1 progressive method to align nucleotide sequences based on the translated amino acid sequences and trimmed the ends. The alignment of 54 sequences and 1074 nucleotide positions was loaded into PhylML and the maximum likelihood tree was constructed using the above options (except that in this case, nucleotide frequencies were ML optimized).

### Genome Annotation

Functional annotation was carried out with Funannotate 1.0.1^120^ in haploid mode using transcript evidence from RNA-seq alignments, *de novo* Trinity assemblies, and genome-guided Trinity assemblies. First, repeats were identified using RepeatModeler^96^ and soft-masked using RepeatMasker ^97^. Second, protein evidence from a UniProtKB/Swiss-Prot-curated database (downloaded on 26 April 2017) was aligned to the genomes using tBLASTn and exonerate^121^, and transcript evidence was aligned using GMAP^122^. Analysis *ab initio* used gene predictors AUGUSTUS v3.2.3^123^ and GeneMark-ET v4.32^124^, trained using BRAKER1^125^, and tRNAs were predicted with tRNAscan-SE^126^. Consensus protein coding gene models were predicted using EvidenceModeler^127^, and finally gene models were discarded if they were more than 90% contained within a repeat masked region and/or identified from a BLASTp search of known transposons against the TransposonPSI^128^ and Repbase^129^ repeat databases. Any fatal errors detected by tbl2asn (https://www.ncbi.nlm.nih.gov/genbank/asndisc/) were fixed. Functional annotation used the following databases and tools: PFAM^41^, InterPro^42^, UniProtKB^43^, Merops^44^, CAZymes^45^, and a set of transcription factors based on InterProScan domains^130^ to assign functional annotations.

### Comparison to Eastern Oyster (*Crassostrea virginica*) Proteins

Zebra mussel genes with functional annotation information were used to identify groups of orthologous genes with Eastern Oyster (*Crassostrea virginica*). Annotated protein sequences from *C. virginica* were downloaded from the C_virginica-3.0 assembly and annotation hosted on NCBI. Zebra mussel protein sequences and *C. virginica* protein sequences were grouped into orthologous groups using OrthoFinder version 2.2.7 ^131^, OrthoFinder was run with BLASTP 2.7.1 for similarity searches, MAFFT 7.305 for alignment, MCL 14.137 for clustering, and RAxML 8.2.11 for tree inference.

### Porting annotations to improved Hi-C scaffolds

Annotations were created against a preliminary set of scaffolds and ported to the final set of scaffolds following Juicebox error correction. Several files were used to perform this process. First, from the preliminary set of scaffolds, annotations were generated in GFF format as described above. For both the preliminary and final set of scaffolds, a .assembly file ^132,133^ and a set of .ordering files ^133^ were produced by the Proximo pipeline ^37^. Finally, the original contig FASTA was obtained. These data were used to identify the location and orientation of each contig in the final scaffolds relative to the preliminary scaffolds, and to generate a new GFF file reflecting the new position of each annotation in the final scaffolds. Annotations were checked with a custom Python script that compares the sequences that correspond to the gene regions on each assembly. Gene sequences were extracted from the assemblies by strand-aware retrieval of the “gene” features in the GFF3 file; that is, if a gene is annotated on the “minus” strand, then the sequence is reverse-complemented. The resulting sequences were then compared between assembly versions to ensure that the re-calculated annotations corresponded to exactly the same sequence. In cases of sequence mismatch, the sequence based on re-calculated annotations was reverse-complemented to diagnose strand conversion issues. Annotations which spanned more than one contig in the preliminary set of scaffolds were discarded during this process; it is likely such annotations, which spanned placeholder unestimated gaps between contigs, were spurious calls ^134^. The script used for porting the annotations is available here: https://github.com/phasegenomics/annotation_mover.

### Accession codes

The *D. polymorpha* genome assembly is available at NCBI (BioProject: PRJNA533175). Sequencing data files are available through the NCBI Sequence Read Archive (BioProject: PRJNA533175, PRJNA533176). Pending data release by NCBI, the *D. polymorpha* genome and annotations can be downloaded using the following links:

Genome sequence: https://zebra_mussel.s3.msi.umn.edu/Dpolymorpha_Assembly.V2.Final_wMito.fasta.gz Annotations: https://zebra_mussel.s3.msi.umn.edu/Dpolymorpha_Assembly.V2.Final_wMito.gff3.gz

## Acknowledgements

SK is supported by the Intramural Research Program of the National Human Genome Research Institute, National Institutes of Health. This work utilized the computational resources of the NIH HPC Biowulf cluster (https://hpc.nih.gov) and the Minnesota Supercomputing Institute (https://www.msi.umn.edu). Funding was from the Minnesota Environment and Natural Resources Trust Fund and the Minnesota Aquatic Invasive Species Research Center.

## Author contributions

MAM and DMG conceived and designed experiments, analyzed data, and wrote the paper. BA prepared PacBio and Hi-C libraries, analyzed data, and wrote the paper. TK, YZ, JA, and KATS analyzed data and helped to assemble and annotate the genome. SM designed experiments, collected samples, isolated DNA. JG analyzed data. AO and EDS analyzed byssal thread attachment proteins. AB carried out sequencing of PacBio and Illumina libraries. JPB carried out nanopore sequencing. AH and HM analyzed data. IL, HM, and SS carried out Hi-C-based scaffolding. SK generated the Canu assembly and ran purge haplotigs. KBB conceived and designed experiments.

## Supplemental Figure Legends

**Supplemental Figure 1.**
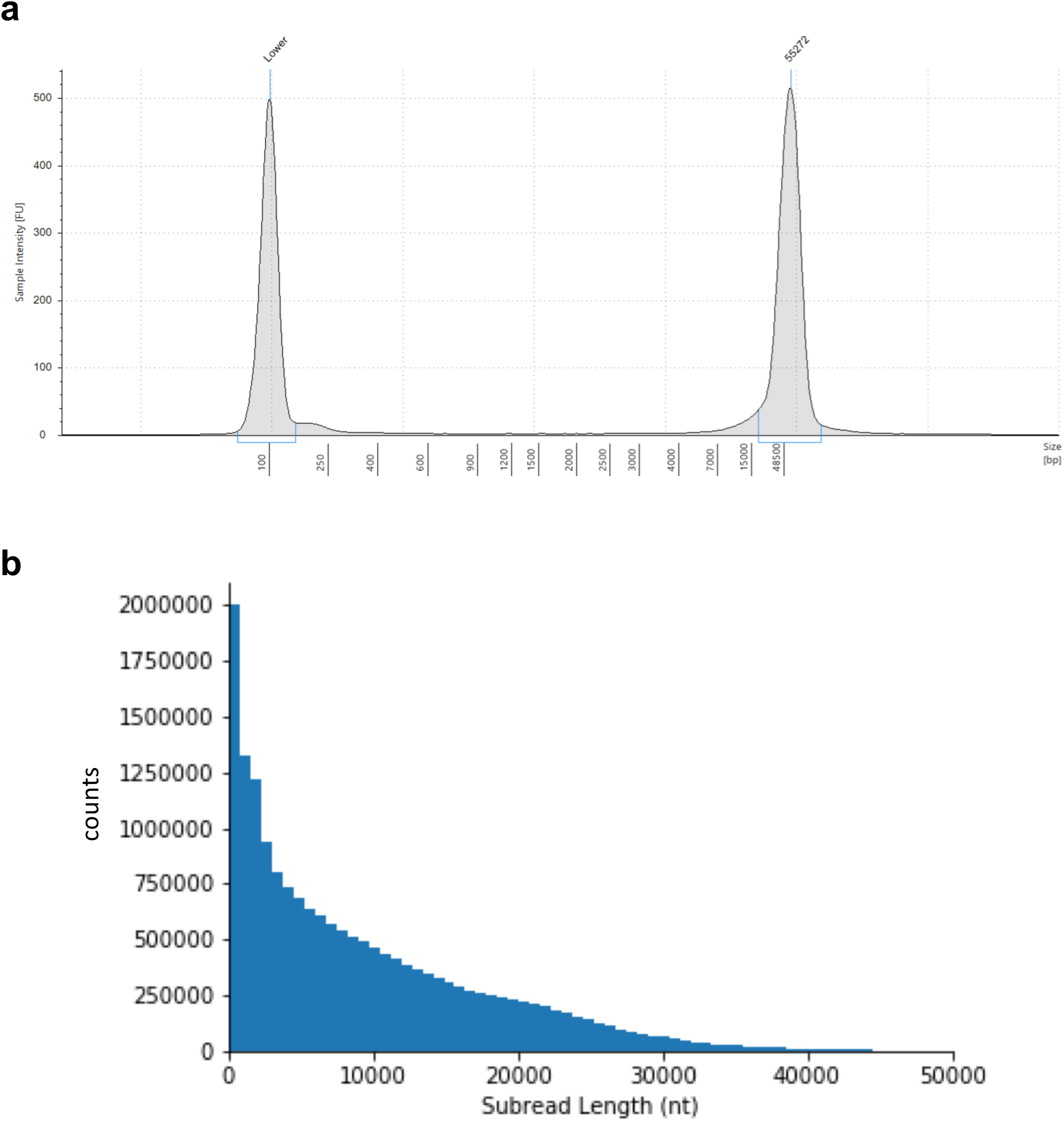
PacBio sequencing library and sequencer output. a) Size-selected PacBio *D. polymorpha* sequencing library. b) Distribution of subread lengths in the PacBio sequencing data set.

**Supplemental Figure 2.**
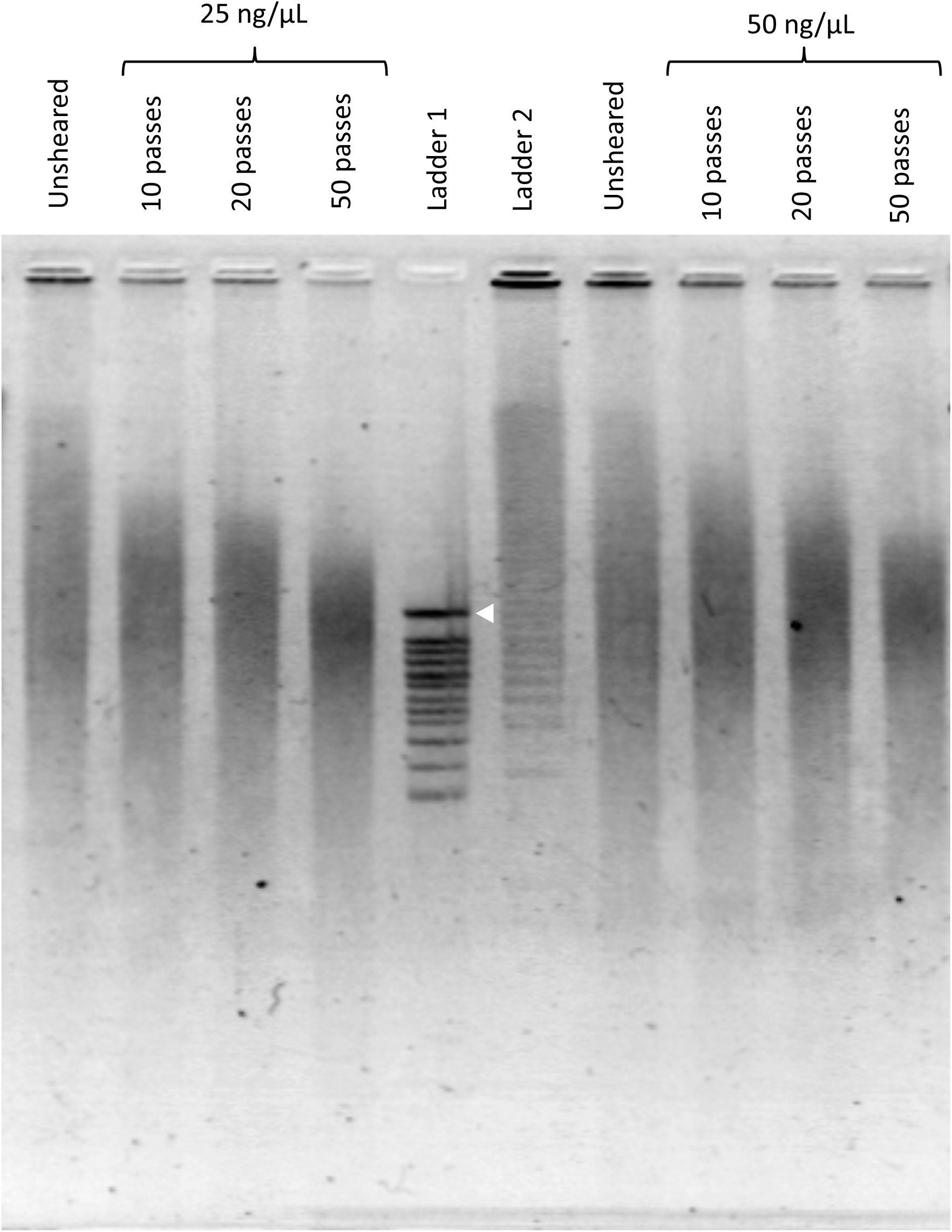
Pulsed-field Gel Electrophoresis of input gDNA. Concentrations indicate DNA concentration at time of shearing. 200 ng DNA loaded for gel visualization in each well. Image cropped, inverted, and adjusted for brightness/contrast. Ladder 1: CHEF DNA Size Standard 8-48 kb (1703707) Ladder 2: CHEF DNA Size Standard 5 kb (1703624). White arrowhead indicates 48.5 kb.

**Supplemental Figure 3.**
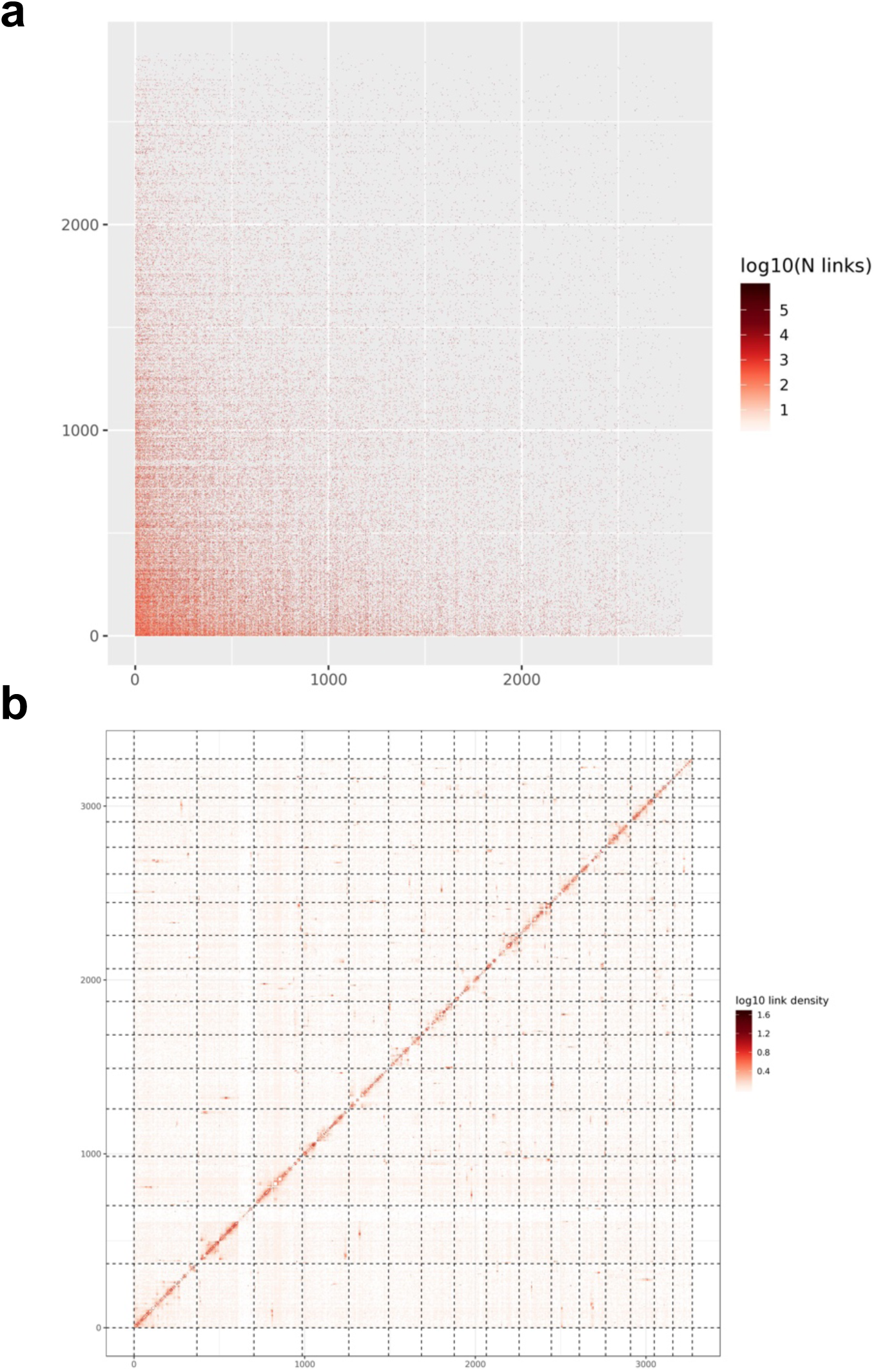
Hi-C pre- and post-scaffolding heatmaps. a) Hi-C pre-scaffolding heatmap. b) Hi-C post-scaffolding heatmap.

**Supplemental Figure 4.**
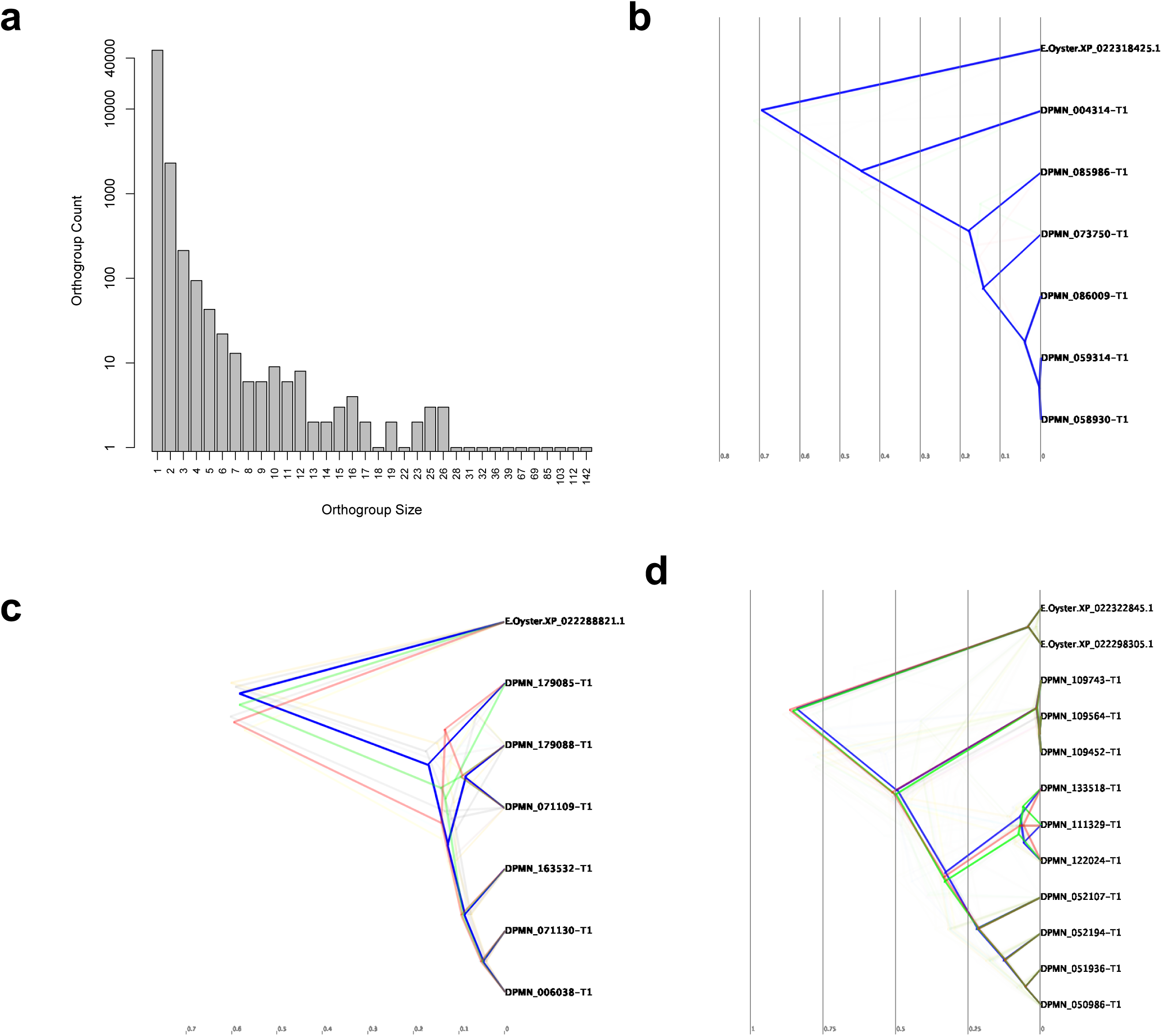
Divergence trees for select orthogroups. a) Histogram of paralog group counts for the *D. polymorpha* genome. b-d) BEAST trees with date estimates for the expansion of several gene families in *D. polymorpha* relative to the Eastern oyster (*C. virginica*). b) OG0000776. c) OG0000670. d) OG0000272.

**Supplemental Figure 5.**
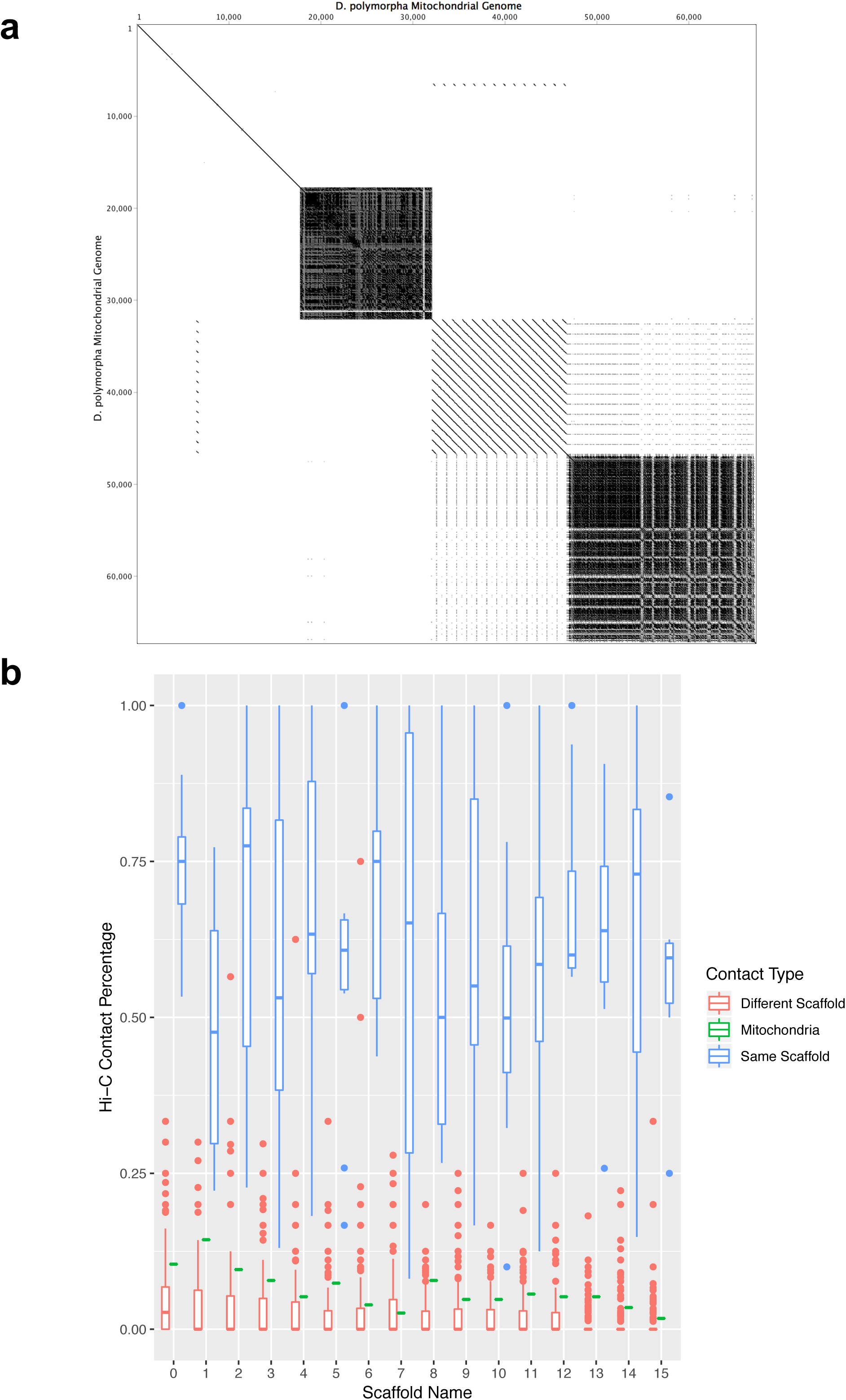
Structure of the *D. polymorpha* mitochondrial genome. a) Dotplot (self-self) of D. polymorpha mitochondrial genome demonstrating large blocks of repetitive sequence elements. Generated in Geneious based on EMBOSS tool dottup. Word size=15, tile size = 20kb. b) Comparison of the normalized number of Hi-C links comes from a contig within the same scaffold versus the normalized number of Hi-C links from a contig assembled on a different scaffold. The links observed between the mitochondrial contig and the chromosomal scaffolds is comparable to the number of links between contigs assembled on different scaffolds.

**Supplemental Figure 6.**
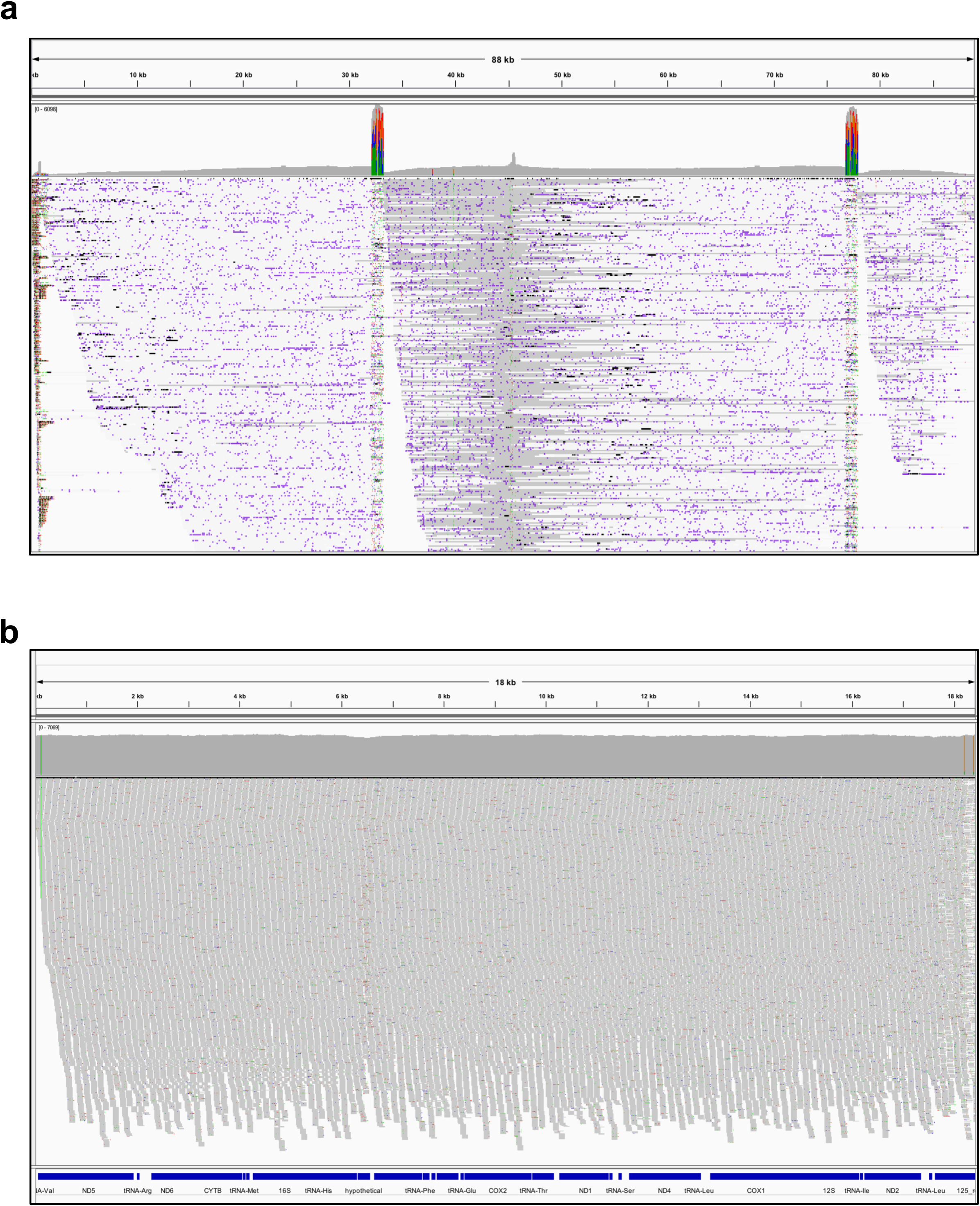
Alignment of PacBio and Illumina reads to the mitochondrial genome. a) Coverage plot of Minimap2-aligned PacBio reads against initial Canu-assembled mitochondrial genome (concatenated) showing area of high coverage which was determined to be a collapsed repeat sequence. b) Paired-end Illumina reads from mixed somatic/germline male tissue were aligned to the mitochondrial genome, demonstrating a lack of SNPs that might otherwise indicate heteroplasmy. Only the coding region is shown, as unambiguous mapping of short reads to highly repetitive sequences is unreliable. Allele threshold for coverage plot = 0.05.

**Supplemental Figure 7.**
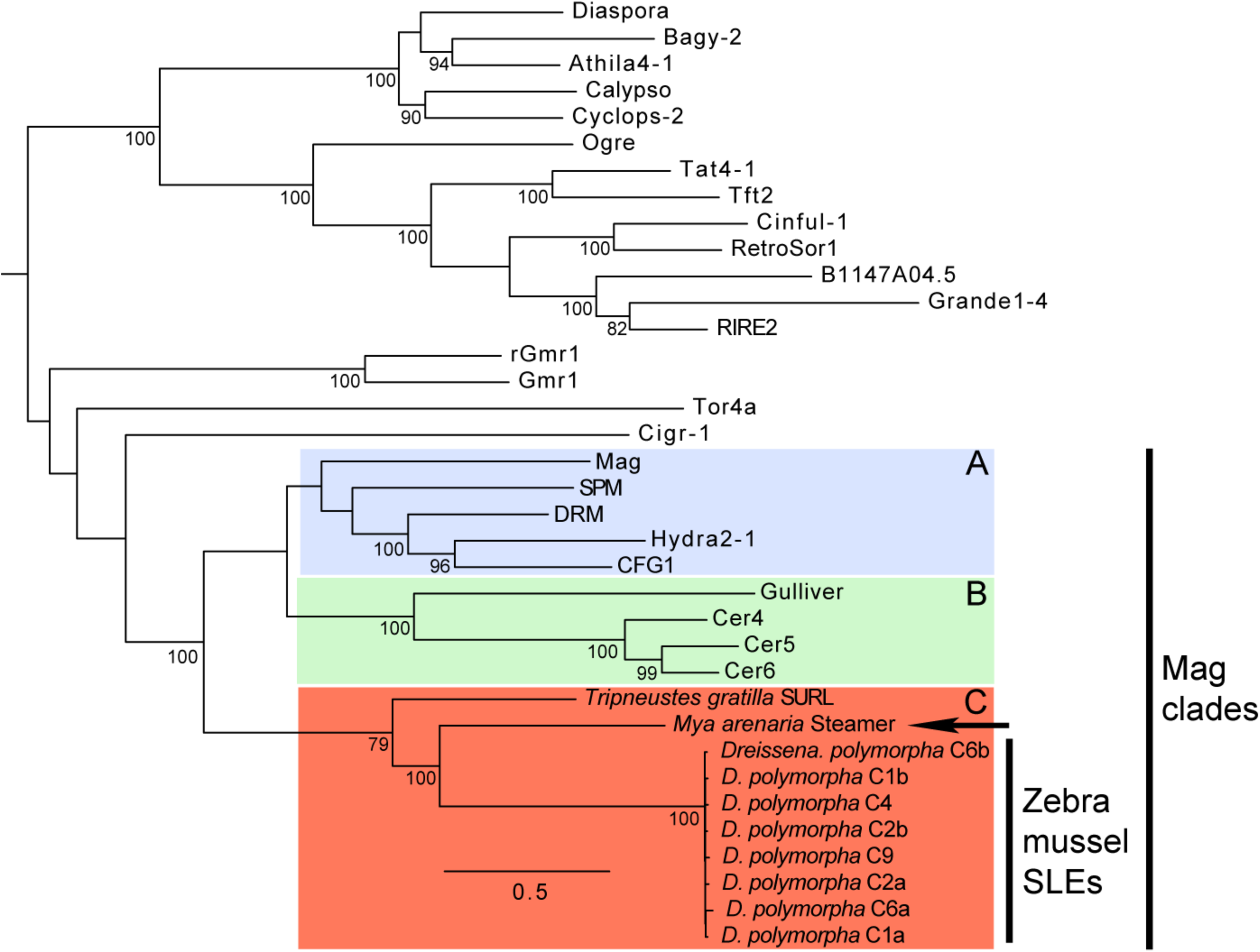
Phylogenetic tree of the *Ty3/Gypsy* family of retrotransposons, including putative SLEs in zebra mussels. Maximum likelihood phylogenetic tree of amino acid sequences from the entire *Gag-Pol* region. The selected model of amino acid sequence evolution was the LG^140^ model +G (rates Gamma-distributed, *α* = 1.566) +I (estimated proportion of invariant sites =.001)+ F (amino-acid frequencies estimated from the data). The analysis included all sequenced elements from branch 2 of the *Ty3/Gypsy* family (including LTR retrotransposons and non-chromodomain retroviruses^141^, but only the Mag clades (A, B, and C in colored boxes) are shown, along with the sister clade. *Steamer* (arrow) from *Mya arenaria* groups with the sea urchin retroelement SURL, in clade C^55^. Here we show that the *D. polymorpha* elements are sister to *Steamer*, confirming that they are SLEs. Bootstrap support values > 70 label the nodes, and the scale bar is expected changes per site from maximum likelihood.

**Supplemental Figure 8.**
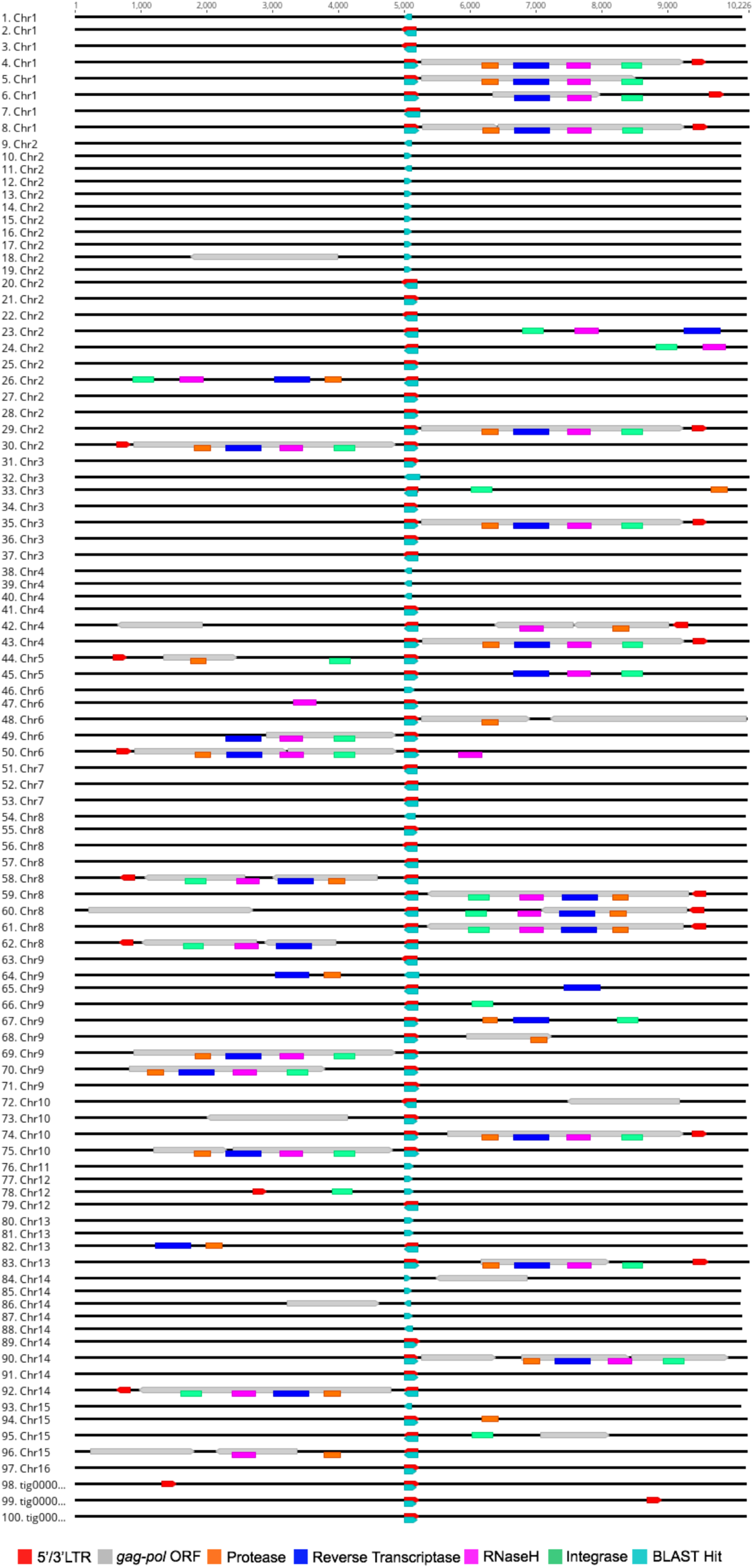
Partial SLEs in the *D. polymorpha* genome. Schematic depicting incomplete SLEs, including LTR-only sequences in the *D. polymorpha* genome. Sequences are centered on an LTR element and additional annotated domains in the SLE ORF are colored as indicated in Fig. 3a.

**Supplemental Figure 9.**
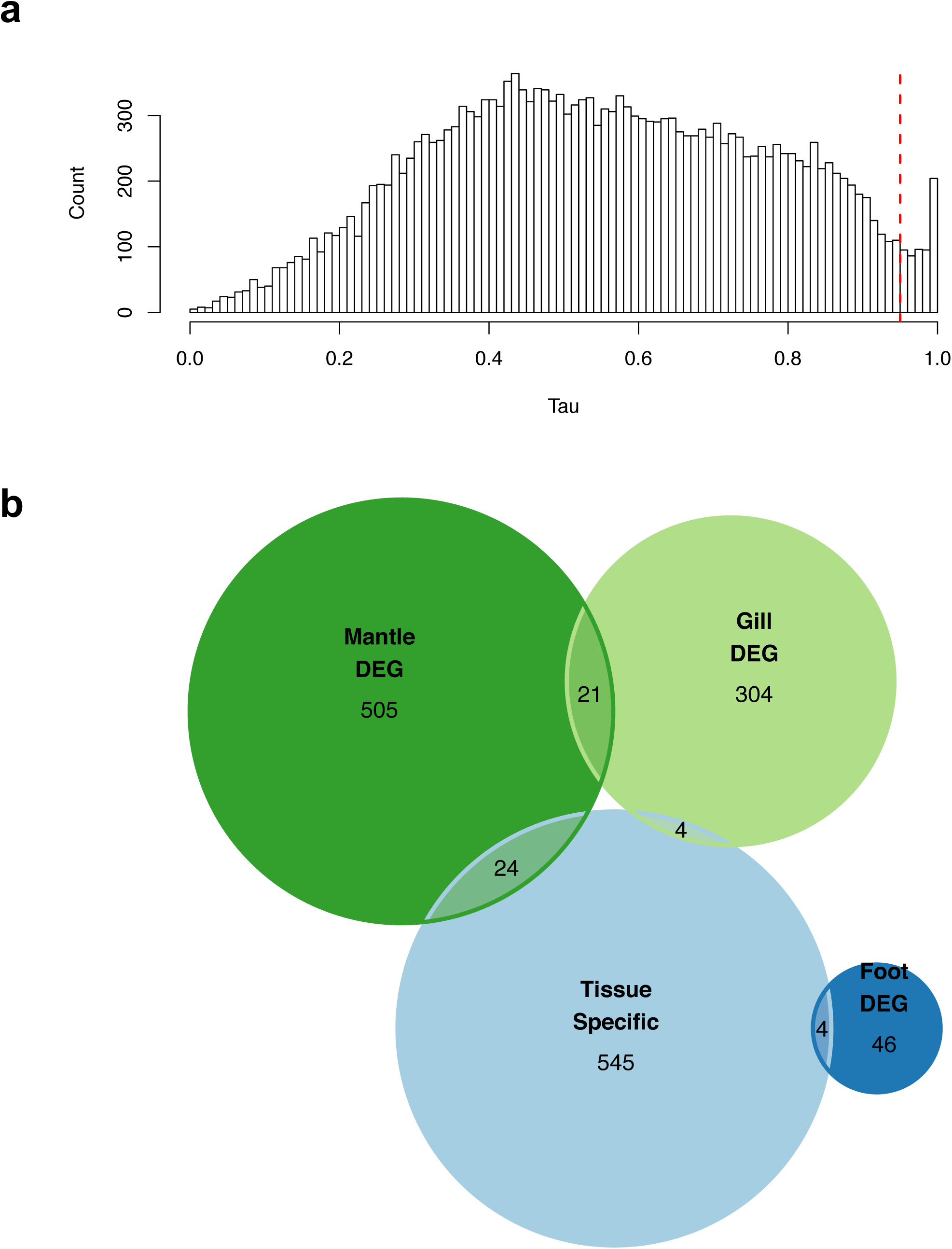
Tissue-specificity scores. a) Histogram depicting the distribution of tissue-specificity (tau) scores for *D. polymorpha* genes. A cut-off of 0.95 was used to define tissue-specific expression (dashed line). b) Venn diagram of tissue-specific genes compared to genes that were differentially expressed under different experimental conditions.

**Supplemental Figure 10.**
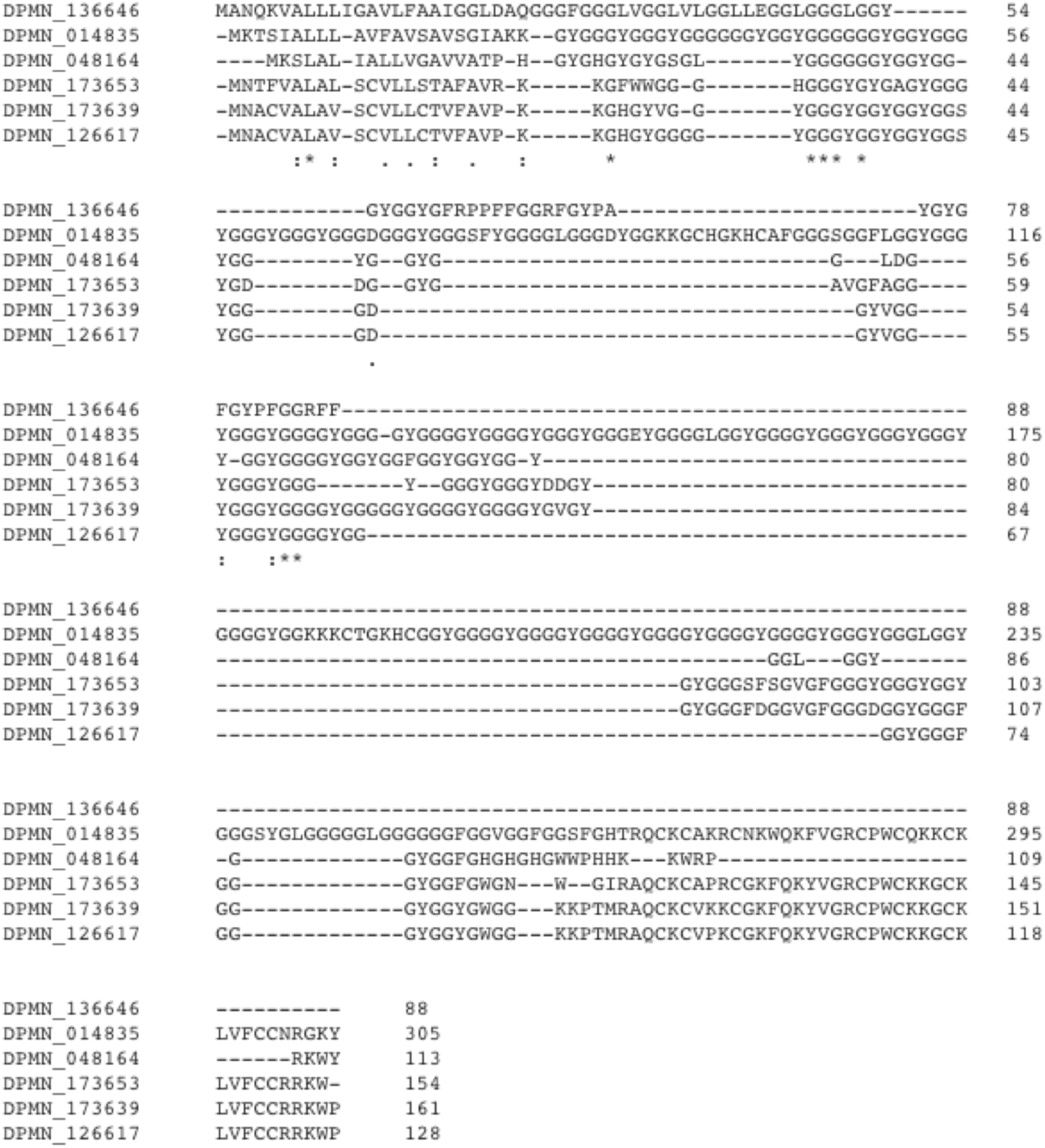
*D. polymorpha* shematrin-like proteins. Multiple sequence alignment of the six shematrin-like proteins identified in the *D. polymorpha* genome using CLUSTAL Omega (https://www.ebi.ac.uk/Tools/msa/clustalo/).

**Supplemental Figure 11.**
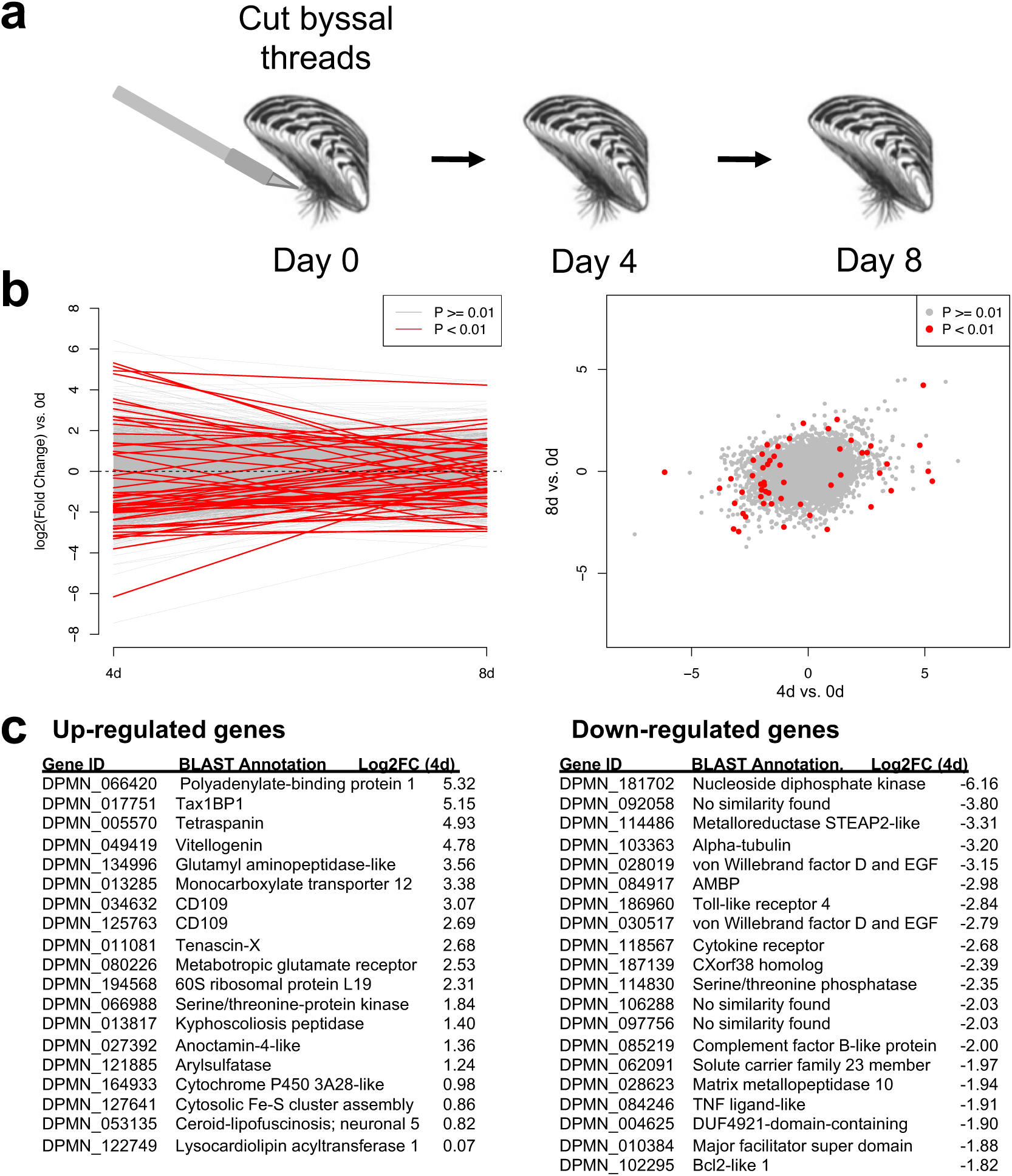
Foot gene expression during byssal thread formation. a) Schematic depicting experimental design; byssal threads were severed at day zero, and dissected foot tissue was collected on days zero, four, and eight (n = 4 animals per condition). b) Gene expression changes (log2 fold-change) relative to the day zero time point. c) List of up- and down-regulated genes in the foot at the day-four time point.

**Supplemental Table 1.** Sequenced bivalve genomes.

| Species | Family | Common name | Commercial interest | Assembly level | Number of scaffolds | Number of contigs | Contig N50 (bp) | Genome length (Mb) | Reference |
| --- | --- | --- | --- | --- | --- | --- | --- | --- | --- |
| *Bathymodiolus platifrons* | Mytilidae | Hydrothermal vent mussel | None | Scaffold | 65,662 | 272,497 | 12,602 | 1,658.2 | Sun et al. 2017 |
| *Chlamys farreri* | Pectinidae | Zhikong (Chinese) scallop | Wild harvest and culture | Scaffold | 96,024 | 148,999 | 21,500 | 779.9 | Li et al. 2017 |
| *Crassostrea gigas* | Ostreidae | Pacific oyster | Hatchery culture—leads aquatic animals in global harvest | Scaffold | 7,659 | 30,460 | 31,239 | 557.7 | Zhang et al. 2012 |
| *Crassostrea virginica* | Ostreidae | Eastern oyster | Wild harvest and hatchery culture | Chromosome | 11 | 669 | 1,971,208 | 684.7 | Gómez-Chiarri et al. 2015 |
| *Mizuhopecten (Patinopectin) yessoensis* | Pectinidae | Yesso scallop | Culture from wild seed | Scaffold | 82,659 | 120,022 | 65,014 | 987.6 | Wang et al. 2017 |
| *Modiolus philippinarum* | Mytilidae | Phillipine horse mussel | None | Scaffold | 74,573 | 301,873 | 18,389 | 2,629.6 | Sun et al. 2017 |
| *Pinctada martensii* | Pteriidae | Akoya pearl oyster | Cultured pearls | Chromosome | 5,039 | 85,944 | 21,518 | 991.0 | Unpublished |

**Supplemental Table 2.**
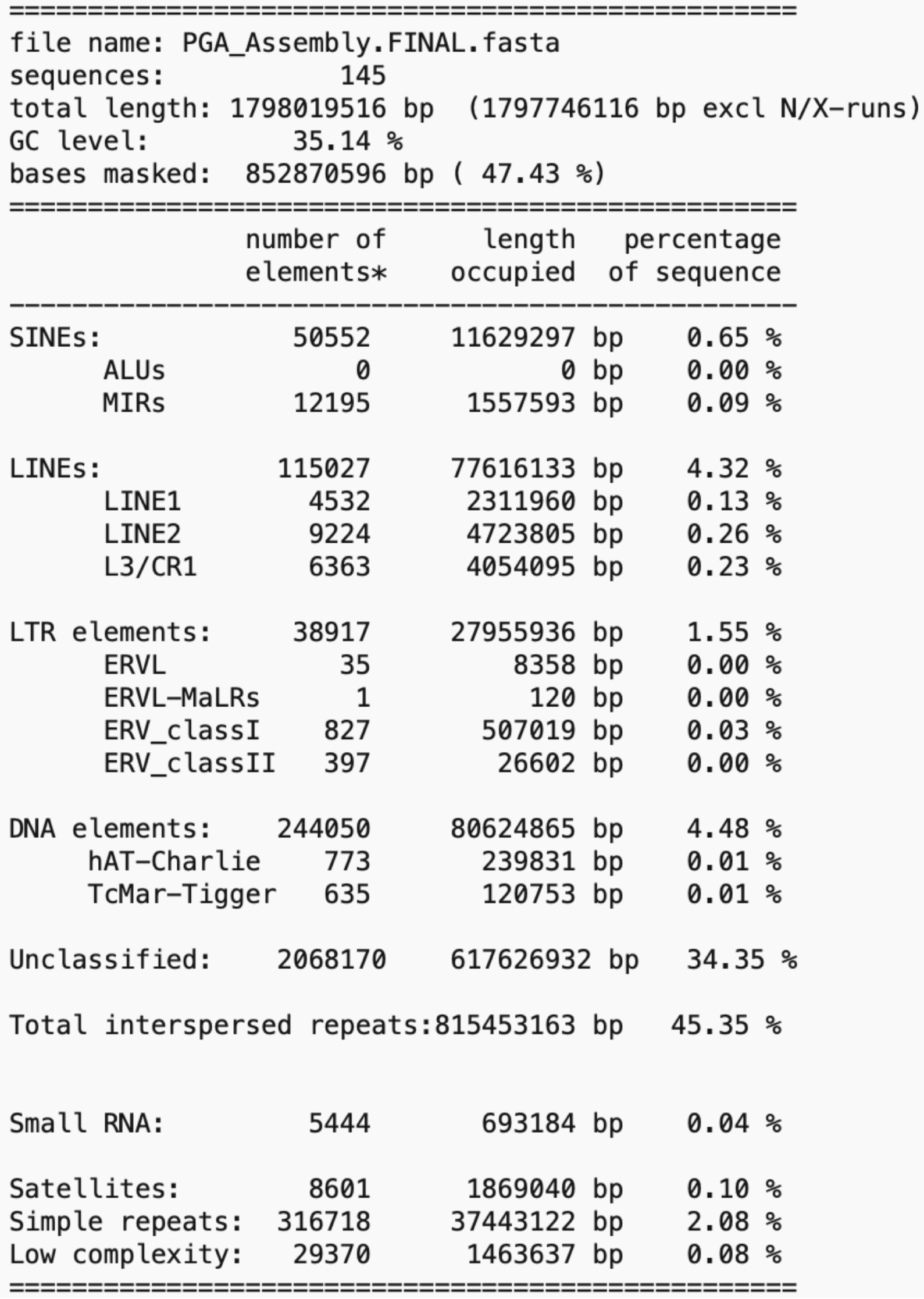
RepeatMasker output.

## Supplemental Files

**Supplemental File 1. Gene IDs for zebra mussel paralogous groups.**

ZM_Paralogous_Groups.csv

**Supplemental File 2. Annotation of zebra mussel paralogous groups.**

ZM_GFF_Paralogous_Groups.xlsx

**Supplemental File 3. Orthogroup gene IDs for zebra mussel and Eastern oyster comparison.**

ZM_vs_EO_Orthogroups.txt

**Supplemental File 4. Annotated orthogroups for zebra mussel and Eastern oyster comparison.**

Orthogroups_EO_ZM_Products.xlsx

**Supplemental File 5. Mantle-specific genes.**

Mantle-specific_genes.csv

**Supplemental File 6. Gill-specific genes.**

Gill-specific_genes.csv

**Supplemental File 7. Foot-specific genes.**

Foot-specific_genes.csv

**Supplemental File 8. Differentially expressed genes in mantle in mussels collected from high and low calcium environments.**

Mantle_DEGs.csv

**Supplemental File 9. Differentially expressed genes in the gill in mussels exposed to different thermal stress conditions.**

Gill_DEGs_.csv

**Supplemental File 10. Differentially expressed genes in the foot in response to severing the byssal threads.**

Foot_DEGs.csv

**Supplemental File 11. Mantle-specific BLAST results.**

Mantle-specific_BLAST_results.xlsx

**Supplemental File 12. Identification of full length Dpfp proteins.**

Dpfp_analysis.docx

**Supplemental File 13. Foot DGE BLAST results.**

Foot_DEG_BLAST_results.xlsx

**Supplemental File 14. Gill DGE BLAST results.**

Gill_DEG_BLAST_results.xlsx

**Supplemental File 15. Metadata associated with RNA-Seq samples.**

Transcriptome_metadata.xlsx

