## Supplementary material for "The Genome of the Zebra Mussel, *Dreissena polymorpha*: A Resource for Invasive Species Research": Dpfp_analysis.docx

**Supplemental Information: Analysis of Byssal Proteins in the *D. Polymorpha* genome**

**Summary**

- Dpfp12 is the missing N-terminal of Dpfp2	**(Page 2)**
- Completed sequence of Dpfp5	**(Page 5)**
- Dpfp6 is not found in any predicted proteins, but is found in unannotated exons	**(Page 10)**
- Dpfp8 matches significantly to the C-terminus of DPMN_074797, which does not have a signal peptide	**(Page 26)**
- All byssal proteins except for Dpfp6 (which was found in unannotated exons) are found in the predicted proteins from the ZM genome, with some proteins matching to several predicted proteins (Dpfp7, Dpfp9, and Dpfp10)	**(Page 30)**

**Dpfp12 is the missing N-terminal of Dpfp2**

Protein DPMN_127665, predicted from the ZM genome, is the byssal protein Dpfp2. This protein is the full construct of two previously identified ZM byssal peptide sequences, Dpfp12 and Dpfp2. Dpfp12 was reported in 2014 in the insoluble byssal extract to contain a signal peptide but no stop codon [1]. Dpfp2 was reported in 1993 as a DOPA containing ZM byssal protein [2] and a more complete sequence of Dpfp2 was determined in 2013 [3]; it did not contain a starting Met residue nor a signal peptide. BLASTp of Dpfp12 and Dpfp2 against the predicted proteins from the ZM genome all match to protein DPMN_127665. Dpfp12 matched to residues 1-63 and Dpfp2 matched to residues 53-155; the peptide sequence YQEKTYPGYPP is common between Dpfp12 and Dpfp2. Thus, the previously determined to be separate sequences of Dpfp2 and Dpfp12 are in fact two components of the same protein, DPMN_127665, where Dpfp12 is the N-terminal and Dpfp2 is the C-terminal. The complete Dpfp2 sequence is:

MFSAAALLLLVSFYGTASGQYWNSYRPYPVYPPKQTYPSYPDKKYPSYPEKTYQEKTYPGYPPK**Q**A**Y**PVYPEKTYPEKTYPAYPTKKSYPEYPEKTYTKKTYEAYPTKDSYTVYPDKKYTEKKYEAYPTKQSYPVYPEKKYPEKPYPGYQDYWGQ

Blue is from Dpfp12, Yellow is from Dpfp2, and Green is found in both. Underlined is the signal peptide. Bold Q = glutamine deamidation, Bold Y = tyrosine hydroxylation (DOPA).

[1] Gantayet, A., Rees, D.J., and Sone, E.D. 2014. Novel Proteins Identified in the Insoluble Byssal Matrix of the Freshwater Zebra Mussel. *Marine Biotechnology* **16**: 144-155. DOI: 10.1007/s10126-013-9537-9.

[2] Rzepecki, L.M. and Waite, J.H. 1993. The byssus of the zebra mussel, *Dreissena polymorpha*. II: Structure and polymorphism of byssal polyphenolic protein families. *Molecular Marine Biology and Biotechnology* **2(5)**: 267-279.

[3] Gantayet, A., Ohana, L., and Sone, E.D. 2013. Byssal proteins of the freshwater zebra mussel, *Dreissena polymorpha*. *Biofouling* **29(1)**: 77-85. DOI: 10.1080/08927014.2012.746672.

BLASTp matches of Dpfp2 and Dpfp12 against ZM genome predicted protein DPMN_127665

Highlighted are the regions that match to DPMN_127665.

Dpfp2-532_**AM229730:** APGRHGGRGNSISSGRPGRYQEKTYPGYPPKQAYPVYPEKTYPEKTYPAYPTKKSYPEYPEKTYTKKTYEAYPTKDSYTVYPDKKYTEKTYEAYPTKDSYTVYPDKKYTEKKYEAYPTKQSYPVYPEKKYPEKPYPGYQDYWGK*IHTP*QQGSQRNGKNYVPRPRPR*SLVNSRG

Query 22	EKTYPGYPPKQAYPVYPEKTYPEKTYPAYPTKKSYPEYPEKTYTKKTYEAYPTKDSYTVY 81

++TYP YP K+ YP YPEKTY EKTYP YP K++YP YPEKTY +KTY AYPTK SY Y

Sbjct 34	KQTYPSYPDKK-YPSYPEKTYQEKTYPGYPPKQAYPVYPEKTYPEKTYPAYPTKKSYPEY 92

Query 82	PDKKYTEKTYEAYPTKDSYTVYPDKKYTEKKYEAYPTKQSYPVYPEKKYPEKPYPGYQDY 141

P+K YT+KTYEAYPTKDSYTVYPDKKYTEKKYEAYPTKQSYPVYPEKKYPEKPYPGYQDY

Sbjct 93	PEKTYTKKTYEAYPTKDSYTVYPDKKYTEKKYEAYPTKQSYPVYPEKKYPEKPYPGYQDY 152

Query 142 WGK 144

WG+

Sbjct 153 WGQ 155

Dpfp12a-352_**AM230355:**

NSLVISSGRPGR*FLRERIQRKTMFSAATLLLLVSFYGTASGQYWNSYRPYPVYPPKQTYPSYPDKKYPSYPEKTYLGRDHANRIPAA

Query 24 MFSAATLLLLVSFYGTASGQYWNSYRPYPVYPPKQTYPSYPDKKYPSYPEKTYLGRDHAN 83

MFSAA LLLLVSFYGTASGQYWNSYRPYPVYPPKQTYPSYPDKKYPSYPEKTY + +

Sbjct 1	MFSAAALLLLVSFYGTASGQYWNSYRPYPVYPPKQTYPSYPDKKYPSYPEKTYQEKTYPG 60

Query 84 RIP 86

P

Sbjct 61 YPP 63

Dpfp12b-352_**AM230369:**

NSLVISSGRPGRWIQRNTMFSAAALLLLVSFYGTASGQYWNSYRPYPVYPPKQTYPSYPDKKYPSYPEKTYLGRDHANRIPAA

Query 19 MFSAAALLLLVSFYGTASGQYWNSYRPYPVYPPKQTYPSYPDKKYPSYPEKTYLGRDHAN 78

MFSAAALLLLVSFYGTASGQYWNSYRPYPVYPPKQTYPSYPDKKYPSYPEKTY + +

Sbjct 1	MFSAAALLLLVSFYGTASGQYWNSYRPYPVYPPKQTYPSYPDKKYPSYPEKTYQEKTYPG 60

Query 79 RIP 81

P

Sbjct 61 YPP 63

Dpfp12c-531_**AM230302:**

GRGNSISVVAAEVRFLRERIQRKTMFSAATLLLLVSFYGTASGQYWNSYRPYPVYPPKQTYPSYPDKKYPSYPEKTSLVN

Query 25 MFSAATLLLLVSFYGTASGQYWNSYRPYPVYPPKQTYPSYPDKKYPSYPEKT 76

MFSAA LLLLVSFYGTASGQYWNSYRPYPVYPPKQTYPSYPDKKYPSYPEKT

Sbjct 1	MFSAAALLLLVSFYGTASGQYWNSYRPYPVYPPKQTYPSYPDKKYPSYPEKT 52

**Dpfp5**

Different reading frames of Dpfp5 cDNA sequence (AM230139) match to several predicted proteins from the ZM genome. The most significant match is with reading frame 532 (E= 3e-44) which matches to DPMN_159966. The only predicted protein with a signal peptide was DPMN_094983, which has matched to Dpfp5-531, the same Dpfp5 sequence reported in 2013 [1]. Dpfp5 is likely DPMN_094983, due to the presence of signal peptide, matches to previously reported sequence, and MW more similar to that previously reported.

[1] Gantayet, A., Ohana, L., and Sone, E.D. 2013. Byssal proteins of the freshwater zebra mussel, *Dreissena polymorpha*. *Biofouling* **29(1)**: 77-85. DOI: 10.1080/08927014.2012.746672.

DPMN_094983:

MFSTVTIVLLVSGCATATISQYNYWPGGKGLYNNYWNRPQQSYPTWRLYDPCDKVYCYPIYCRYGQYTPQGECCPQCTPGSYRPGSWNNVGQQGNAVSGLGNNVGSQGNAVSGGWNNVGSQGNSVSGGWNNVGSQGNSVSGGWNNVGSQGNAVSGGWNNVGSQGNSVSGGWNNVGSQGNSVSGGWNHVGSQGNAVSG MW: 18392.52

DPMN_159966:

MSSATKKNTLGKVITLASKGMMWTGMRISLAGKVMPWAGREMTLVSKRMPLVGQGIPWDGKGITWAGKLLEVQIKWIPFGNDTYLT

Dpfp5_AM230139-531:

GRGNSISSGRPGRYNSWPPKPNQPQQPQQPQQPPQPPRYPQPSYPAYPPQQSYPAYPPKQSYPTYPPKQSYPAYPPKQSYPTNPPYNPCDAVYCRPIYCNYGQYTPQGECCPQCNPGTYLPEKWSWKGNNVVGDQEKYVGEGNNVGEQRNDVDGNENIVGGQSNAVGGKGNDVGEQKNAVGGSGNTVGWQGNNVGG*TP*STSAATTLITSEF

Matches to DPMN_094983 (E= 3e-34)

Query 76	PKQSYPTNPPYNPCDAVYCRPIYCNYGQYTPQGECCPQCNPGTYLPEKWSWKGNNVVGDQ 135

P+QSYPT	Y+PCD VYC PIYC YGQYTPQGECCPQC PG+Y P W	N VG Q

Sbjct 39	PQQSYPTWRLYDPCDKVYCYPIYCRYGQYTPQGECCPQCTPGSYRPGSW-----NNVGQQ 93

Query 136 EKYV-GEGNNVGEQRNDVDGNENIVGGQSNAVGGKGNDVGEQKNAVGGSGNTVGWQGNNV 194

V G GNNVG Q N V G N VG Q N+V G N+VG Q N+V G N VG QGN V

Sbjct 94	GNAVSGLGNNVGSQGNAVSGGWNNVGSQGNSVSGGWNNVGSQGNSVSGGWNNVGSQGNAV 153

Query 195 GG 196

G

Sbjct 154 SG 155

DPMN_094983:

MFSTVTIVLLVSGCATATISQYNYWPGGKGLYNNYWNRPQQSYPTWRLYDPCDKVYCYPIYCRYGQYTPQGECCPQCTPGSYRPGSWNNVGQQGNAVSGLGNNVGSQGNAVSGGWNNVGSQGNSVSGGWNNVGSQGNSVSGGWNNVGSQGNAVSGGWNNVGSQGNSVSGGWNNVGSQGNSVSGGWNHVGSQGNAVSG

Dpfp5_AM230139-532:

AAGIRFRAAARAGTTAGHRNQTNHNNHSSHNNHRSLLVILNHHIRHILHNNHIRHILQNSHIRHILQNNHIRHILQNNHIRQTLRIILVTLCIAVRSIAIMDNTHLKANAARSATQAPICQKSGAGRGIMSSATKKNTLGKVITLASKGMMWTGMKISLAGKVMPWAGREMTLASKRMPLVGQEIPWDGKGITWAGKLLEVPRPRPR*SLVN

Matches to DPMN_159966 (E= 3e-44)

Query 130 MSSATKKNTLGKVITLASKGMMWTGMKISLAGKVMPWAGREMTLASKRMPLVGQEIPWDG 189

MSSATKKNTLGKVITLASKGMMWTGM+ISLAGKVMPWAGREMTL SKRMPLVGQ IPWDG

Sbjct 1	MSSATKKNTLGKVITLASKGMMWTGMRISLAGKVMPWAGREMTLVSKRMPLVGQGIPWDG 60

Query 190 KGITWAGKLLEV 201

KGITWAGKLLEV

Sbjct 61	KGITWAGKLLEV 72

DPMN_159966:

MSSATKKNTLGKVITLASKGMMWTGMRISLAGKVMPWAGREMTLVSKRMPLVGQGIPWDGKGITWAGKLLEVQIKWIPFGNDTYLT

Dpfp5_AM230139-351:

EFTSD*RGRGRGTSRSLPAHVIPLPSHGIS*PTNGILLLANVISLPAHGITLPANDIFIPVHIIPLLANVITFPNVFFLVADDIIPLPAPLFWQIGAWVALRAAFALRCVLSIIAIDRTAIHSVTRIIRRVCRI*LFWRICRI*LFWRICRI*LFWRICRI*LLWRICRI*WLRITRRLRWLLWLLWLLWLVWFRWPAVVPARAAARNRIPAA

Matches to DPMN_160100 (E= 3e-08)

Query 16	SLPAHVIPLPSHGIS*PTNGILLLANVISLPAHGITLPANDIFIPVHIIPLLANVITFPN 75

S PAH IPL +H I P +GI L A+VI LPAHGI L A I +P H IPL A+VI P

Sbjct 647 STPAHGIPLRAHVIQLPAHGIPLRAHVIQLPAHGIPLRARVIQLPAHGIPLRAHVIQPPL 706

Query 76	VFF 78

F

Sbjct 707 TVF 709

DPMN_160100:

MFPSLRSFPSELTLFYLRIPLRANVILSPHSLASPPHVILSMQSLASPRYSIYAFPSEPTLFYRRIPLLTNQNVNPRTRYSLASPRYYISAFPCEPTLFYLRIPLQANQNQAHVILSPHSLASPRYSISAFPSEPTLFYLRIPFRSNVIISPHSLESQPINVILSPHSLASQLELLLPEHGIPLRAHIIQPTAYGIPLQTHSFPSEATLFYLRIPLRAHVIQLPAHGIPLRAHNVNHPNTVFPCEPTLFCLRIPLRVHVILSPHSLFYLRIPLRVHVILSPHSLANQPERHAPEHGIPLRAHVILSPHSLRAHVIQSPHSLASPRYSISAFPSEPTLLYLRIPFRSNVIISPHSLERQRYSISEFPSDPTLFYLGIPLRARVILSPHSLASQPELQPPEHGIPLRTHVIQPTAYGIPLQTHSIPSEATLFYLRIPLRAHVIQLPAHGIPLRAHVIQFLAHGIPLRARVMQLPAHGIPLRARVMQLPAHGIPLRAHVIQFLAHGIPLRARVMQLPAHGIPLRAHVMQFLAHGIPLRARVMQLPAHGIPLRAHVFQLPAHGIPLRAHVIQLPAQAHIIQLPPHGIPLRAHVIQLPSHGIPLRAHVIQLTRSRYSIASPRYSTPRSRYSLASPRYSTPRSRYSLASPRYSTPAHGIPLRAHVIQLPAHGIPLRAHVIQLPAHGIPLRARVIQLPAHGIPLRAHVIQPPLTVFPCEPTLFNPRSRYSIASPRYSTPAHGIPLRANVIQLPAHGIPLRAHVIQPPLTTKRSMKALLSFWC

Dpfp5_AM230139-353:

IH**LAWSRPRYFKEFTRPRYSLAIPRYFLTHQRHSFARQRHFPSRPRHYFARQRYFHSRPHHSFARQRYYLPQRIFLGRRRHYSPSSSTFLANRCLGCTAGSIRLEVCIVHNCNRSDGNTQRHKDYTEGLSDMIVLEDMPDMIVLEDMSDMTVLEDMPDMIVVEDMPDMMVEDNEEAAVVVVAAVVVVVGLVSVASCCTCPGGRSKSNSRG

Matches to DPMN_159965 (E= 4e-21)

Query 134 MIVLEDMPDMIVLEDMSDMTVLEDMPDMIVVEDMPDMMVEDNEEAAVVVVAAVVVVVGLV 193

MIVLEDMPDMIVLEDMSDM VLEDMPDMI VEDMPDMMVEDNEEA VVVVAAVVVVVGLV

Sbjct 1	MIVLEDMPDMIVLEDMSDMIVLEDMPDMIAVEDMPDMMVEDNEEAVVVVVAAVVVVVGLV 60

Query 194 SVASCCT 200

SVASCCT

Sbjct 61	SVASCCT 67

DPMN_159965:

MIVLEDMPDMIVLEDMSDMIVLEDMPDMIAVEDMPDMMVEDNEEAVVVVVAAVVVVVGLVSVASCCTDLSVV

**Dpfp6**

We didn’t initially find any Dpfp6 in the predicted protein sequences. We had weak matches to DPMN_151689, DPMN_157295, and DPMN_186587. None of the conserved regions reported in 2014 [1] (YDYDGPYDK, NPGPYDYDGPYDK, KPDPYGTDWQYDKK, KPGPYDYDGPYDK) were found in any proteins predicted by the ZM genome. This could be due to filtering of low-complexity regions by BLAST.

Most likely, as previously reported Dpfp6 is part of a longer Dpfp1 sequence. More detailed examination of the Dpfp1/6 genomic locus identified unannotated exons corresponding to the previously published Dpfp6 sequence, **see further detail beginning Page 21**.

[1] Gantayet, A., Rees, D.J., and Sone, E.D. 2014. Novel Proteins Identified in the Insoluble Byssal Matrix of the Freshwater Zebra Mussel. *Marine Biotechnology* **16**: 144-155. DOI: 10.1007/s10126-013-9537-9.

Dpfp6a_AM229723-531

GRGNSLVISVVAAEVHRSHMDLVLYHKVHRSHMDLVSYHMVHRSHMDLVFYHMVHCSHMDLDFYHMVHRSHMDLVFYPLIYRGHMDLFSYHIANRCHMDLVFYHMVHRSHMDLGFYHMVHRSRTCPGGRSKSLVNSRG

DPMN_151589 2e-08

Query 15	VHRSHMDLVLYHKVHRSHMDLVSYHMVHRSHMDLVFYHMVHCSHMDLDFYHMVHRSHMDL 74

+ R +DLV+	+ R +DLV	+ R +DL+	+	+DL	++ R +DL

Sbjct 19	ICRRCVDLVIIQFICRRCVDLVILQFICRRCVDLMIMQFICRRCVDLVIIQLICRRCVDL 78

Query 75	VFYPLIYRGHMDLFSYH-IANRCHMDLVFYHMVHRSHMDLGFYHMVHR 121

V	I R +DL	I RC +DL+	+ R ++DL	+ R

Sbjct 79	VILQFICRRCVDLMIMQFICRRC-LDLLILQFICRRYVDLVIMQFICR 125

Dpfp6a_AM229723-351

GRGNSLVISSGRPGRYDYDGPYDKNPGPYDYDGPYDKKPDPYGTDWQYDKKTGPYVPDKSEDKKPGPYDYDGPYDKNPGPYDYNGPYDKKPGPYDYDGPYDKKPGPYDYDGPYDIKPGPYDYDVPRPRPR*SLVNSRG

DPMN_157295 1e-09

Query 18	YDGPYDKNPGPYDYDGPYDKKPDPYGTDWQYDKKTGPYVPDKSEDKKPGPYDYDGPYDKN 77

+D YD+	P +D YD+ DP	D YD+	P V D	D+	P +D YD+

Sbjct 92	WDALYDEAADPAVWDALYDEAADPAVWDALYDEAADPAVWDALYDEAADPAVWDALYDEA 151

Query 78	PGPYDYNGPYDKKPGPYDYDGPYDKKPGPYDYDGPYDIKPGPYDYD 123

P ++ YD+	P +D YD+	P +D YD	P +D

Sbjct 152 ADPAVWDALYDEAADPAVWDALYDEAADPAVWDALYDEAADPAVWD 197

Dpfp6b_AM229736-531

GRGNSISVVAAEVHRSHMDLVLYHKVHRSHMDLVSYHMVHRSHMDLVFYHMVHCSHMDLDFYHMVHRSHMDLVFYPLIYRGHMDLFSYHIANRCHMDLVFYHMVHRSHMDLGFYHMVHRSHMDLVFYPQRDRTSAATTLITSEF

DPMN_151589 3e-10

Query 13	VHRSHMDLVLYHKVHRSHMDLVSYHMVHRSHMDLVFYHMVHCSHMDLDFYHMVHRSHMDL 72

+ R +DL++	+ R +DLV	+ R +DLV	+	+DL	+ R +DL

Sbjct 6	ICRRCVDLMIMQFICRRCVDLVIIQFICRRCVDLVILQFICRRCVDLMIMQFICRRCVDL 65

Query 73	VFYPLIYRGHMDLFSYH-IANRCHMDLVFYHMVHRSHMDLGFYHMVHRSHMDLV 125

V	LI R +DL	I RC +DL+	+ R +DL	+ R ++DLV

Sbjct 66	VIIQLICRRCVDLVILQFICRRC-VDLMIMQFICRRCLDLLILQFICRRYVDLV 118

Dpfp6b_AM229736-533

PREFD*RGRGRGTS*SYGPGFIS*GPS*SYGPGFLSYGPS*SYGPGFLSYGPL*SYGPGFLSYGPS*SYGPGFLSSDLSGTYGPVFLSYCQSVPYGSGFLSYGPS*SYGPGFLSYGPS*SYGPGFLSSTGSYLGRDHANH**I

DPMN_186587 8e-07

Query 18	GPGFIS*GPS*SYGPGFLSYGPS*SYGPGFLSYGPL*SYGPGFLSYGPS*SYGPGFLSSD 77

GPG S G + + GPG SYG	+ GP F SYG	+ GPG SYG + + GPG S D

Sbjct 45	GPGLDSYGLASNPGPGLDSYGLDSNPGPWFDSYGLACNPGPGLDSYGLASNPGPGLDSYD 104

Query 78	LSGTYGPVFLSYCQSVPYGSGFLSYGPS*SYGPGFLSYGPS*SYGPGFLS 127

L+	GP	SY +	G G SYG + + GPG SYG + + GPG S

Sbjct 105 LASNPGPGLDSYGLASNPGPGLDSYGLASNPGPGLDSYGLASNPGPGLDS 154

Dpfp6b_AM229736-353

IH**LAWSRPRYDPVEDKKPGPYDYDGPYDKNPGPYDYDGPYDKKPDPYGTDWQYDKKTGPYVPDKSEDKKPGPYDYDGPYDKNPGPYDYNGPYDKKPGPYDYDGPYDKKPGPYDYDGPYDIKPGPYDYDVPRPRPR*SNSRG

DPMN_157295 2e-10

Query 9	RPRYDPVEDKKPGPYDYDGPYDKNPGPYDYDGPYDKKPDPYGTDWQYDKKTGPYVPDKSE 68

+D + D+	P +D YD+	P +D YD+ DP	D YD+	P V D

Sbjct 89	LAVWDALYDEAADPAVWDALYDEAADPAVWDALYDEAADPAVWDALYDEAADPAVWDALY 148

Query 69	DKKPGPYDYDGPYDKNPGPYDYNGPYDKKPGPYDYDGPYDKKPGPYDYDGPYD 121

D+	P +D YD+	P ++ YD+	P +D YD+	P +D YD

Sbjct 149 DEAADPAVWDALYDEAADPAVWDALYDEAADPAVWDALYDEAADPAVWDALYD 201

Dpfp6c_AM229737-531

GRGNSISVVAAVVHRSHMDLVLYHKVHRSHMDLVSYHMVHRSHMDLVFYHMVHCSHMDLDFYHMVHRSHMDLVFYPLIYRGHMDLFSYHIANRCHMDLVFYHMVHRSHMDLGFYHMVHRSHMDLVFYPQRDRTCPGGRSKSLVN

DPMN_151589 4e-10

Query 13	VHRSHMDLVLYHKVHRSHMDLVSYHMVHRSHMDLVFYHMVHCSHMDLDFYHMVHRSHMDL 72

+ R +DL++	+ R +DLV	+ R +DLV	+	+DL	+ R +DL

Sbjct 6	ICRRCVDLMIMQFICRRCVDLVIIQFICRRCVDLVILQFICRRCVDLMIMQFICRRCVDL 65

Query 73	VFYPLIYRGHMDLFSYH-IANRCHMDLVFYHMVHRSHMDLGFYHMVHRSHMDLV 125

V	LI R +DL	I RC +DL+	+ R +DL	+ R ++DLV

Sbjct 66	VIIQLICRRCVDLVILQFICRRC-VDLMIMQFICRRCLDLLILQFICRRYVDLV 118

Dpfp6c_AM229737-533

PREFD*RGRGRGTS*SYGPGFIS*GPS*SYGPGFLSYGPS*SYGPGFLSYGPL*SYGPGFLSYGPS*SYGPGFLSSDLSGTYGPVFLSYCQSVPYGSGFLSYGPS*SYGPGFLSYGPS*SYGPGFLSSTGSYLPGRPLEITSEF

DPMN_186587 6e-07

Query 18	GPGFIS*GPS*SYGPGFLSYGPS*SYGPGFLSYGPL*SYGPGFLSYGPS*SYGPGFLSSD 77

GPG S G + + GPG SYG	+ GP F SYG	+ GPG SYG + + GPG S D

Sbjct 45	GPGLDSYGLASNPGPGLDSYGLDSNPGPWFDSYGLACNPGPGLDSYGLASNPGPGLDSYD 104

Query 78	LSGTYGPVFLSYCQSVPYGSGFLSYGPS*SYGPGFLSYGPS*SYGPGFLS 127

L+	GP	SY +	G G SYG + + GPG SYG + + GPG S

Sbjct 105 LASNPGPGLDSYGLASNPGPGLDSYGLASNPGPGLDSYGLASNPGPGLDS 154

Dpfp6c_AM229737-352

NSLVISSGRPGRYDPVEDKKPGPYDYDGPYDKNPGPYDYDGPYDKKPDPYGTDWQYDKKTGPYVPDKSEDKKPGPYDYDGPYDKNPGPYDYNGPYDKKPGPYDYDGPYDKKPGPYDYDGPYDIKPGPYDYDVPRPRPR*SNSRG

DPMN_157295 3e-11

Query 7	SGRPGRYDPVEDKKPGPYDYDGPYDKNPGPYDYDGPYDKKPDPYGTDWQYDKKTGPYVPD 66

+ P +D + D+	P +D YD+	P +D YD+ DP	D YD+	V D

Sbjct 34	AADPAVWDALYDEAADPAVWDALYDEAADPAVWDALYDEAADPAVWDALYDEAADLAVWD 93

Query 67	KSEDKKPGPYDYDGPYDKNPGPYDYNGPYDKKPGPYDYDGPYDKKPGPYDYDGPYDIKPG 126

D+	P +D YD+	P ++ YD+	P +D YD+	P +D YD

Sbjct 94	ALYDEAADPAVWDALYDEAADPAVWDALYDEAADPAVWDALYDEAADPAVWDALYDEAAD 153

Query 127 PYDYD 131

P +D

Sbjct 154 PAVWD 158

These are the Dpfp6 cDNA translated reading frames that correspond to the reported protein sequence (highlighted is the region that is reported):

>Dpfp6a_AM229723-351

GRGNSLVISSGRPGRYDYDGPYDKNPGPYDYDGPYDKKPDPYGTDWQYDKKTGPYVPDKSEDKKPGPYDYDGPYDKNPGPYDYNGPYDKKPGPYDYDGPYDKKPGPYDYDGPYDIKPGPYDYDVPRPRPR*SLVNSRG

>Dpfp6b_AM229736-353

IH**LAWSRPRYDPVEDKKPGPYDYDGPYDKNPGPYDYDGPYDKKPDPYGTDWQYDKKTGPYVPDKSEDKKPGPYDYDGPYDKNPGPYDYNGPYDKKPGPYDYDGPYDKKPGPYDYDGPYDIKPGPYDYDVPRPRPR*SNSRG

>Dpfp6c_AM229737-352

NSLVISSGRPGRYDPVEDKKPGPYDYDGPYDKNPGPYDYDGPYDKKPDPYGTDWQYDKKTGPYVPDKSEDKKPGPYDYDGPYDKNPGPYDYNGPYDKKPGPYDYDGPYDKKPGPYDYDGPYDIKPGPYDYDVPRPRPR*SNSRG

Gantayet 2014 reports that some peptide motifs of the Dpfp6 sequence match to Dpfp1. BLASTp of Dpfp1 against Dpfp6 confirms this:

blastp -db Dpfp6_translated.fasta -query C:\Users\Anjo\Desktop\ZM_Genome\Dpfp6\Dpfp1.fasta -out C:\Users\Anjo\Desktop\ZM_Genome\Dpfp6\Dpfp1onDpfp6

[Dpfp6a_AM229723-351 170 5e-55]

[Dpfp6b_AM229736-353 167 1e-53]

[Dpfp6c_AM229737-352 167 1e-53]

The Dpfp1 cDNA reading frame that corresponds to the reported sequence is 531:

ILQSINQGTWMFSVVSFCLLAAGFGSSLGGSSDWTEKTSQSTIPTISGWSFFTTKSPLNPTLFTTKRPEYVTLSPVYPTKIPNYTTKPPVYPTKVPEYPTKDPTYPTFKTPEYPTKVPEYPTKVPTYPTFQTPEYPTPTKYPVYPSQSPAYPTQYPEYPSQYPVYPDQYPVYPNQYPVKQDHDPVYPPRSPLYGWRRPVYPKKTPVYPYLPLYPGYQPEYHRRPPVYPPVYPYDPVEDKKPGPYDYDGPYDKNPGPYDYDGPYNKKPNPYGTDWQYDKKTGPYVPIKPDDKKPNPYGTDWQYDKKTGPYVPDKSEDKKPGPYDYDGPYDKNPGPYDSDGPYNKKPGPYDYDGPYDKNPGPYDYNGPYDKKPGPYDYDGPYDIKPGPYDYDVPYDKKPDPYDTDGPYDKKTGPYVPDKPDDKKTDPYVPDVPLEPPGPLGK*SCQQDKQGIDVELVQMTCISIRIDTLLLC*HTNK*IRS

Dpfp1 corresponds to predicted protein [DPMN_160000 423 5e-148] and [DPMN_160107 263 2e-85]:

>DPMN_160000

MFSVVSFCLLAAGFGSSLGGSSDWTEKTSQSTIPTISGWSFFTTKSPLNPTLFTTKRPEYVTLSPVYPTKIPNYTTKPPVYPTKVPEYPTKDPTYPTFKTPEYPTKVPEYPTKVPTYPTFQTPEYPTPTKYPVYPSQSPAYPTQYPEYPSQYPVYPDQYPVYPNQYPVKQDHDPVYPPRSPLYGWRRPVYPKKTPVYPYLPLYPGYQPEYHRRPPVYPPVYPYDPVGKCDGEYCSSLFYFNGLT

>DPMN_160107

MFSAVTFLLLAASFGSSLSGASPDVTEKPSNQPTLPTFSGWSGFPTKSPLNPILYTLFATKRPEYVTLSPVYSTKVPEYPTKVPTYQTFKTPEYPTKVPEYPTKIPTYPTFKTPEYPTPTKYPVYPSQSPAYPTQYPEYPSPYSEYPSKYPVYPDQYPVYPNQYPFDLAQYPVYPNQYPVEQDQYPPRSPLYRGRRPVYPNEPPVYPDLPLYPGYQPGYHRRPPVYPPVYPYDPVGKCDGEYCSSLFYFNGLT

Mapping these sequences onto the Dpfp1 translated cDNA sequence:

DPMN_160000

ILQSINQGTWMFSVVSFCLLAAGFGSSLGGSSDWTEKTSQSTIPTISGWSFFTTKSPLNPTLFTTKRPEYVTLSPVYPTKIPNYTTKPPVYPTKVPEYPTKDPTYPTFKTPEYPTKVPEYPTKVPTYPTFQTPEYPTPTKYPVYPSQSPAYPTQYPEYPSQYPVYPDQYPVYPNQYPVKQDHDPVYPPRSPLYGWRRPVYPKKTPVYPYLPLYPGYQPEYHRRPPVYPPVYPYDPVEDKKPGPYDYDGPYDKNPGPYDYDGPYNKKPNPYGTDWQYDKKTGPYVPIKPDDKKPNPYGTDWQYDKKTGPYVPDKSEDKKPGPYDYDGPYDKNPGPYDSDGPYNKKPGPYDYDGPYDKNPGPYDYNGPYDKKPGPYDYDGPYDIKPGPYDYDVPYDKKPDPYDTDGPYDKKTGPYVPDKPDDKKTDPYVPDVPLEPPGPLGK*SCQQDKQGIDVELVQMTCISIRIDTLLLC*HTNK*IRS

DPMN_160107

ILQSINQGTWMFSVVSFCLLAAGFGSSLGGSSDWTEKTSQSTIPTISGWSFFTTKSPLNPTLFTTKRPEYVTLSPVYPTKIPNYTTKPPVYPTKVPEYPTKDPTYPTFKTPEYPTKVPEYPTKVPTYPTFQTPEYPTPTKYPVYPSQSPAYPTQYPEYPSQYPVYPDQYPVYPNQYPVKQDHDPVYPPRSPLYGWRRPVYPKKTPVYPYLPLYPGYQPEYHRRPPVYPPVYPYDPVEDKKPGPYDYDGPYDKNPGPYDYDGPYNKKPNPYGTDWQYDKKTGPYVPIKPDDKKPNPYGTDWQYDKKTGPYVPDKSEDKKPGPYDYDGPYDKNPGPYDSDGPYNKKPGPYDYDGPYDKNPGPYDYNGPYDKKPGPYDYDGPYDIKPGPYDYDVPYDKKPDPYDTDGPYDKKTGPYVPDKPDDKKTDPYVPDVPLEPPGPLGK*SCQQDKQGIDVELVQMTCISIRIDTLLLC*HTNK*IRS

Dpfp6a-351

ILQSINQGTWMFSVVSFCLLAAGFGSSLGGSSDWTEKTSQSTIPTISGWSFFTTKSPLNPTLFTTKRPEYVTLSPVYPTKIPNYTTKPPVYPTKVPEYPTKDPTYPTFKTPEYPTKVPEYPTKVPTYPTFQTPEYPTPTKYPVYPSQSPAYPTQYPEYPSQYPVYPDQYPVYPNQYPVKQDHDPVYPPRSPLYGWRRPVYPKKTPVYPYLPLYPGYQPEYHRRPPVYPPVYPYDPVEDKKPGPYDYDGPYDKNPGPYDYDGPYNKKPNPYGTDWQYDKKTGPYVPIKPDDKKPNPYGTDWQYDKKTGPYVPDKSEDKKPGPYDYDGPYDKNPGPYDSDGPYNKKPGPYDYDGPYDKNPGPYDYNGPYDKKPGPYDYDGPYDIKPGPYDYDVPYDKKPDPYDTDGPYDKKTGPYVPDKPDDKKTDPYVPDVPLEPPGPLGK*SCQQDKQGIDVELVQMTCISIRIDTLLLC*HTNK*IRS

Dpfp6b-353

ILQSINQGTWMFSVVSFCLLAAGFGSSLGGSSDWTEKTSQSTIPTISGWSFFTTKSPLNPTLFTTKRPEYVTLSPVYPTKIPNYTTKPPVYPTKVPEYPTKDPTYPTFKTPEYPTKVPEYPTKVPTYPTFQTPEYPTPTKYPVYPSQSPAYPTQYPEYPSQYPVYPDQYPVYPNQYPVKQDHDPVYPPRSPLYGWRRPVYPKKTPVYPYLPLYPGYQPEYHRRPPVYPPVYPYDPVEDKKPGPYDYDGPYDKNPGPYDYDGPYNKKPNPYGTDWQYDKKTGPYVPIKPDDKKPNPYGTDWQYDKKTGPYVPDKSEDKKPGPYDYDGPYDKNPGPYDSDGPYNKKPGPYDYDGPYDKNPGPYDYNGPYDKKPGPYDYDGPYDIKPGPYDYDVPYDKKPDPYDTDGPYDKKTGPYVPDKPDDKKTDPYVPDVPLEPPGPLGK*SCQQDKQGIDVELVQMTCISIRIDTLLLC*HTNK*IRS

Dpfp6c-352

ILQSINQGTWMFSVVSFCLLAAGFGSSLGGSSDWTEKTSQSTIPTISGWSFFTTKSPLNPTLFTTKRPEYVTLSPVYPTKIPNYTTKPPVYPTKVPEYPTKDPTYPTFKTPEYPTKVPEYPTKVPTYPTFQTPEYPTPTKYPVYPSQSPAYPTQYPEYPSQYPVYPDQYPVYPNQYPVKQDHDPVYPPRSPLYGWRRPVYPKKTPVYPYLPLYPGYQPEYHRRPPVYPPVYPYDPVEDKKPGPYDYDGPYDKNPGPYDYDGPYNKKPNPYGTDWQYDKKTGPYVPIKPDDKKPNPYGTDWQYDKKTGPYVPDKSEDKKPGPYDYDGPYDKNPGPYDSDGPYNKKPGPYDYDGPYDKNPGPYDYNGPYDKKPGPYDYDGPYDIKPGPYDYDVPYDKKPDPYDTDGPYDKKTGPYVPDKPDDKKTDPYVPDVPLEPPGPLGK*SCQQDKQGIDVELVQMTCISIRIDTLLLC*HTNK*IRS

Dpfp6 and the DPMN sequences map to distinct components of the Dpfp1 cDNA sequence.

ZM Genome Predicted Protein Sequence | Dpfp6

ILQSINQGTWMFSVVSFCLLAAGFGSSLGGSSDWTEKTSQSTIPTISGWSFFTTKSPLNPTLFTTKRPEYVTLSPVYPTKIPNYTTKPPVYPTKVPEYPTKDPTYPTFKTPEYPTKVPEYPTKVPTYPTFQTPEYPTPTKYPVYPSQSPAYPTQYPEYPSQYPVYPDQYPVYPNQYPVKQDHDPVYPPRSPLYGWRRPVYPKKTPVYPYLPLYPGYQPEYHRRPPVYPPVYPYDPVEDKKPGPYDYDGPYDKNPGPYDYDGPYNKKPNPYGTDWQYDKKTGPYVPIKPDDKKPNPYGTDWQYDKKTGPYVPDKSEDKKPGPYDYDGPYDKNPGPYDSDGPYNKKPGPYDYDGPYDKNPGPYDYNGPYDKKPGPYDYDGPYDIKPGPYDYDVPYDKKPDPYDTDGPYDKKTGPYVPDKPDDKKTDPYVPDVPLEPPGPLGK*SCQQDKQGIDVELVQMTCISIRIDTLLLC*HTNK*IRS

MFSVVSFCLLAAGFGSSLGGSSDWTEKTSQSTIPTISGWSFFTTKSPLNPTLFTTKRPEYVTLSPVYPTKIPNYTTKPPVYPTKVPEYPTKDPTYPTFKTPEYPTKVPEYPTKVPTYPTFQTPEYPTPTKYPVYPSQSPAYPTQYPEYPSQYPVYPDQYPVYPNQYPVKQDHDPVYPPRSPLYGWRRPVYPKKTPVYPYLPLYPGYQPEYHRRPPVYPPVYPYDPVGKCDGEYCSSLFYFNGLT

BLASTn Dpfp1 cDNA and Dpfp6 cDNA against the ZM genome.

*The yellow-highlighted bits match to the genome. The blue-highlighted bit matches to Dpfp6.* The parts of the Dpfp1 cDNA that match to the ZM genome are not the same parts that match to Dpfp6. The part of Dpfp6 that matches to the ZM genome is not in the part of Dpfp1 that matches to the ZM genome. The section of the Dpfp1 cDNA that matches to Dpfp6 is not found in the ZM genome, **so where does this cDNA come from**?

>Dpfp1_AAF265353

ATACTTCAGAGCATAAACCAAGGTACTTGGATGTTCTCCGTGGTATCATTCTGTCTTCTTGCGGCGGGCTTCGGCTCGTCATTGGGTGGGAGCTCTGATTGGACAGAAAAAACCTCACAATCAACTATACCGACAATTAGCGGATGGTCTTTTTTTACAACTAAATCTCCGTTAAATCCAACTCTATTTACAACGAAACGTCCGGAATATGTAACTCTATCCCCGGTATATCCAACTAAAATTCCGAACTATACAACAAAACCTCCGGTATATCCAACTAAAGTTCCGGAATATCCAACGAAAGATCCGACATATCCAACTTTCAAAACTCCGGAATATCCAACAAAAGTTCCGGAATATCCAACGAAAGTTCCGACATATCCAACTTTCCAAACTCCGGAATATCCCACTCCTACAAAATATCCAGTATATCCATCTCAATCTCCTGCATATCCTACTCAGTACCCTGAATATCCGTCTCAATATCCTGTATATCCCGATCAGTATCCAGTATATCCGAATCAGTATCCGGTAAAACAAGATCACGATCCAGTGTATCCACCACGATCACCGTTGTATGGATGGAGACGTCCGGTATATCCAAAAAAAACTCCGGTATACCCATATCTACCGCTATATCCGGGTTATCAACCAGAATATCACCGACGCCCTCCAGTATATCCTCCGGTGTATCCGTACGATCCCGTTGAGGATAAAAAACCAGGTCCATATGACTACGATGGACCATATGATAAAAACCCAGGTCCATATGACTACGATGGACCATATAATAAAAAACCAAATCCATATGGCACCGATTGGCAATATGATAAGAAAACAGGTCCATATGTCCCCATTAAACCAGATGATAAAAAACCAAATCCATATGGCACCGATTGGCAATATGATAAGAAAACAGGTCCATATGTCCCCGATAAATCAGAGGATAAAAAACCAGGTCCATATGACTACGATGGACCATATGATAAAAACCCAGGTCCATATGACTCCGATGGGCCATATAATAAGAAACCAGGTCCATATGATTACGATGGACCATATGATAAAAATCCAGGTCCATATGACTACAATGGACCATATGATAAAAAACCAGGTCCATATGACTACGATGGACCATATGATATAAAACCAGGTCCATATGACTACGATGTACCTTATGATAAAAAACCAGATCCATATGACACCGATGGGCCATATGATAAGAAAACAGGTCCATATGTCCCCGATAAACCAGATGACAAAAAAACAGATCCATATGTCCCCGATGTTCCATTAGAACCTCCTGGACCATTGGGAAAGTAAAGTTGTCAACAAGACAAGCAAGGCATCGACGTTGAATTAGTACAGATGACATGTATCTCAATACGAATCGACACGTTATTGCTATGTTGACATACTAATAAATAAATACGATCA

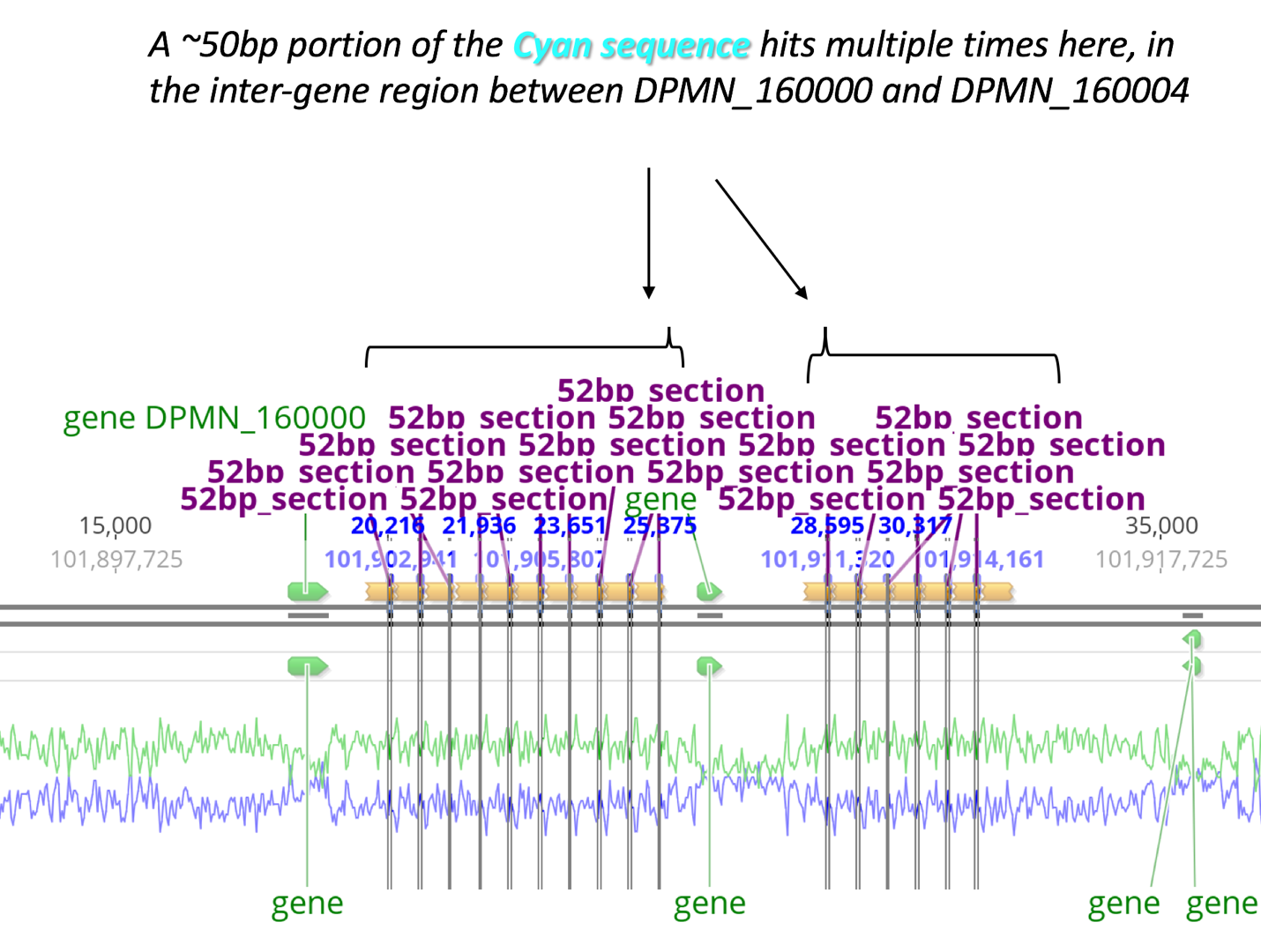

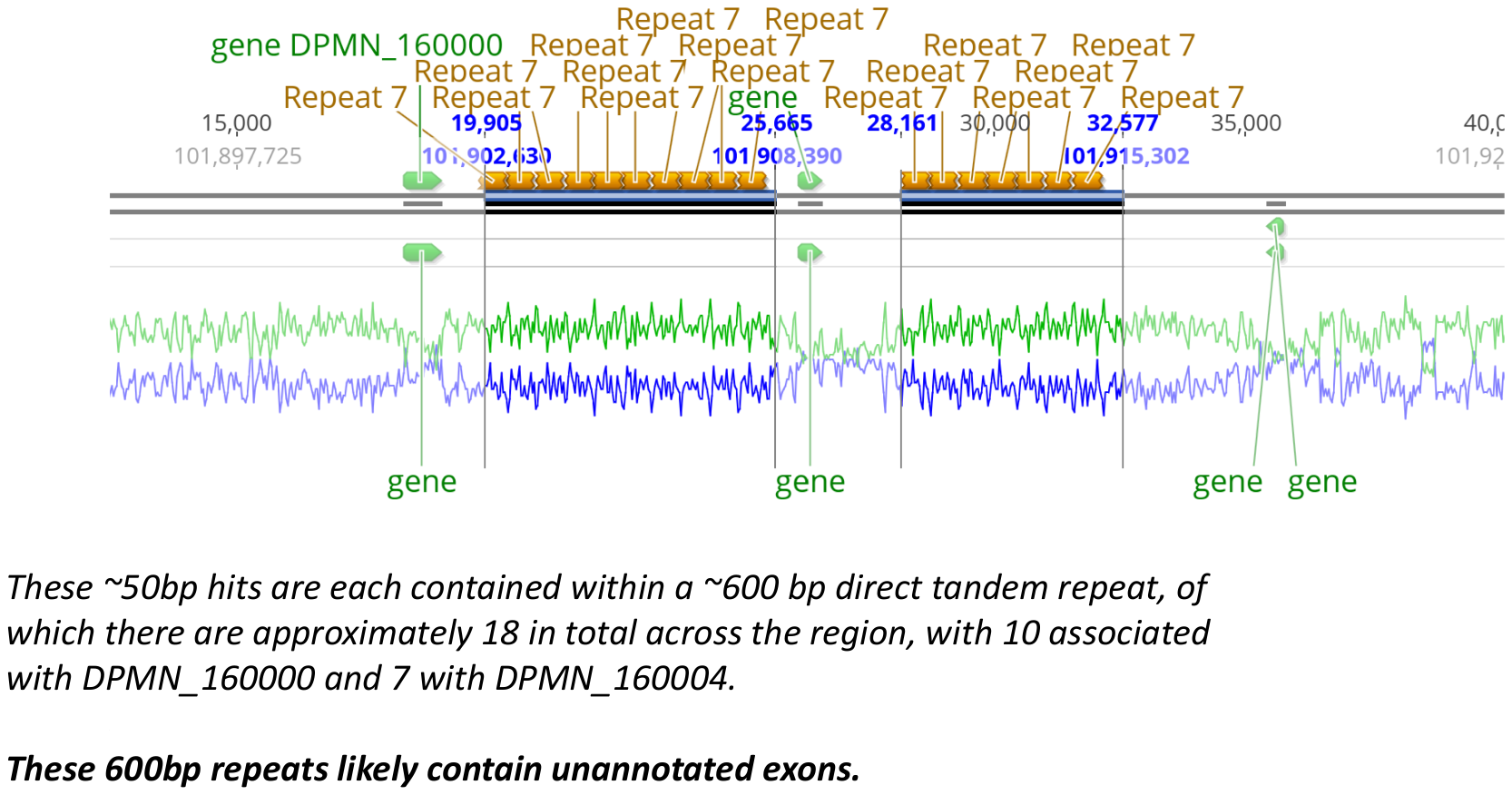

A self-self dotplot of DPMN160000 shows these repetitive sequences clearly:

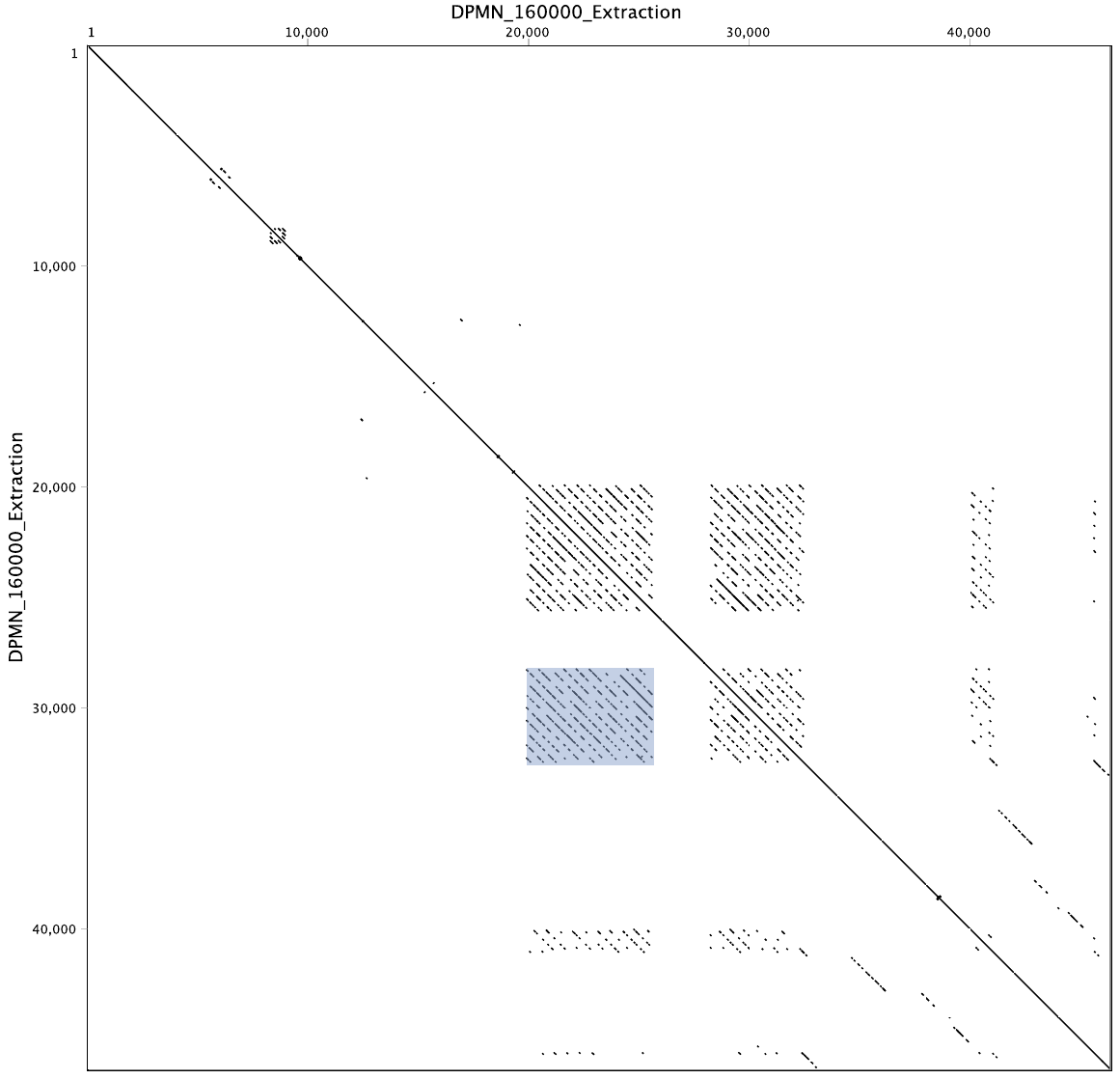

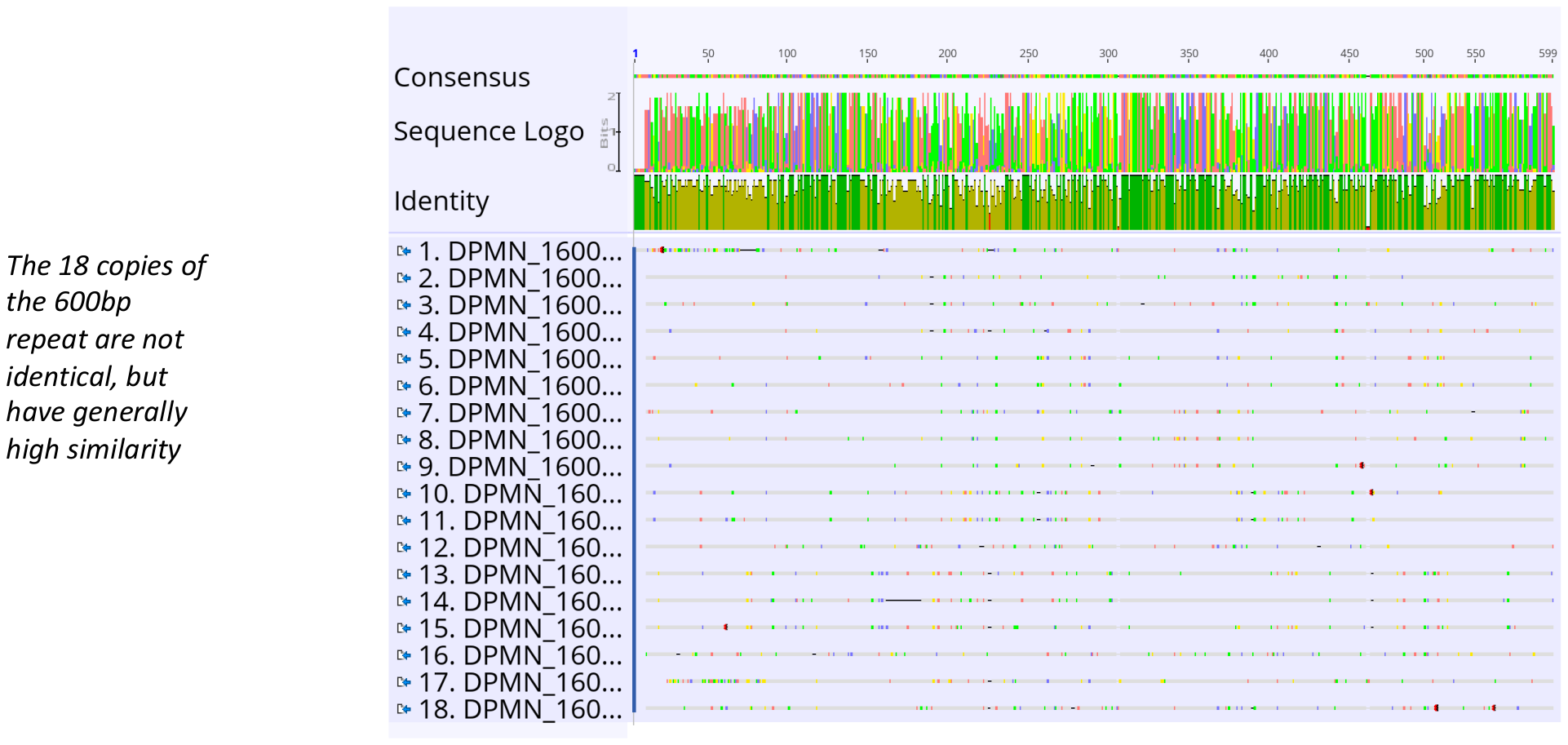

>Dpfp6a_AM229723

GGCCGCGGGAATTCACTAGTGATTAGCGTGGTCGCGGCCGA[GGTACATCGTAGTCATATGGACCTGGTTTTATATCATAAGGTCCATCGTAGTCATATGGACCTGGTTTCTTATCATATGGTCCATCGTAGTCATATGGACCTGGTTTTTTATCATATGGTCCATTGTAGTCATATGGACCTGGATTTTTATCATATGGTCCATCGTAGTCATATGGACCTGGTTTTTTATCCTCTGATTTATCGGGGACATATGGACCTGTTTTCTTATCATATTGCCAATCGGTGCCATATGGATCTGGTTTTTTATCATATGGTCCATCGTAGTCATATGGACCTGGGTTTTTATCATATGGTCCATCG]TAGTCGTACCTGCCCGGGCGGCCGCTCGAAATCACTAGTGAATTCCCGCGGCC

>Dpfp6b_AM229736

GGCCGCGGGAATTCGATTAGCGTGGTCGCGGCCGAGGTACATCGTAGTCATATGGACCTGGTTTTATATCATAAGGTCCATCGTAGTCATATGGACCTGGTTTCTTATCATATGGTCCATCGTAGTCATATGGACCTGGTTTTTTATCATATGGTCCATTGTAGTCATATGGACCTGGATTTTTATCATATGGTCCATCGTAGTCATATGGACCTGGTTTTTTATCCTCTGATTTATCGGGGACATATGGACCTGTTTTCTTATCATATTGCCAATCGGTGCCATATGGATCTGGTTTTTTATCATATGGTCCATCGTAGTCATATGGACCTGGGTTTTTATCATATGGTCCATCGTAGTCATATGGACCTGGTTTTTTATCCTCAACGGGATCGTACCTCGGCCGCGACCACGCTAATCACTAGTGAATTC

>Dpfp6c_AM229737

GGCCGCGGGAATTCGATTAGCGTGGTCGCGGCCGTGGTACATCGTAGTCATATGGACCTGGTTTTATATCATAAGGTCCATCGTAGTCATATGGACCTGGTTTCTTATCATATGGTCCATCGTAGTCATATGGACCTGGTTTTTTATCATATGGTCCATTGTAGTCATATGGACCTGGATTTTTATCATATGGTCCATCGTAGTCATATGGACCTGGTTTTTTATCCTCTGATTTATCGGGGACATATGGACCTGTTTTCTTATCATATTGCCAATCGGTGCCATATGGATCTGGTTTTTTATCATATGGTCCATCGTAGTCATATGGACCTGGGTTTTTATCATATGGTCCATCGTAGTCATATGGACCTGGTTTTTTATCCTCAACGGGATCGTACCTGCCCGGGCGGCCGCTCGAAATCACTAGTGAATTC

Scripts

>makeblastdb -in C:\Users\Anjo\Desktop\ZM_Genome\Dpfp6\Dpfp6_translated.fasta -dbtype prot

>blastp -db Dpfp6_translated.fasta -query C:\Users\Anjo\Desktop\ZM_Genome\Dpfp6\Dpfp6_Gantayet.fasta -out C:\Users\Anjo\Desktop\ZM_Genome\Dpfp6\whichrf

>blastp -db Dpfp6_translated.fasta -query C:\Users\Anjo\Desktop\ZM_Genome\Dpfp6\Dpfp1.fasta -out C:\Users\Anjo\Desktop\ZM_Genome\Dpfp6\Dpfp1onDpfp6

>makeblastdb -in C:\Users\Anjo\Desktop\ZM_Genome\Dpfp6\Dpfp1_translated.fasta -dbtype prot

>blastp -db Dpfp1_translated.fasta -query C:\Users\Anjo\Desktop\ZM_Genome\Dpfp6\Dpfp1_Gantayet.fasta -out C:\Users\Anjo\Desktop\ZM_Genome\Dpfp6\whichdpfp1rf

>makeblastdb -in C:\Users\Anjo\Desktop\ZM_Genome\Dpfp6\Dpfp1-531.fasta -dbtype prot

>blastp -db Dpfp1-531.fasta -query C:\Users\Anjo\Desktop\ZM_Genome\Dpfp6\Dpfp6_translated.fasta -out C:\Users\Anjo\Desktop\ZM_Genome\Dpfp6\Dpfp6onDpfp1

>blastn -db ZM_Asm_Chr.fasta -query C:\Users\Anjo\Desktop\ZM_Genome\Dpfp6\Dpfp1and6_cDNA.fasta -out C:\Users\Anjo\Desktop\ZM_Genome\Dpfp6\cDNAonGenome

>makeblastdb -in C:\Users\Anjo\Desktop\ZM_Genome\Dpfp6\Dpfp6_cDNA.fasta -dbtype nucl

>blastn -db Dpfp6_cDNA.fasta -query C:\Users\Anjo\Desktop\ZM_Genome\Dpfp6\Dpfp1_cDNA.fasta -out C:\Users\Anjo\Desktop\ZM_Genome\Dpfp6\Dpfp1cDNAonDpfp6cDNA

>makeblastdb -in C:\Users\Anjo\Desktop\QMSeq\ReesQMTranscriptome.fasta -dbtype nucl

>blastn -db ReesQMTranscriptome.fasta -query C:\Users\Anjo\Desktop\ZM_Genome\Dpfp6\Dpfp6_cDNA.fasta -out C:\Users\Anjo\Desktop\ZM_Genome\Dpfp6\Dpfp6cDNAonQM

Tblastn -db ReesQMTranscriptome.fasta -query C:\Users\Anjo\Desktop\ZM_Genome\Homology\DpfpNewHomology.fasta -out C:\Users\Anjo\Desktop\ZM_Genome\Homology\DpfpNewHomologyResults

**Dpfp8**

Dpfp8 matches to the C-terminus of DPMN_074797. This sequence does not contain a signal peptide. A significant portion of the full protein sequence (the first 551 residues) has not been found in previous transcriptomics and LC-MS/MS studies.

Dpfp8:

MVNITNINTQITNIVVNQTSIQNNQLTIMNTLNVVETRVNNTEINVVQITQTTTNIQVQVTEITTEITNIYLNINNVTNQVTNVQTTLVNVTQQQISNTNILNNMTVQVTNIDIQLNQVNVNVNNISVFVNQVSVTVINIETELNIVMSNVTQIEHKIISVGATTEELQRHEDIMIENFMLLSVNITNINIAVFNLTNVVNVVDLNVKGTTGEINGIQKNMNQINIYIKDLDSDINDVSGNVFNIQNNINMIMVNITDIDVRIGNVGGGDQGSLDRIFGSFGKFEERITVITSNQNGITDRLGQIDVRINNTQTGIKEISMKVHDIENDMNSVSINTHQLGVNLNTVSTKITNVETNVNLIMVNITNIDIKVGRIGSDQVTTVKTQDSITKNIQVLVDRQSLIQTSQGKMFDRLGYMDNRLNSTQVNLNIMSGSLTHMEGTIDSLYRNVSKINGTVGQVGSGQIDLQYIQELLSHKLDALEGDQTKAFNSQQEIRVQIKNVMEAVNGIVTTISSKNGSCVGGLNFNDFIGKYATKGLQEDLEKKLDYMSAEQRKSQTVQDHMSVRLDNVLKVLGGVATGNKYSSDEIATLVGSTGGGSVNTGGYSKGTYPVPYGTGGVSGYKSGGR

Dpfp8a_AM230242-531

GRGNSISVVAAEVQDHMSVRLDNVLKVLGGVATGNKYSSDEIATLVGSTGGGSVNTGGYSKGTYPVPYGTGGVSGYKSGGR*MDFIGTASAMGDIKWFKLKNDYQLPNDGLKANIISYRLLGRNNAAYLPGRPLEITSEF

Matches to DPMN_074797

Query 10	AAEVQDHMSVRLDNVLKVLGGVATGNKYSSDEIATLVGSTGGGSVNTGGYSKGTYPVPYG 69

+ VQDHMSVRLDNVLKVLGGVATGNKYSSDEIATLVGSTGGGSVNTGGYSKGTYPVPYG

Sbjct 555 SQTVQDHMSVRLDNVLKVLGGVATGNKYSSDEIATLVGSTGGGSVNTGGYSKGTYPVPYG 614

Query 70	TGGVSGYKSGGR 81

TGGVSGYKSGGR

Sbjct 615 TGGVSGYKSGGR 626

MVNITNINTQITNIVVNQTSIQNNQLTIMNTLNVVETRVNNTEINVVQITQTTTNIQVQVTEITTEITNIYLNINNVTNQVTNVQTTLVNVTQQQISNTNILNNMTVQVTNIDIQLNQVNVNVNNISVFVNQVSVTVINIETELNIVMSNVTQIEHKIISVGATTEELQRHEDIMIENFMLLSVNITNINIAVFNLTNVVNVVDLNVKGTTGEINGIQKNMNQINIYIKDLDSDINDVSGNVFNIQNNINMIMVNITDIDVRIGNVGGGDQGSLDRIFGSFGKFEERITVITSNQNGITDRLGQIDVRINNTQTGIKEISMKVHDIENDMNSVSINTHQLGVNLNTVSTKITNVETNVNLIMVNITNIDIKVGRIGSDQVTTVKTQDSITKNIQVLVDRQSLIQTSQGKMFDRLGYMDNRLNSTQVNLNIMSGSLTHMEGTIDSLYRNVSKINGTVGQVGSGQIDLQYIQELLSHKLDALEGDQTKAFNSQQEIRVQIKNVMEAVNGIVTTISSKNGSCVGGLNFNDFIGKYATKGLQEDLEKKLDYMSAEQRKSQTVQDHMSVRLDNVLKVLGGVATGNKYSSDEIATLVGSTGGGSVNTGGYSKGTYPVPYGTGGVSGYKSGGR

Dpfp8b_AM230362-533

PREFDFERPPGQVQDHMSVRLDNVLKVLGGVATGNKYSSDEIATLVGSTGGGSVNTGGYSKGTYPVPYGTGGVSGYKSGGR*MDFIGTASAMGDIKWFKLKNDYQLPNDGLKANIISYRLLGRNNAAYLGRDHANH**I

Matches to DPMN_074797

Query 7	ERPPGQVQDHMSVRLDNVLKVLGGVATGNKYSSDEIATLVGSTGGGSVNTGGYSKGTYPV 66

+R	VQDHMSVRLDNVLKVLGGVATGNKYSSDEIATLVGSTGGGSVNTGGYSKGTYPV

Sbjct 552 QRKSQTVQDHMSVRLDNVLKVLGGVATGNKYSSDEIATLVGSTGGGSVNTGGYSKGTYPV 611

Query 67	PYGTGGVSGYKSGGR 81

PYGTGGVSGYKSGGR

Sbjct 612 PYGTGGVSGYKSGGR 626

MVNITNINTQITNIVVNQTSIQNNQLTIMNTLNVVETRVNNTEINVVQITQTTTNIQVQVTEITTEITNIYLNINNVTNQVTNVQTTLVNVTQQQISNTNILNNMTVQVTNIDIQLNQVNVNVNNISVFVNQVSVTVINIETELNIVMSNVTQIEHKIISVGATTEELQRHEDIMIENFMLLSVNITNINIAVFNLTNVVNVVDLNVKGTTGEINGIQKNMNQINIYIKDLDSDINDVSGNVFNIQNNINMIMVNITDIDVRIGNVGGGDQGSLDRIFGSFGKFEERITVITSNQNGITDRLGQIDVRINNTQTGIKEISMKVHDIENDMNSVSINTHQLGVNLNTVSTKITNVETNVNLIMVNITNIDIKVGRIGSDQVTTVKTQDSITKNIQVLVDRQSLIQTSQGKMFDRLGYMDNRLNSTQVNLNIMSGSLTHMEGTIDSLYRNVSKINGTVGQVGSGQIDLQYIQELLSHKLDALEGDQTKAFNSQQEIRVQIKNVMEAVNGIVTTISSKNGSCVGGLNFNDFIGKYATKGLQEDLEKKLDYMSAEQRKSQTVQDHMSVRLDNVLKVLGGVATGNKYSSDEIATLVGSTGGGSVNTGGYSKGTYPVPYGTGGVSGYKSGGR

**Mapping Byssal Proteins to Predicted Proteins**

Dpfp1 = DPMN_160000

MFSVVSFCLLAAGFGSSLGGSSDWTEKTSQSTIPTISGWSFFTTKSPLNPTLFTTKRPEYVTLSPVYPTKIPNYTTKPPVYPTKVPEYPTKDPTYPTFKTPEYPTKVPEYPTKVPTYPTFQTPEYPTPTKYPVYPSQSPAYPTQYPEYPSQYPVYPDQYPVYPNQYPVKQDHDPVYPPRSPLYGWRRPVYPKKTPVYPYLPLYPGYQPEYHRRPPVYPPVYPYDPVGKCDGEYCSSLFYFNGLT

Dpfp2 = Dpfp12 = Dpfp122 = DPMN_127665

MFSAAALLLLVSFYGTASGQYWNSYRPYPVYPPKQTYPSYPDKKYPSYPEKTYQEKTYPGYPPKQAYPVYPEKTYPEKTYPAYPTKKSYPEYPEKTYTKKTYEAYPTKDSYTVYPDKKYTEKKYEAYPTKQSYPVYPEKKYPEKPYPGYQDYWGQ

Dpfp5 = DPMN_094983, DPMN_159966, or DPMN_159965

MFSTVTIVLLVSGCATATISQYNYWPGGKGLYNNYWNRPQQSYPTWRLYDPCDKVYCYPIYCRYGQYTPQGECCPQCTPGSYRPGSWNNVGQQGNAVSGLGNNVGSQGNAVSGGWNNVGSQGNSVSGGWNNVGSQGNSVSGGWNNVGSQGNAVSGGWNNVGSQGNSVSGGWNNVGSQGNSVSGGWNHVGSQGNAVSG

MSSATKKNTLGKVITLASKGMMWTGMRISLAGKVMPWAGREMTLVSKRMPLVGQGIPWDGKGITWAGKLLEVQIKWIPFGNDTYLT

MIVLEDMPDMIVLEDMSDMIVLEDMPDMIAVEDMPDMMVEDNEEAVVVVVAAVVVVVGLVSVASCCTDLSVV

Dpfp6 = Identified unannotated exons corresponding to this protein

Dpfp7 = DPMN_059800, DPMN_059781, and DPMN_090834. But significant hits to several others

MFSTVTLVLLVSCCGAAFSSWSPYWNSYLPGQGSGKGGYWNSNVPKYGSYWPQQYPSYSGSYWPGWGNNVGSQGNSVRGYGNAVGSQGNDVSGYGNDVGWQWNSVDGKGNYVGSQWNSVN

MFSTVTLVLVVSCFGAALGTYKPYWNSYLPVQGAGNGGYWNSYVPQYGNYGPQKYPGSYWPGAWGGWQGDNVGSQKNSVDGTGNYVGWQKNYVN

MFSTVTLVLFMSCCGVALSSPYWNSYWPGKGSGKDGYWNSYVPQYGNYWPQQYPGYPGGGWQGDNVGSQSNSVDGTGNYVGWQKNYVN

Dpfp8 = DPMN_074797 or DPMN_074964

MVNITNINTQITNIVVNQTSIQNNQLTIMNTLNVVETRVNNTEINVVQITQTTTNIQVQVTEITTEITNIYLNINNVTNQVTNVQTTLVNVTQQQISNTNILNNMTVQVTNIDIQLNQVNVNVNNISVFVNQVSVTVINIETELNIVMSNVTQIEHKIISVGATTEELQRHEDIMIENFMLLSVNITNINIAVFNLTNVVNVVDLNVKGTTGEINGIQKNMNQINIYIKDLDSDINDVSGNVFNIQNNINMIMVNITDIDVRIGNVGGGDQGSLDRIFGSFGKFEERITVITSNQNGITDRLGQIDVRINNTQTGIKEISMKVHDIENDMNSVSINTHQLGVNLNTVSTKITNVETNVNLIMVNITNIDIKVGRIGSDQVTTVKTQDSITKNIQVLVDRQSLIQTSQGKMFDRLGYMDNRLNSTQVNLNIMSGSLTHMEGTIDSLYRNVSKINGTVGQVGSGQIDLQYIQELLSHKLDALEGDQTKAFNSQQEIRVQIKNVMEAVNGIVTTISSKNGSCVGGLNFNDFIGKYATKGLQEDLEKKLDYMSAEQRKSQTVQDHMSVRLDNVLKVLGGVATGNKYSSDEIATLVGSTGGGSVNTGGYSKGTYPVPYGTGGVSGYKSGGR

MVNITNIDIRLGSFGGNISSLGSRNQNIFDILNKFGSVIDLTQTSVTNITRLVSGIGANVDNLTIVVDTLDVNFNSMINNVNVIQNNLNNVSGNVVNVINTVNDIDMTVNNIGGDLVNVTSIVNNIQVNIVSITNNINDIDINIGNLNSDQSVLAQNQNTFNSTVNNLLTVVNNAQTNINSVSQNVQDIGVNFNGLSTVVTNINRDLTGLSSVVNNIQVVQNNIVVDVDNVVNSISNVTVVVNNIQNNVQNIIVNITNIDVTMNSIVVNQTVIQNNQGNIFGVLNGLGNQLTTTTGNVNSLRDLVFQIDINVTSISGDVTNIGGRVSKLTVDIDNIIVNINSVSVNVNNIEGDLNLLFVNITNINNAVGIISSNNSNLLSLTDILSNRLTNVTGQLASASSDIVIISQTVQTVQLNVNNLNNNVNGINGNLNNLTFVVNDIGVTQNTMDGKFNTFVQNINSVTNIANNIQNNVNNMVVNITNIDSTSGAVVVNQTNLQNNQLNIVNLLGSIGGRLTNVDSNINIINLNLTNIQGNIVDLSGGFGNLRADVNVISGDVNNVVSNLQNVNGIMNNIESDLYLMLVNLTDIDTRIGGLGTDQNALQQNINTIGNFLQTFGNRLNVSDMSIFNLSQTVIAISQNVNQINVNLQNVEGDVNVLTNNINNIVGDLGGVSGDVDNLTINVNAVINQVTNIANNVNVMMVNITSINSNVGDIIVNQTQIQQAQNQFGGALNIVNQNVNNVSNAINVISTTMNNIQININNITNNVFNVEGNVNDLTVVVSNIQVNQNNLVTGFNRFGTNVDNLSVTVNAIQSDIGTLIVNVTNIDVTVGSILFNQTNLQNNQVNIVNVVSGINNQVNNTVGQLNFLNNAFTNFNANITGITSIINRQGGDISSLIVQVNQIEVNVNSVSVTVNNIDNDVNIVMMNVTNIGNAIGVLNVNQTTLQQNTQAIANTITNIQNQINAVNSNVVNVVNTITDIDVNVNTVTNNVNNIQVDLNNITNIVNNIQVIQNNMTVAFNDFGSDLNSLSITVNNVINDIDIVMVNITNIGSSMSTVMADQTGIQNNQMNIVNLVNNLGGQLNLTGGNLGSLTQLVNDINGVVNQITVNVNNINGDIGNLTVVVNNVQNTVKNYNVAITNIQNDIDIVMVNVTTINSAVGVFDNDLRSVFGGQTSIFGVLNFLRGQINATNVNINVIVNDIDVIQQNIMNVTFDVANVGNNVNSVSNQVVNIVNNVNGLSTNFGNFQENFNIAVVNITNINQQVNTIGTQINNIESNMNLVMVNITNIGVIINNINSNQTILFDNDANIFNNLNTIGNQVNVTSQSVVNINQIVSNIGGDLNSLSVTVNNVQQNINRVNGVVNSIENDIVNINNNFNIVNSNLNNVSLQVNNIVVDMDVIMVNVTNINGNIRNVIDNQTFMLQGLETTNVNIGNIINVIGVTQNNVNFINNTVTAIGVNVNSINSEIEVINVNVNSLSMNVNNVINNINVIQNDVTKINNTIEQISVSVNTLNVNVNAITVQVTNIEVDLNFVMVNVTNIGSSIRELTVNQTSIVQTQQNIVVALTNIENRVNVTQVSVVKITQTINDIDVTINDISVEVNDIDNSVTNINTQISNIQVNFANLTTVVNTVVNDIDIINVNINNIQVTINTIMVNITDINVNIATIANNQTTIENTQVDIYNQLNNLTNIVQTNSNNIITINQQITEIAVNVTNVAVSVKNIAVEVNNINVQVVNINMTLVEATTNILNIQQNMINVTNIVNDIDVNVNVVMVNITSINTQITNILVNQTSIQNNQLTIMNTLSVVETRVNNTEINVVQITQTTTNIQVQVTDITTKITNIYLNINNVTNQVTNVQTTLVSVTQQQISNTEILNNMTVQVTDIDIRLNQVNVMVNNISVYVSQVSVTVVNIETELNVVMNNVTQIEHKIISVGATTEELQRHENVMIENFMLLSANITNINIAVFNLTNVVNVVDLNVKGTTGEINGIQKNMNQINIYIKDLDSDINDVSGNVFNIQNNINMIMVNITDIDVRIGNVGSGDQGSLDRIFGSFGKFEERITVITSNQNGITDRLGQIDVRINNTQTGIKEISMKVHDIENDMNSVSINTHQLGINLNTVSTKVANVETNVNLIMVNITNIDIKVGRIGSDQVTTVKTQDSITKNIQVLVDRQSLIQTSQGKMFDRLGYMDNRLNSTQVNLNIMSGSLTHMEGTIDSLFRNISNINGTVGQVGSGQIDLQYIQELLSHKLDALEGDQTKAFNYQQEIRLQIKNVMEAVNGIVTTISSKNGSCVGGINFNDFIGKYATKGLQEDLEKKLDYMSAEQRKSQTVQDQMSVRLDNVLKVLGGVATGNKYSSDEIATLVGSTAAGSVNNGGYGKGTLPVGYGTGGGSGNKFGGR

Dpfp9 = DPMN_129085 and DPMN_129263 But significant hits to several others (see spreadsheet and results file)

MLRILRATTKKRSLKMNIKQLMCLLVAAVALLAIAPVANAQYYDYGYGGNNYGYPGNYGYGGNYGGYPGNYGDYDNYGGGWLYKILGGGGKGKGKWGGYGGYGK

MNTKQLMCVLYAAVVLLAVANAQYYDYGYGGNNYGYPGNYGYGGNYGGYPGNYGDYDNYGGGWLYKILGGGGKGKGKWGGYGGYGK

Dpfp10 = DPMN_159708 and DPMN_159905 But significant hits to several others (see spreadsheet and results file)

MLSAVSFLLLVTLYVTVSSQTYKGYPPPKPYPKDPCYKVYCPPIYCPKGQYTPPGECCPRCKKGYGYQDPDPYFPGGK

MLSAVSFLLLVTLYVTVSSQTYKGYNPPKPYPKDPCYKVYCPPIYCPKGQYTPPGECCPRCKKGYGYQDPDPYFPGGK

Dpfp11 = DPMN_141683 and DPMN_141010

MLSAVTLLLLVSCCGMALGQWGGDLCRPVYPPWNCIAVLCAPATNCRYGSFTPKGHCCSVCIGIEINTLFGCIFTDSTDII

MLSAVTLLLLVSCCGMALGQWGGDLCRPVYPPWNCIAVLCAPATNCRYGSFTPYGKCCSVCIGIEINTLFGCIFTDSTDII
